# Structure of the *Thomasclavelia ramosa* immunoglobulin A protease reveals a modular and minimizable architecture distinct from other immunoglobulin A proteases

**DOI:** 10.1101/2024.12.31.630911

**Authors:** Norman Tran, Aaron Frenette, Todd Holyoak

## Abstract

Immunoglobulin A proteases (IgAPs) are a diverse group of enzymes secreted from bacteria that inhabit human mucosal tissues. These enzymes have convergently evolved to cleave human immunoglobulin A as a means of modulating and evading host immunity. Only two of three known IgAP families have been biochemically characterized beyond their initial discovery. Here, we show using solution-scattering, steady-state kinetic, and crystallographic approaches that the protease from *Thomasclavelia ramosa*, representing the uncharacterized third family, has a truly modular and minimizable protein architecture. This analysis also revealed a unique metal-associated domain that likely functions as a molecular spacer and generated a working hypothesis detailing the structural mechanism behind the enzyme’s high substrate specificity. Our work provides the first in-depth biochemical account of this IgAP family, paving the way for advancing clinically relevant IgAP-related research and our understanding of IgAPs as a whole.

## Introduction

Immunoglobulin A proteases (IgAPs) were first described in 1973 as an enzyme active in fecal matter capable of cleaving human immunoglobulin A1 (IgA1) into its Fab and Fc domains (Fig. 1a).^1,2^ By physically decoupling the domains responsible for detecting and responding to antigens, IgAPs are able to reduce the effectiveness of adaptive immune responses in order to facilitate infection.^1^ IgAPs are hence often classified as virulence factors, essential proteins involved in bacterial pathogenesis,^3^ as most strains and species that produce IgAPs are strongly correlated with being pathogenic.^4–6^ Once IgA1 has been proteolyzed, free Fabs can also bind to epitopes to shield the pathogen from opsonization as another mechanism of immune evasion.^7^

**Figure 1.**
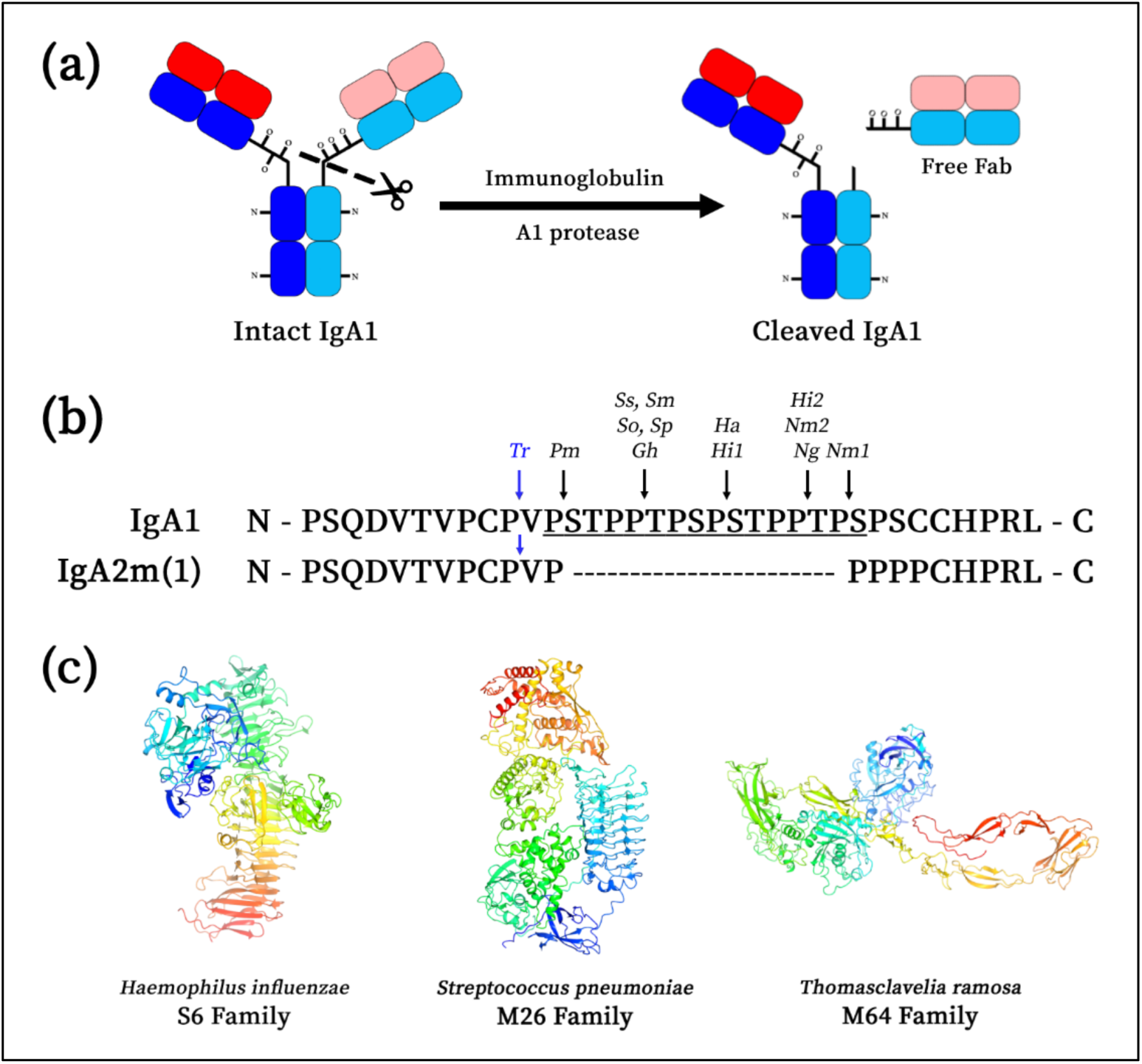
IgA and Its Proteases’ Cleavage Sites and Structures. **(a)** IgA is cleaved at its hinge by IgAPs to separate the Fab and Fc domains as a mechanism of immune modulation. Although only one hinge is shown to be cleaved for simplicity, IgAPs are capable of cleaving both hinges. **(b)** Most IgAPs cleave IgA1 in its elongated hinge (underlined). The protease from *Thomasclavelia ramosa* (*Tr*; blue) is the only IgAP known to cleave both IgA1 and IgA2 allotype A2m(1). The IgAPs from *Prevotella melaninogenicus* (*Pm*), *Streptococcus sanguinis* (*Ss*), *S. mitis* (*Sm*), *S. oralis* (*So*), *S. pneumoniae* (*Sp*), *Gemella haemolyans* (*Gh*), *Haemophilus influenzae* (*Hi*), *H. aegyptius* (*Ha*), *Neisseria meningitidis* (*Nm*), and *N. gonorrhoeae* (*Ng*) only cleave the IgA1 hinge. There are also two IgAP isozymes that can be found in certain serotypes of *H. influenzae* and *N. meningitidis* that cleave at different locations along the hinge. Protein sequences for IgA1 are from UniProt Accession P01876 and IgA2m(1) from NCBI GenBank Gene 3494. **(c)** IgAPs fall under three known enzyme families based on their MEROPS database assignment. One representative member from each family is shown as a ribbon structure and colored as a rainbow from its N-(blue) to C-terminus (red). These members include the serine-type *H. influenzae* Type 1 protease (S6; PDB 3H09), *S. pneumoniae* metalloprotease (M26; PDB 6XJB), and *T. ramosa* metalloprotease (M64; AlphaFold2 model).

Many bacterial species that produce IgAPs are heavily associated with lower respiratory tract infections (LRTIs),^8–10^ one of the leading causes of death for all ages worldwide.^11^ Because of IgAPs’ role in the bacterial virulence of LRTI-associated pathogens,^1^ there have been efforts to develop IgAP inhibitors to treat LRTIs as an alternative to traditional antibiotics.^12–17^ IgAPs have also been of interest in human disease research for their ability to discern between IgA and other structurally similar substrates.^18–23^ This makes them good potential therapeutics to treat diseases associated with aberrant IgA^24^ as their high substrate specificity may lower the risk of off-target effects relative to small-molecule drugs.^25^ One of these diseases is immunoglobulin A nephropathy (IgAN), a chronic and debilitating autoimmune disease characterized by the buildup of IgA1 aggregates in the kidney.^26^ IgAPs have shown to be a promising IgAN therapeutic through various *in vivo*,^27^ *ex vivo*,^28^ and *in vitro* experiments.^29^ Regardless of whether IgAPs are to be studied for their role in pathogenicity, as a drug target, or as a protease-based therapeutic for IgAN, an in-depth biochemical analysis of IgAP structure and function is needed to advance these biological and clinical endeavors.

A plethora of IgAPs have since been identified from bacteria that inhabit other mucosal tissues such as the upper respiratory tract,^8^ gastrointestinal system,^30^ urogenital tract,^31^ and oral cavity.^32^ These studies showed that all IgAPs cleave the IgA hinge C-terminal to a proline residue (Fig. 1b).^21,23,32–35^ Although these IgAPs were historically categorized based on their catalytic mechanism,^1,2,18^ the advent of genome sequencing and, more recently, language-model-driven structural prediction revealed that these initial categories more realistically represented enzyme families (Fig. 1c) – a set of homologous enzymes that have similar protein sequences, biochemical characteristics, and structures.^36,37^ Despite IgAPs having been described as serine, cysteine, and metal-dependent proteases, only serine and metal-dependent IgAPs have been bioinformatically annotated, assigned enzyme families, and structurally characterized.^20,38,39^

Of the three known IgAP families (Fig. 1c), only the S6 and M26 IgAPs have had their structures experimentally determined.^19,20,39^ The last remaining metal-dependent M64 family, as represented by the IgAP from *Thomasclavelia ramosa*,^21^ previously known as *Clostridium ramosum*,^40^ is the least understood; its structure has only been computationally modeled and sparsely characterized beyond its initial discovery.^22,41^ Despite this, the *T. ramosa* IgAP has several properties that make it an untapped system to study. Unlike the majority of IgAPs which can only cleave IgA1 (Fig. 1b),^1^ the *T. ramosa* IgAP can cleave IgA1 and IgA2, specifically IgA2 allotype A2m(1).^21,35^ The enzyme is also predicted to have a modular domain architecture, contrary to the IgAPs from *Haemophilus influenzae* (S6) and *Streptococcus pneumoniae* (M26) (Fig. 1c), which cannot be substantially reduced in size without abolishing function.^20,38,39^ These large structural and functional differences raise several interesting questions: Is the enzyme truly modular and can it be substantially minimized in contrast to other IgAPs? Are the non-proteolytic domains involved in catalysis and, if not, what are their functions? And, what are the structural mechanisms underlying its substrate specificity?

This work will address these questions through in-depth biochemical and structural analysis of the *T. ramosa* IgAP. This will be accomplished through three complementary and intertwined experimental approaches: (1) *Small-angle X-ray scattering (SAXS)*. SAXS was used to validate the modularity of the enzyme architecture and determine the overall conformation of the full-length enzyme to ascribe functions to each domain. (2) *Steady-state kinetics*. In the event that the enzyme is truly modular and minimizable, the activity and substrate specificity of these minimized constructs should be assessed to ensure they retain their function. A gel-based kinetic assay was developed to obtain Michaelis-Menten parameters for meaningful comparisons. And (3), *Crystallography.* The crystal structure of the proteolytically active domain was solved to generate hypotheses about how the protease functions, why the enzyme is so specific, and what structures are important for catalysis.

## Results and Discussion

### AlphaFold2 Suggests That the T. ramosa IgAP Has a Modular Domain Architecture

The *T. ramosa* IgAP AlphaFold2 model was used as the basis for splitting the protease into putative domains (Fig. 2a).^42,43^ This model was generated using the entire secreted protease sequence (UniProt accession #Q9AES2, residues 31-1195), which was previously shown to be active when recombinantly expressed in *Escherichia coli*.^41^ The domain bioinformatically predicted to be an M64 protease was arbitrarily assigned to be the middle of the protein (middle domain; MD) and the other domains were given names relative to the MD (Fig. 2b). This yielded six putative domains which were recombinantly expressed and biochemically characterized in isolation or as multi-domain constructs (*Suppl.* Fig. 1). The boundaries of these domains were initially set based on AlphaFold2’s PAE metric^42^ and subsequently manually adjusted based on the model’s predicted enthalpic interactions.

**Figure 2.**
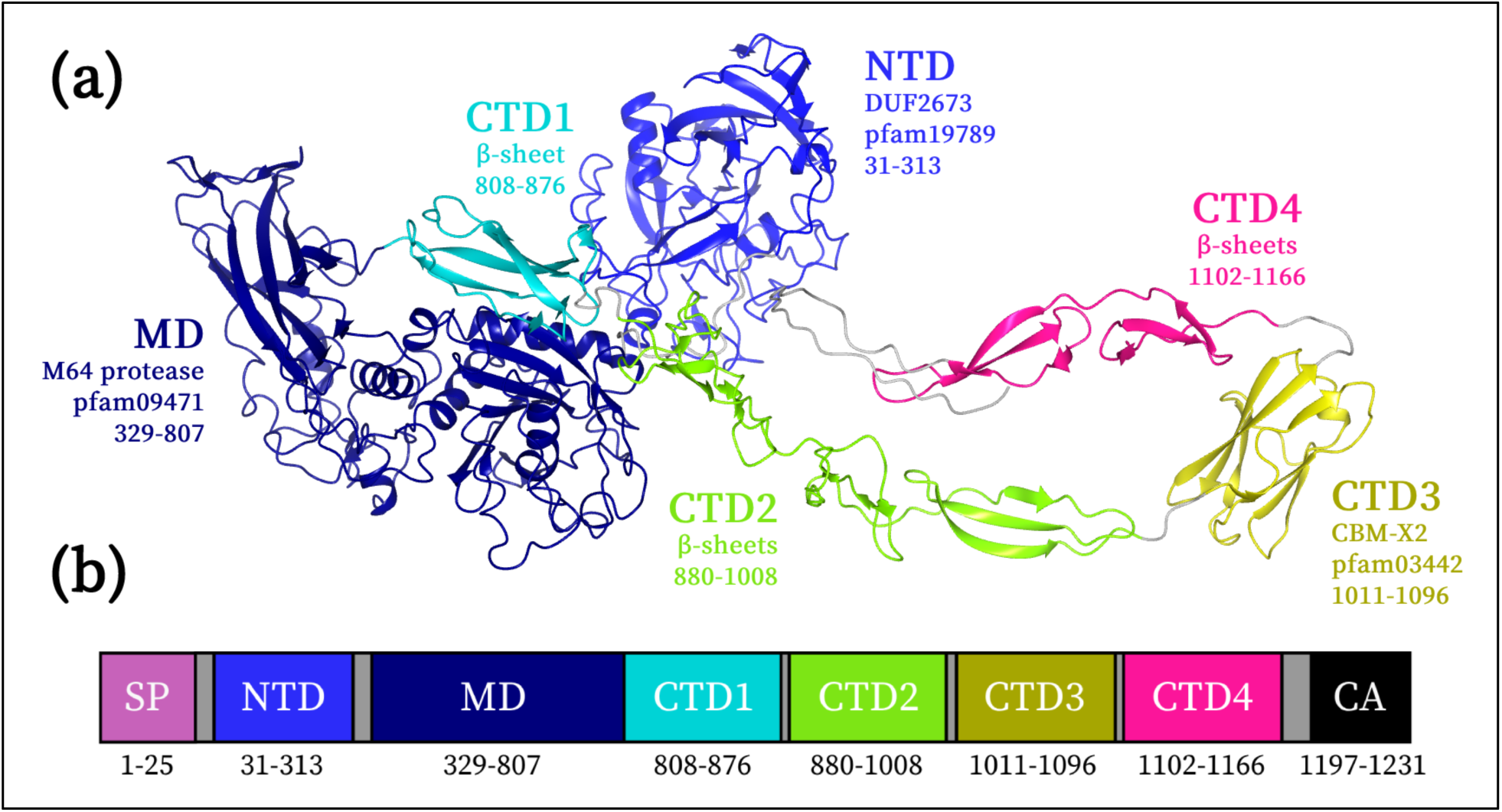
Putative Domains of the *T. ramosa* IgAP. **(a)** The full-length AlphaFold2 model was split into six putative domains joined together by flexible linkers (grey): N-terminal domain (NTD; blue), middle or protease domain (MD; dark blue), C-terminal domain #1 (CTD1; cyan), C-terminal domain #2 (CTD2; green), C-terminal domain #3 (CTD3; yellow), and C-terminal domain #4 (CTD4; pink). The domains’ bioinformatic annotations, if any, along with their Pfam accession and residue numbers are listed accordingly. **(b)** These domains were named for their position relative to the proteolytically active MD. The immature protein is shown schematically and contains a signal peptide (SP) and cell anchor (CA) which are removed during secretion.

There were a few additional factors that needed to be considered when determining these domain boundaries: (1) although CTD2 and CTD4 at first appear to be comprised of several separate β-sheets, key cysteine and histidine residues are predicted to form zinc-binding pockets that link the sheets (*Suppl.* Fig. 2).^44^ These sheets are therefore hypothesized to come together as a single structural unit; and (2) despite the AlphaFold2 PAE metric^42^ and PISA analysis^45^ suggesting that the MD and CTD1 form a domain together, the AlphaFold2 model suggests that the interface is relatively hydrophilic while RosettaFold predicts CTD1 as its own separate domain (*Suppl.* Fig. 3).^46^ As such, constructs for the MD with and without CTD1 were made to further assess the modularity and functional necessities of the protease domain.

### The Full-Length Protease Takes on an Extended Rod-Like Conformation in Solution

Small-angle X-ray scattering (SAXS) was used to reveal how individual domains behaved in and outside of the context of the full-length enzyme. This method was particularly powerful in understanding this enzyme as there are no restrictions on the protein sample, meaning that measurements can be done on inherently flexible multi-domain constructs (*Suppl.* Fig. 1ac).^47^ Programs such as FoXS and CORAL were used to better account for inter-domain flexibility to validate and optimize these predicted multi-domain models.^48,49^ SAXS can thus test the hypothesis of whether these proteins are domains in the strictest sense (*i.e.*, part of a protein that retains both its structure and function in isolation)^50^ and reveal what other biological roles they might have in the absence of more difficult-to-obtain experimental information. SAXS data parameters and statistics for all constructs are summarized in *Suppl. Table 1* and their scattering curves and Guinier plots are shown in *Suppl.* Fig. 4.

The SAXS data for the full-length enzyme was first analyzed to determine whether AlphaFold2 accurately captured the enzyme’s solution conformation (Fig. 3a). It came as a surprise to find that the simulated scatter of this predicted model (Fig. 2a) did not fit the observed SAXS data at low scattering angles (Fig. 3b), suggesting that this model had an overall conformation notably different to its solution structure. To derive a more representative model, the AlphaFold2 model was relaxed using simulated annealing (CORAL program) under the assumption that the linkers between the assigned domains (Fig. 2, grey) were flexible.^48^ This solution structure (Fig. 3c) suggests that the full-length enzyme takes on an extended conformation more consistent with the particle’s relatively large R_g_ and D_max_ values. This shape primarily results from the rod-like structure of the C-terminal tail (CTD1-4), which is also seen in the CORAL-relaxed solution structures of the CTD1-4 and MD+CTD1-4 proteins (*Suppl.* Fig. 5a-f). The normalized Kratky and P(r) plots of these constructs indicate that they are well folded and retain their elongated conformation when produced in isolation (Fig. 3ef). In contrast to the C-terminal tail, the position of the NTD relative to the MD did not significantly differ between the predicted and relaxed solution structures for the full-length (Fig. 3c) and NTD+MD proteins (*Suppl.* Fig. 5g*-i*), despite the NTD-MD linker being by far the longest inter-domain linker in the whole protease (NTD-MD linker: 15 residues; other linkers: 3-5 residues). The elongated conformation of the full-length enzyme is thus primarily due to the C-terminal tail.

**Figure 3.**
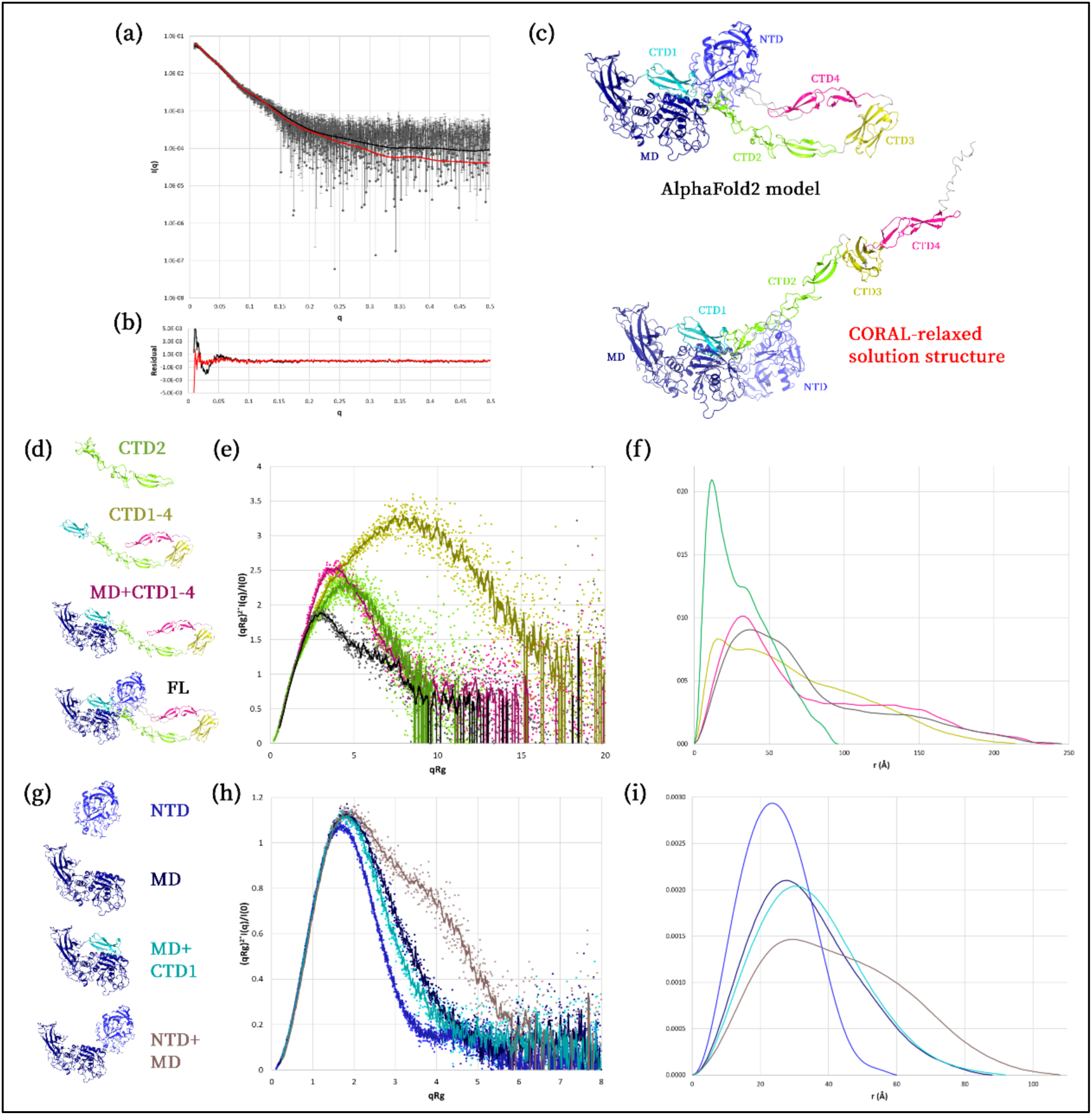
SAXS Analysis of *T. ramosa* IgAP Constructs. **(a)** The scattering curve for the full-length enzyme is presented as the mean intensity ± one standard deviation for all measured angles. FoXS was used to generate the simulated solution-scattering intensities of the AlphaFold2 (black line) and CORAL-relaxed (red line) full-length models, which are overlayed onto the scattering curve. **(b)** These simulated scattering curves were compared to the full-length SAXS data and residuals were calculated for each angle. The CORAL-relaxed model had a better data-to-model fit relative to the AlphaFold2 model, with FoXS χ^2^ values of the CORAL-relaxed and AlphaFold2 fits being 0.65 and 0.93, respectively. **(c)** The CORAL-relaxed model showed that the full-length enzyme had an extended C-terminal tail, which was not predicted in the AlphaFold2 model. Constructs that were predicted to be **(d)** elongated and **(g)** globular based on their AlphaFold2 model were analyzed with **(e, h)** normalized Kratky and **(f, i)** P(r) plots. The color scheme for each construct corresponds to those in **(d)** and **(g)**.

### CTD1 and CTD3 Are Structurally Homologous to Known Periscope Domains

PDBeFold^51^ was used to find structural homologues of the CTDs using their AlphaFold2 models to understand their putative function in the context of the C-terminal tail. CTD1 is structurally homologous to a domain in the *Listeria monocytogenes* InlB protein (*Suppl.* Fig. 6a), a virulence factor involved in host-cell entry.^52^ This domain, known as the B-repeat, has a β-grasp ubiquitin-like fold and was initially thought to play a purely structural role as a molecular spacer.^52^ The B-repeat has since been shown to have an additional receptor-binding function that facilitates pathogenicity.^53^ CTD3 is structurally homologous to a domain in human titin (*Suppl.* Fig. 6b), a structural protein involved in muscle contraction.^54^ This domain has an immunoglobulin I-type fold and is one of the many tandem repeats that gives titin its rigid and elongated structure.^54^ Both β-grasp ubiquitin-like and immunoglobulin-like folds (*i.e.*, β-sandwiches including the I-type fold) are well-characterized domains known to exist as tandem repeats that assist in the spacing or projection of other functional domains away from the surface of the cell.^55^ This projection allows these proteins to sense and influence a wider extracellular environment, enabling them to mediate cell-cell contacts and biofilm formation.^55^ These so-called periscope domains can also bind to small molecules^56–58^ and form larger oligomeric structures,^59^ amongst many other functions (*e.g.*, InlB B-repeat).^52,55,60^ Due to the large diversity within these periscope domains,^55^ it is hard to prescribe function solely based on these factors alone. However, it is clear that their structural homology with CTD1 and CTD3 strongly suggest that the function of these domains is to contribute to the C-terminal tail’s extended conformation.

### CTD2 and CTD4 Represent an Uncharacterized Fold of Metal-Associated Periscope Domains

PDBeFold^51^ was unable to find a reliable (< 2.5Å Cα-RMSD) structural homologue of CTD2 or CTD4. As CTD2 and CTD4 are predicted to be structurally homologous to each other, SAXS data for CTD2 were analyzed to ensure that this lack of homology was not due to an inaccurate search model. CTD2 and CTD4 were previously hypothesized to contain zinc-binding sites that bridge the gap between individual β-sheets within each domain (*Suppl.* Fig. 2). When CTD2 was produced in isolation, its SAXS data indicated that the protein was well folded (Fig. 3e) but had strikingly large R_g_ and D_max_ values despite its low molecular weight. It turns out that the AlphaFold2 model of CTD2, although it does not take metal into account during modeling, is very consistent with the solution-scattering data (*Suppl.* Fig. 7). In other words, these large biophysical values are only possible if CTD2 exists as a fully extended rod (Fig. 3f) with the sheets placed in an end-to-end conformation.

Although CTD2 and CTD4 seem to be unique in their fold and intra-domain organization, they share features with periscope domains from the *Staphylococcus aureus* SasG and *S. epidermidis* Aap proteins. For one, the repeated G5 domain from SasG and Aap have similar secondary-structure characteristics to CTD2 and CTD4, in that they are both enriched with single β-sheets that do not form β-sandwiches.^59^ Secondly, with the exception to one of these sheets (*Suppl.* Fig. 2, sheet #1), CTD2 and CTD4 lack a defined hydrophobic core, drawing similarities to the G5 domains from Aap.^59^ Moreover, surface-bound SasG and Aap can interact with the same proteins on other cells in a metal-dependent manner to facilitate cell-cell interactions, a key phase in biofilm formation.^59^ Aap dimerization is mediated by zinc-binding motifs that are found on the surfaces of each protomer, which assemble the two protomers in an antiparallel fashion.^59^ Despite SasG and Aap’s ability to be functionally regulated by zinc, it remains unclear whether this holds true for CTD2 and CTD4. There are, however, structural differences that may provide additional insights. The zinc-binding motifs in Aap are atypical due to the lack of cysteines and presence of acidic residues in their binding motifs, resulting in pliable metal-coordination sites that can accommodate metals other than zinc.^59^ Additionally, the zinc ions bound to one protomer are partially coordinated with water molecules to allow for dimerization to occur.^59,61^ In stark contrast, the predicted zinc-binding sites for CTD2 and CTD4 are solely composed of cysteine and histidine residues (*Suppl.* Fig. 2). These residues are predicted to fully coordinate the zinc ions, leaving no additional coordination sites for oligomerization. These zinc-binding motifs are also well-documented for their role in the structural stabilization of DNA-binding proteins.^44^ This suggests that the role of zinc in these domains is purely structural, however, whether or not CTD2 and CTD4 have additional functions still remains an interesting future line of questioning. Thus, we propose that CTD2 and CTD4 represent an uncharacterized periscope domain fold whose function perhaps relies on and is regulated by zinc in the bacterium’s environment.

There is strong evidence, through structural and functional homology, to suggest that all of the CTDs are periscope domains. Although the C-terminal tail does not contain identical repeats, as seen in canonical Periscope Proteins,^55^ its elongated shape and modular domain architecture is nevertheless reminiscent of these canonical structures; the CTDs are thus periscope domains that are repeated with respect to their function, not their structure. The combined function of the CTDs is thus to project the proteolytically active M64 middle domain away from the cell surface to augment its immune-evasion function.

### The MD Retains Its Fold and Shape When Produced in Isolation of the Full-Length Enzyme

It was previously hypothesized that CTD1 is a separate domain from MD. As previously discussed, the SAXS data for CTD1-4 suggested that CTD1 can fold independent of the MD when functioning in the context of the C-terminal tail (Fig. 3ef). SAXS analysis of the MD and MD+CTD1 proteins likewise finds that the MD can fold without CTD1 (Fig. 3h); and, although there are slight differences in their R_g_values and P(r) distributions, its shape is also retained without CTD1 (Fig. 3i). This supports the hypothesis that MD, not MD+CTD1, is the smallest self-contained protease fold. The NTD, on the other hand, is also well folded when produced in isolation or in the context of the MD (Fig. 3h). SAXS data for the NTD+MD construct reinforces this modular hypothesis as its distinct bimodal P(r) distribution (Fig. 3i) indicates that the NTD and MD act as two separate entities tethered by a flexible linker.^62^ Finally, all of these results are supported by the fact that the AlphaFold2 models of NTD, MD, and MD+CTD1 are very consistent with their solution-scattering data (*Suppl.* Fig. 7), further cementing the idea that all *T. ramosa* IgAP domains retain their fold and structure in isolation of the full-length protein.

One major question remains regarding the modularity of the *T. ramosa* IgAP. Although SAXS has been an immensely powerful tool that has revealed that all of the putative domains are structurally modular, true domains also retain function in addition to fold when produced as a standalone entity.^50^ The CTDs fit this classification as their function is intrinsically tied to their structure. The NTD and MD, however, have yet to be discussed in this context. The function of the NTD is unknown and hence it is classified as a domain of unknown function.^63^ There is some bioinformatic analysis suggesting that these domains may be involved in viral hypermutation,^64^ however experimental evidence for this has been lacking. The classification of the NTD as a true domain, therefore, still remains inconclusive.

More pertinent to the rest of the discussion and study of the protein as a whole is the proteolytic function of the MD. It can be argued that due to the protease’s specificity for IgA,^18–23^ it may be possible that some of the other domains may play a role in substrate association and recognition. In other words, the fact that the MD is the smallest fold that contains the predicted structures necessary for catalysis is not sufficient evidence to say it also retains the complete catalytic functionality of the full-length enzyme. SAXS by itself is thus unable to conclude that the MD is a true domain and a thorough investigation into the kinetic capabilities of all MD-containing constructs must be undertaken.

### Minimizing the Enzyme to the MD Does Not Reduce Protease Activity nor Substrate Specificity

All IgAPs are known to have high substrate specificity as they are able to selectively cleave IgA over other immunoglobulins and are unable to cleave the IgA hinge peptide in isolation.^18–23^ Although the structural mechanism by which this occurs in the *T. ramosa* IgAP has yet to be described, the only substrate that contains all of the structural features required to assess this activity is the native IgA molecule. As no robust and generalizable IgAP assay that uses IgA1, and not small-molecule substrates,^14,65,66^ has been comprehensively described in the literature, we developed a gel-based assay capable of deriving Michaelis-Menten steady-state parameters that could be meaningfully compared between MD variants. A thorough discussion of the assay’s method-development process, assumptions, drawbacks, and results are detailed in *Suppl. Discussion.* Briefly, the substrate and products of the reaction are separated and visualized on an SDS-PAGE gel (*Supp.* Fig. 8). The steady-state initial rates of IgA1 cleavage were determined by measuring the intensity of a product band and quantifying how it changes over time (*Supp.* Fig. 12). This was done for various constructs across a range of IgA1 concentrations to yield several Michaelis-Menten curves (Fig. 4) and their corresponding kinetic parameters (*Suppl. Table 2*).

**Figure 4.**
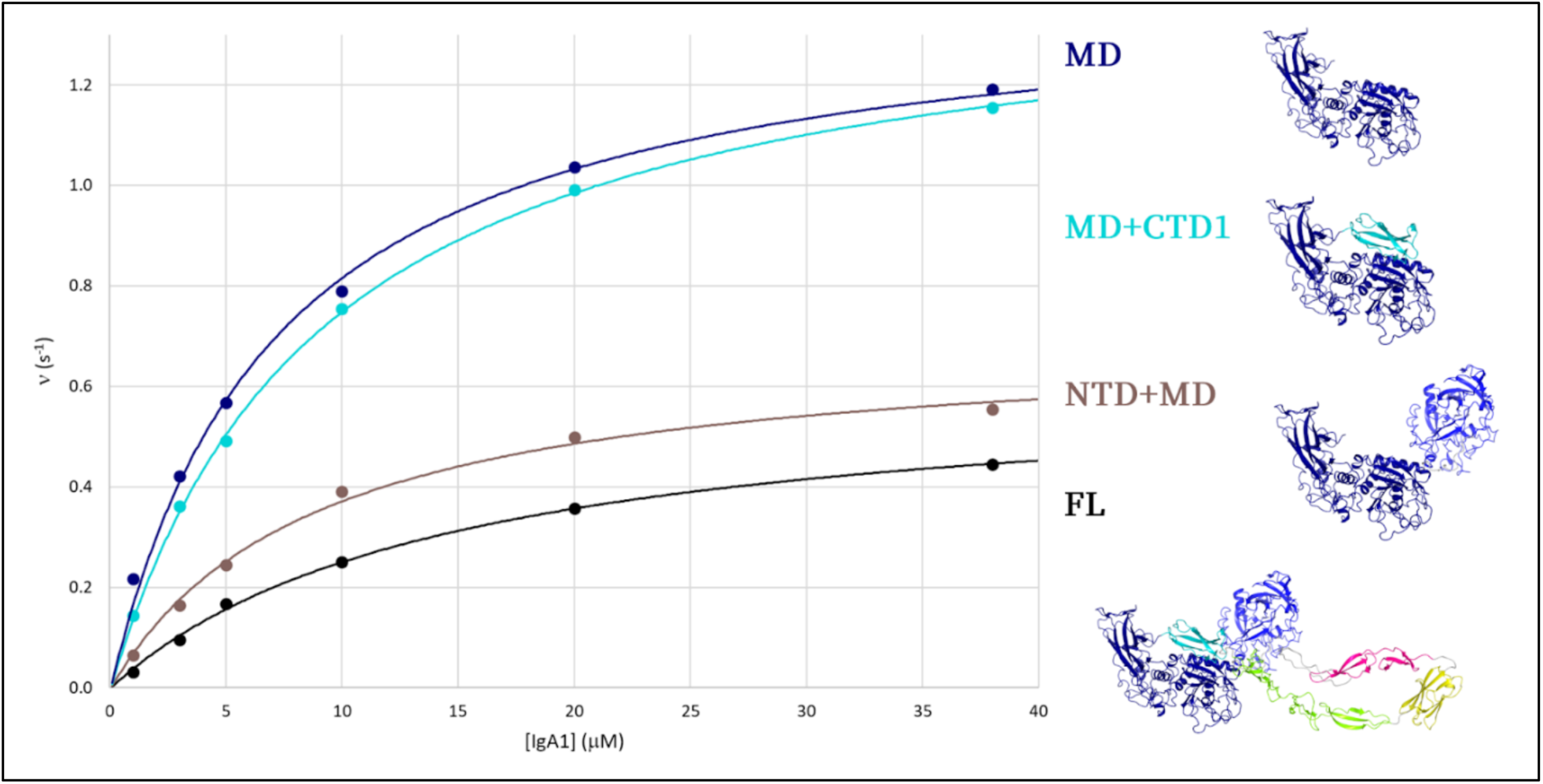
Michaelis-Menten Curves for Various Domain Constructs. The normalized initial rates (ν) for the full-length (black), NTD+MD (brown), MD+CTD1 (cyan), and MD (blue) enzymes were plotted against IgA1 concentration and fit to the Michaelis-Menten equation. The Michaelis-Menten parameters that best fit these data are summarized in *Suppl. Table 2*. The rates of cleavage increase across all substrate concentrations as the enzyme is minimized to the smallest construct (MD).

These kinetic data show a clear increase in both fundamental kinetic parameters (k_cat_ and k_cat_/K_M_)^67^ across all IgA1 concentrations as the full-length enzyme was minimized to the MD. While large increases in these values were seen when the C-terminal tail and NTD were removed, only small changes were seen when CTD1 was truncated. This reinforces the hypothesis that the CTD1 does not have an impact on the structure or function of the MD. Further discussion of these parameters can be found in *Suppl. Discussion*. In support of the SAXS data conclusions, these kinetic data show that the *T. ramosa* IgAP has a truly modular domain architecture as it can be minimized to only the MD (∼40% of the total enzyme size) without negatively impacting proteolytic function. Rather, the truncated MD construct is even more active *in vitro* compared to the full-length construct.

It was also necessary to verify that the minimized MD construct still retained the high substrate specificity characteristic of the full-length enzyme ^22,23^ and that the observed increased kinetic activity was not due to the enzyme having substrate promiscuity. To test this possibility, a synthetic octapeptide, based on the IgA1 hinge cleavage site (Fig. 1b), was incubated with the full-length and MD constructs. MALDI-ToF mass spectrometry was then used to detect the mass of peptides in the reaction mixture following incubation. These data demonstrated that there was no evidence of peptide cleavage by either construct (*Supp.* Fig. 14a-c), whereas, all substrate was cleaved when the reaction was carried out using IgA1 instead of the octapeptide (*Supp.* Fig. 14d).

### Crystal Structure of the Protease Domain Shows Conserved Metzincin Structural Features

Many questions remain regarding the protease function of the *T. ramosa* IgAP and how it is able to retain its activity and substrate specificity upon minimization to just the MD. The high-resolution crystal structure of the MD was solved to gain insight into these phenomena (Fig. 5a). The MD was only able to be crystallized in the context of the MD+CTD1 construct as CTD1 made the protein more globular and crystallizable. As CTD1 is neither needed in the folding nor activity of the MD, as previously discussed, it will be omitted from the following discussions. All crystallographic data collection and refinement statistics are summarized in *Suppl. Table 3*.

**Figure 5.**
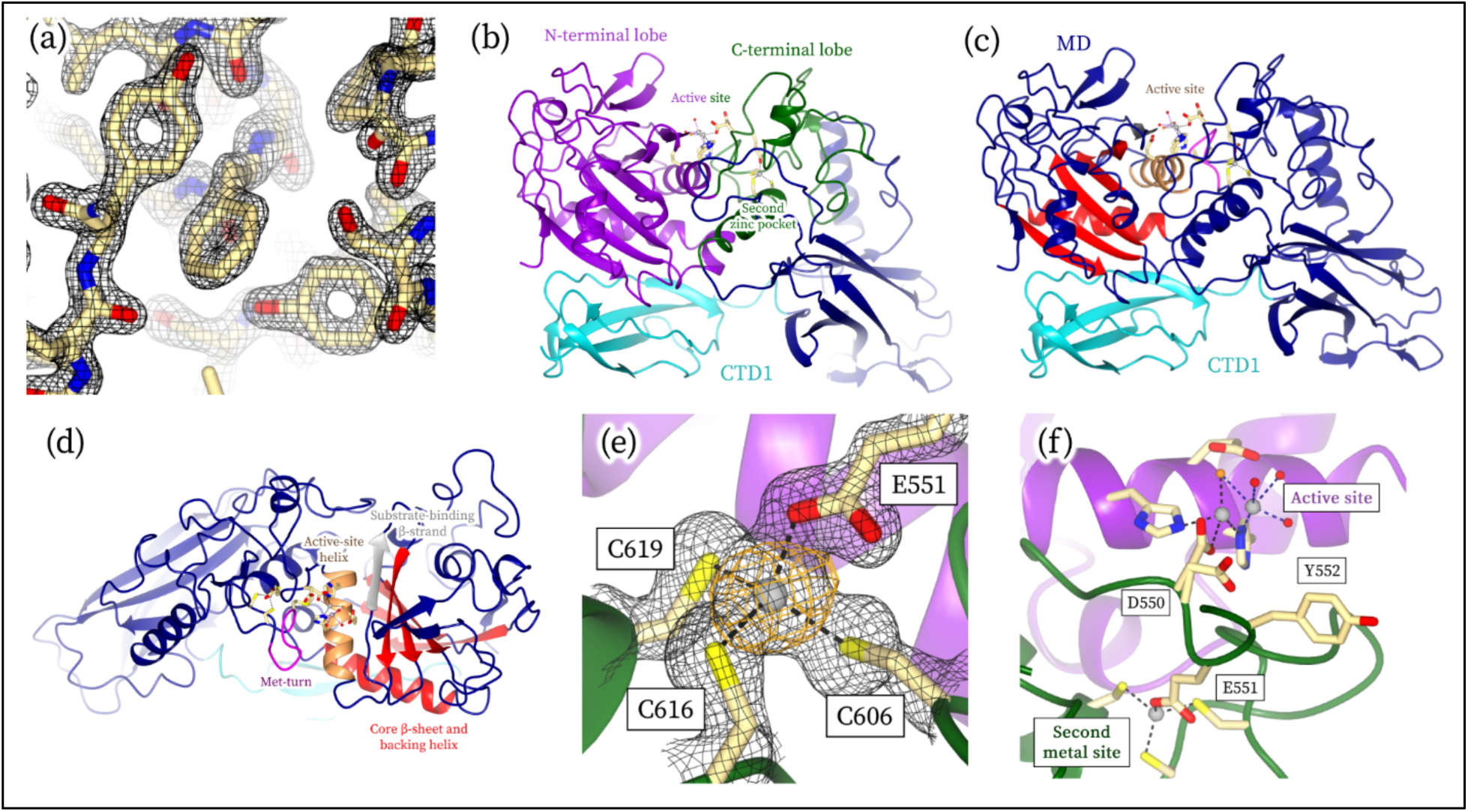
Crystal Structure of the *T. ramosa* IgAP Middle Domain. The crystal structure of the protease domain was solved in the context of the MD+CTD1 construct. An example of the good map-to-model fit is shown in panel **(a)** (black density at 2.5 σ 2Fo-Fc). **(b)** The structure of the zinc-supplemented MD+CTD1 holoenzyme. Residues involved with zinc (grey spheres) binding are shown as cylinders. The active site is found at the interface between the N-(purple) and C-terminal (green) lobes. **(c)** The metzincin fold of the *T. ramosa* IgAP contains: 1) An alpha helix backing a large beta sheet (red). The incoming peptide substrate makes an antiparallel beta-strand interaction with the last strand of this beta sheet adjacent to the active site (grey). 2) An active-site alpha helix (brown) that contains the HEXXH motif. And 3) a loop containing a conserved methionine called the Met-loop (magenta). A top-down view of the active site and metzincin features is shown in **(d)**. **(e)** The C-terminal lobe also contains a second zinc-binding pocket. This zinc is coordinated (black dashes) by the side chains of three cysteines and one glutamic acid (black density at 1.5 σ 2Fo-Fc; anomalous difference map at 5.0 σ in orange) and is linked to the active site by a conserved loop (D550-Y552) that bridges both metal-binding sites **(f)**.

The MD is structurally homologous to a large family of metalloproteases known as metzincins.^68^ Metzincins have several structural features that make up the catalytic fold that are conserved in the MD.^68^ These include an active-site zinc ion that prepares the substrates for the reaction (mechanism summarized in *Suppl.* Fig. 15).^68^ This zinc ion is housed in the peptide-binding cleft, which is formed from two characteristic lobes of the metzincin fold (Fig. 5b).^68^ It is also coordinated by key residues found on an α-helix in the N-terminal lobe (metalloprotease HEXXH motif) and D550 in the C-terminal lobe (Fig. 5c).^68^ The metzincin fold also contains a core beta sheet in the N-terminal lobe and a conserved Met-turn in the C-terminal lobe, which stabilize the incoming peptide and zinc-binding site, respectively (Fig. 5d).^68^ Unlike the majority of metzincins, the MD contains an additional non-catalytic zinc-binding site in the C-terminal lobe (Fig. 5be). This second metal-binding site is adjacent to the active site and is bridged by a loop (D550 to Y552) that is involved in the coordination of both metals (Fig. 5f). It is unclear if this second zinc-binding site has any biological role in addition to structural stabilization. However, given its proximity to the active site, one can speculate that the second site may modulate the affinity of zinc to the active site as removing zinc from the second site likely impacts the dynamics of the bridging loop and thus one of the active-site zinc ligands. The role of the second zinc-binding site therefore remains speculative and more experimental evidence is needed to make any claims on its specific function.

### Conformational Heterogeneity in the Active Site is Hypothesized to Impart Substrate Specificity

Early studies on metalloproteases have shown that all active-site residues and water molecules are well positioned in their catalytically competent state ready for an incoming substrate (*e.g.*, *Suppl.* Fig. 16).^68–71^ The first crystal structure of MD+CTD1 solved, however, depicts a conformationally heterogeneous active site (Fig. 6ab) in contrast to these model metalloproteases. Not only is the active-site metal found in two positions, but one of the zinc-binding residues (D550) is too far to coordinate the metal in either position. This heterogeneity was hypothesized to be due to the enzyme binding to a mixture of metals, as zinc was only added in trace amounts during expression. To resolve this, the purified enzyme was chelated and crystallized in its metal-free state. These crystals were then used as-is or soaked with 1 mM ZnCl_2_ prior to cryoprotection and freezing. The crystal structure of the as-purified enzyme was compared to the metal-chelated and zinc-supplemented structures to look for structural changes at the two metal-binding sites. Datasets were also collected above and below the zinc absorption edge to experimentally confirm the metal identity at both sites.^72^

Anomalous difference maps of the as-purified crystal structure (Fig. 6ci) support the hypothesis that the enzyme was bound to a heterogeneous mixture of transition metals in the active site. The second metal-binding site, on the other hand, is solely selective for zinc as there is no anomalous signal in the below-edge data for any of the structures collected, regardless of their metalated state. When the enzyme was chelated, the active-site metal was replaced with two water molecules that occupy the same positions as the zinc in the as-purified structure (Fig. 6cii). When this metal-chelated protein was treated with 1 mM ZnCl_2_, both sites became fully and unambiguously occupied with zinc (Fig. 6ciii). This zinc-supplemented structure therefore represents the holoenzyme of biological interest and will be the focus of further discussion. The active site of the holoenzyme was still conformationally heterogeneous despite the full occupancy of zinc at the active site (Fig. 6de). The same general features were seen in both as-purified and holoenzyme structures (Fig. 6ab), with the zinc ion split approximately equally between the two positions. However, unlike in the as-purified structure, the zinc that is in the catalytically active position now coordinates with D550 (Fig. 6e). These structures suggest that the conformational heterogeneity seen in the active site is an inherent property of the protein itself and not an artifact due to promiscuous metal binding.

**Figure 6.**
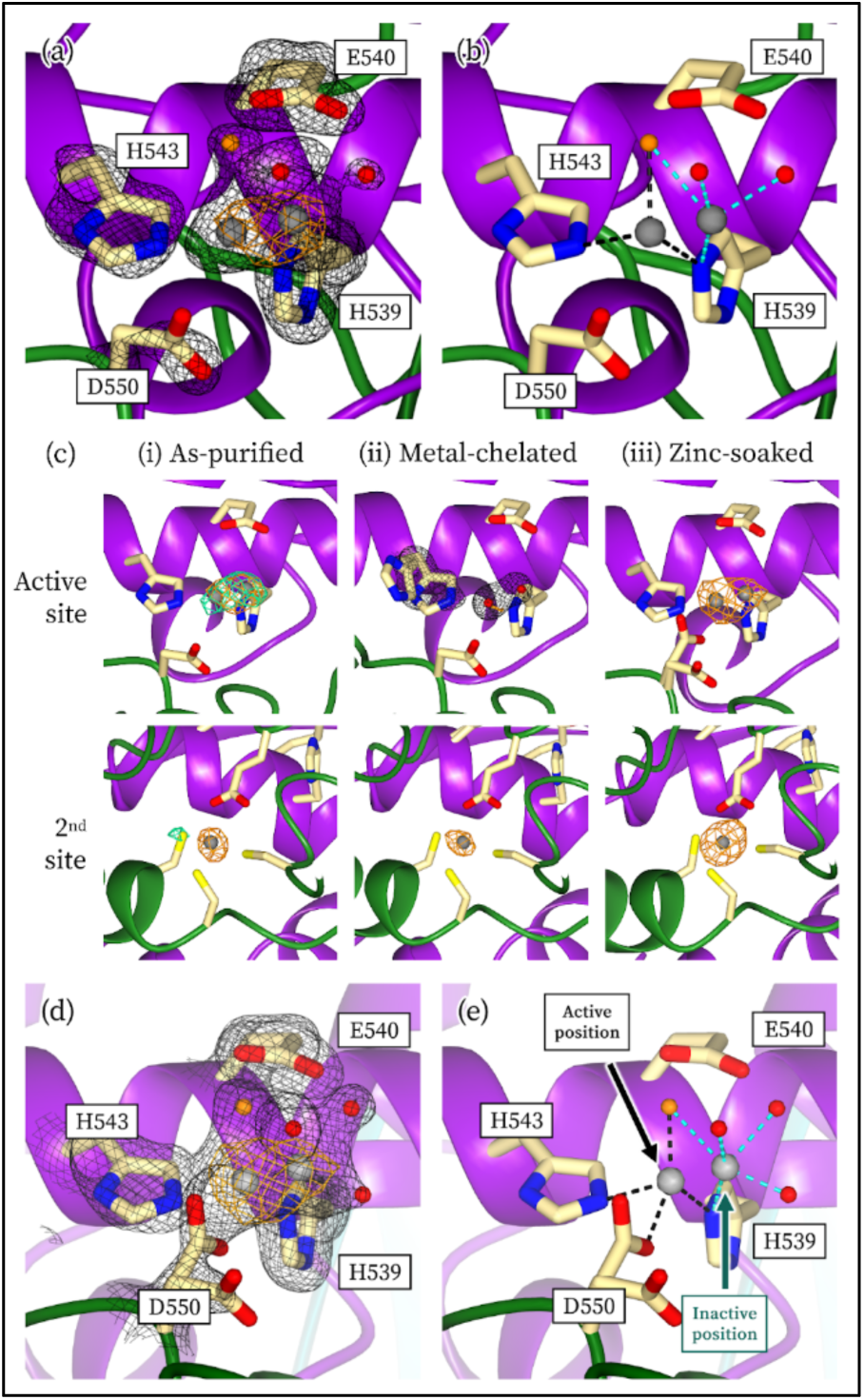
The Active Site of the Protease Domain is Inherently Conformationally Heterogeneous. **(a)** The structure of the as-purified MD+CTD1 enzyme shows that the active-site zinc (grey spheres) can be modelled in two positions (orange anomalous difference density at 4.0 σ, black 2Fo-Fc density at 1.3 σ). The nucleophilic water molecules for all panels are colored in orange. **(b)** These two positions have separate sets of coordinating ligands (black and cyan dashes). **(c)** To ensure that this heterogeneity was not artifactual, metal-chelated and zinc-supplemented structures of the enzyme were solved. The anomalous density maps for the active-(top row) and second sites (bottom row) are shown. Maps for the above-(orange) and below-zinc-absorption edge (green) data are contoured at ∼0.10 e^-^/A^3^ and overlayed over the protease domain structures. *(i)* The as-purified enzyme is bound to a heterogeneous mixture of metals at the active site, a notable portion of which is not zinc (3.9 σ above-edge; 3.8 ο below-edge). The second site is always selective for zinc as no anomalous signal is seen in the below-edge data. *(ii)* The metal-chelated enzyme has no metal at the active site (4.3 σ above-edge; 4.4 σ below-edge) and instead has two water molecules that occupy the same space (black density; 1.3 σ 2Fo-Fc). H543 takes on two conformations due to the lack of active-site metal. *(iii)* Once zinc is soaked back into the crystal, both pockets become saturated with zinc (5.3 σ above-edge; 5.2 σ below-edge) and display strong anomalous densities in the above-edge but not in the below-edge data. **(d)** This zinc-supplemented structure still shows conformational heterogeneity in the active site, suggesting that it is an inherent property of the enzyme (orange anomalous difference density at 5.0 σ, black 2Fo-Fc density at 1.3 σ). **(e)** When zinc is in its active position, it is properly coordinated with all residues and water molecules appropriately positioned for catalysis (black dashes, *Suppl.* Fig. 15). The zinc in its inactive position is coordinated with an alternate set of water molecules (cyan dashes).

This active-site heterogeneity is hypothesized to have significant functional consequences on how the *T. ramosa* IgAP is able to select for its substrate. In the S6 IgAP family, several elongated loops are thought to sterically occlude the entrance to the active site when no IgA1 is bound as a structural mechanism responsible for substrate specificity.^19^ These gating loops would then be pinned back when distal interactions occur between the enzyme and IgA1 to open the active site for proteolysis.^19^M26 IgAP family members have a functionally analogous beta-hairpin structure that has been shown to gate the active site when no IgA1 is bound.^20,39^ In contrast, the *T. ramosa* IgAP protease domain does not have any structural means of sterically occluding the active site to prevent non-specific proteolysis, yet it is somehow able to distinguish between the IgA1 hinge peptide and the whole IgA1 molecule. These structural data, however, suggest the M64 IgAP has an active site that can switch between catalytically competent and inactive conformations in its ground state (Fig. 6e). We propose that this conformational heterogeneity provides the enzyme with a mechanism for substrate selectivity that does not rely on physical gating of the active site. Rather, the disordered active site of the holo-enzyme increases the entropic barrier in substrate binding to the point where non-specific substrates are unable to reasonably overcome this barrier. It is only upon interacting with the biological substrate, IgA, that distal interactions between IgA and the enzyme, like the S6 and M26 IgAPs,^19,20,39^ collapse the active site to a single entropically favorable conformation ready for catalysis. The substrate that is directly adjacent to the active site in this activated binary complex is the IgA hinge and hence it is the only molecule that is proteolyzed.

### Conclusions and Outlook

This work provides an in-depth structural and functional analysis of the biochemical workings of the *T. ramosa* IgAP. This is the first time an IgAP has been shown to have a modular domain architecture, meaning that the enzyme can be minimized to its core protease domain without losing its structure or catalytic function. Reducing the enzyme down to its functional core provides a promising path towards a protein-based therapy for IgAN as this construct is not only a better enzyme compared to the full-length construct but also maintains its high substrate specificity. Through solving the crystal structure of the protease domain, a working hypothesis for the structural basis of the enzyme’s substrate specificity was developed. Unlike the two other IgAP families, we propose that the *T. ramosa* IgAP is not gated by physical loops that occlude the active site but instead relies on an increased entropic barrier for substrate binding to limit non-specific proteolysis. The modularity of the enzyme also provided an opportunity to elucidate the function of the non-proteolytic domains that surround the M64 fold, which has previously been overlooked. We propose that the C-terminal tail contains several periscope domains that project the proteolytic domain out away from the surface of the bacterium; this is the first report of an IgAP associated with this type of extended protein stalk. However, whether some of these uncharacterized metal-associated domains (CTD2 and CTD4) have other functional roles involved in the bacterium’s colonization and survival in the gut is an important future line of investigation.

## Online Methods

### General Information

Bacterial growth media components were purchased from Bioshop Canada (Burlington, ON, Canada). Chemicals used for protein purification and crystallization were purchased from ChemImpex Inc. (Wood Dale, IL, USA) with the exception of IgePal-CA630, which was purchased from Thermo Fisher Scientific (Waltham, MA, USA). Crystallization screens were purchased from Molecular Dimensions (Holland, OH, USA). Crystal mounts, loops, and equipment (UniPuck system) were purchased from MiTeGen (Ithaca, NY, USA). All SDS-PAGE stain and destain reagents were purchased from BioShop (Burlington, ON, Canada).

### Expression Vector Cloning

The *T. ramosa* IgAP sequence (UniProt accession #Q9AES2; GenBank accession#AY028440) was used and modified as the open reading frame for all protein expression vectors. Protein expression constructs for the NTD (residues 31-313), MD (329-807), MD+CTD1 (329-876), CTD2 (880-1008), NTD+MD (31-807), MD+CTD1-4 (329-1195), CTD1-4 (808-1170), and full-length enzyme (31-1195) were codon optimized for *Escherichia coli*, synthesized *de novo*, and cloned into their respective vectors by GenScript (Piscataway, NJ, USA). To prevent proteolytic issues in purification,^41^ the FL and MD+CTD1-4 inserts were cloned into a dual-tagged vector (modified pJ401; DNA2.0; Menlo Park, CA, USA). These inserts were cloned into a pJ401 vector using dual *BamHI* cut sites (GenScript) C-terminal to a maltose-binding protein (MBP) tag and a thrombin cut site and N-terminal to a hexahistidine tag (His-tag). The rest of the constructs were subcloned from the full-length pJ401 construct into a SUMOstar vector (LifeSensors Inc.; Malvern, PA, USA) using *BsaI* and *XhoI* cut sites (GenScript) C-terminal to a His-tag and small ubiquitin-related modifier (SUMO) tag. Full vector and primary amino acid sequences for all constructs are listed in *Suppl. Table 4*.

### Protein Expression and Purification

Heterologous expression for all constructs was done in *E. coli* BL21(DE3) cells. A 16-to-18-hour LB overnight culture grown at 37°C was inoculated into ZYP-5052 autoinduction media^73^ at a ratio of 50 mL overnight culture to 1 L final media volume with a minimum headspace-to-media ratio of 4:1. All media were supplemented with 50 μg/mL kanamycin and the autoinduction media was supplemented with the recommended amount of trace metals.^73^ Cells were grown at 20°C and shaken at 150 rpm for 20-24 hours before being harvested via centrifugation at 6,000xg and stored as cell pellets at −80°C.

All pJ401 constructs were purified with an identical procedure. This purification protocol was similar but distinct to the protocol used for all SUMOstar constructs. All purification steps were carried out at 4°C, unless otherwise specified, and pH values were set at room temperature. The initial purification steps for pJ401 and SUMOstar constructs were the same: cell pellets were thawed in Buffer A (25 mM HEPES, pH 7.5, 0.5 M NaCl, 10 mM imidazole, 1 mM TCEP), passed twice through a FRENCH Pressure Cell (Thermo Fisher Scientific; Waltham, MA, USA) at 1,100 psi for cell lysis, and cell debris was removed via high-speed centrifugation at 17,000xg. The clarified cell lysate was then incubated with NiNTA resin (Qiagen; Hilden, Germany), equilibrated in Buffer A, for one hour. The resin was first washed with 25 column volumes (CVs) of Buffer B (25 mM HEPES, pH 7.5, 0.1% (v/v) IgePal CA-630, 10 mM imidazole, 1 mM TCEP) to remove non-specific hydrophobically bound contaminants followed by 40 CVs of Buffer A. The protein was eluted with Buffer C (25 mM HEPES, pH 7.5, 0.5 M NaCl, 300 mM imidazole, 1 mM TCEP).

For constructs in pJ401 (FL and MD+CTD1-4), the protein was then incubated with amylose resin (NEB; Ipswich, MA, USA), equilibrated in Buffer D (25 mM HEPES, pH 7.5, 0.5 M NaCl, 1 mM TCEP), for one hour. The amylose resin was washed with 20 CVs of Buffer D before being eluted with Buffer E (25 mM HEPES, pH 7.5, 0.5 M NaCl, 20% (w/v) glucose, 1 mM TCEP). The elution was concentrated to ∼3 mL with a 10 kDa molecular-weight-cut-off (MWCO) stirred-cell and centrifugal concentrator (Millipore Sigma; Burlington, MA, USA). Three rounds of 15 mL Buffer D washes were performed via centrifugal concentration for buffer exchange to remove the glucose. The protein was then subsequently digested with ∼2 mg bovine thrombin (BioPharm; Bluffdale, UT, USA) overnight at room temperature before being treated with 1 mM phenylmethylsulfonyl fluoride (PMSF; 100 mM stock in 100% ethanol) and incubated with two consecutive rounds of amylose resin for an hour each, equilibrated in Buffer D, to remove residual uncut protein and MBP. The final flowthrough was concentrated to less than 1 mL and loaded onto a pre-packed Superdex 200 Increase 10/300 GL column (Cytiva; Marlborough, MA, USA), equilibrated in Buffer D, run at 0.5 mL/min. Fractions not in the void volume peak were concentrated to less than 1 mL and injected onto a pre-packed HiLoad Superdex 75 pg 16/600 (Cytiva), equilibrated in Storage/Crystallization Buffer (25 mM HEPES, pH 7.5 and 1 mM TCEP), and run at 0.5 mL/min.

For constructs in SUMOstar (with the exception of CTD2, see below), the protein eluted off of the first NiNTA resin was digested with SUMO protease overnight (expressed and purified in-house from Addgene plasmid pCDB302).^74^ The cleaved protein was dialyzed against Buffer D for a minimum of 6 hours followed by another round of dialysis overnight to remove residual imidazole. The protein was then incubated with a second round of NiNTA resin, equilibrated in Buffer D, to remove the cleaved SUMO tag and any remaining uncleaved fusion. NiNTA flowthrough was concentrated to less than 1 mL and loaded onto a pre-packed HiLoad Superdex 75 pg 16/600, pre-equilibrated in Storage Buffer, run at 0.5 mL/min.

As the CTD2 NiNTA elution had a green hue, ZnCl_2_ was added to the elution to a final concentration of 1 mM. After an hour of incubation, the solution became colorless. To prevent residual ZnCl_2_ from interfering with the second NiNTA step, the elution was concentrated to ∼3 mL, loaded onto a manually packed P-6DG desalting column (BioRad; Hercules, CA, USA) equilibrated in Buffer D, before being incubated with a second round of NiNTA resin and continuing on from the protocol for SUMOstar above.

The final purity of all proteins was qualitatively analyzed using SDS-PAGE. Pure fractions eluting from the final size-exclusion column were concentrated, frozen in 20-30 μL pellets via direct immersion in liquid nitrogen, and stored at −80°C. Protein concentration was measured using 1% mass extinction coefficients calculated from their primary sequence using the ProtParam program in Expasy (*Suppl. Table 4*).^75^

### Small-Angle X-ray Scattering Data Collection and Processing

SAXS data were collected at the Cornell High-Energy Synchrotron Source (CHESS) ID7A beamline using an X-ray wavelength of 1.1013 Å and an EIGER 4M detector.^76^ All proteins were dialyzed against 25 mM HEPES, pH 7.5 with 1 mM TCEP overnight using a 12-14 kDa MWCO D-Tube Mini Dialyzer (Millipore Sigma) to obtain a matching solution for buffer subtraction. Protein concentration was remeasured after dialysis as previously described. A 30 μL volume of sample was injected into the sample capillary for analysis. Samples were oscillated within the capillary during exposure to reduce radiation damage (2-3 oscillations/sec). 15 images were collected for each sample, with the detector being exposed for one sec/image with no attenuation.

Images were azimuthally averaged and buffer subtracted in RAW (version 2.1.4).^77^ Processed data were subsequently analyzed in RAW for Guinier and Kratky analysis,^77^ GNOM (ATSAS package, version 3.2.1) for P(r) analysis,^78^ and FoXS for simulating model scatter (web server, accessed February 27^th^, 2024).^49^ The P(r) function was calculated using all measured data. The P(r) function was only calculated on truncated data (high q cutoff at ∼0.3 or 0.4) if the P(r) fit to the scattering curve showed skewed residuals at q > 0.3 or 0.4, respectively. Data collection and statistics for all SAXS data are summarized in *Suppl. Table 1*.

CORAL was used to optimize the multidomain models to the SAXS scattering curves.^48^ Where applicable, the individual domains of the NTD, MD+CTD1, CTD2, CTD3, and CTD4 were modeled with flexible linker lengths of 15, 4, 4, and 5 residues between each domain, respectively. The full-length enzyme was the only construct that was modeled with a 29-residue flexible region at its C-terminus. The MD+CTD1 was modeled using its crystal structure and NTD, CTD2, CTD3, and CTD4 were modeled with their AlphaFold2-predicted models,^42,43^ which were truncated based on the boundaries outlined previously.

### Preparation of the Substrate, Enzymes, and Internal Standards for the Kinetic Assay

Monomeric myeloma IgA1 (Athens Research and Technology, Athens, GA, USA) was loaded onto a pre-packed Superdex 200 Increase 10/300 GL column (Cytiva; Marlborough, MA, USA), equilibrated in 25 mM HEPES, pH 7.5, and run at 0.5 mL/min. All buffer-exchanged fractions containing IgA1 were pooled, concentrated with a 50 kDa molecular-weight-cutoff (MWCO) centrifugal concentrator (Millipore Sigma; Burlington, MA, USA), frozen in 20 μL pellets via direct immersion in liquid nitrogen, and stored at −80°C. Protein concentration was measured using the 1% mass extinction coefficient of 13.2, as provided by the manufacturer. IgA1 was used at the frozen concentration (for a 38 μM IgA1 reaction) and diluted with 25 mM HEPES, pH 7.5 accordingly for reactions containing 1, 3, 5, 10, and 20 μM IgA1.

30 μL of each *T. ramosa* IgAP enzyme construct was diluted to 500 μL with 25 mM HEPES, pH 7.5 + 1 mM TCEP in the presence of 25% (v/v) Chelex 100 resin (BioRad) and was mixed overnight at room temperature. Chelex 100 resin was prepared and equilibrated by adding fresh Chelex 100 resin to a gravity column, washed with 50 CVs of 18.2 MΩcm water, 10 CVs of 0.5 M sodium phosphate buffer, pH 6.3, 50 CVs of 18.2 MΩcm water. Chelex 100 resin was separated by light centrifugation at 1,000xg for 5 minutes and the supernatant was incubated with 1 mM ZnCl_2_ for two hours at room temperature. The zinc-supplemented supernatant was then desalted back into 25 mM HEPES, pH 7.5 + 1 mM TCEP with a 0.5 mL 7 kDa MWCO Zeba Spin Desalting Column (Thermo Scientific, Waltham, MA, USA) and the enzyme concentration was remeasured using 1% mass extinction coefficients theoretically determined from their primary sequence using the ProtParam program in Expasy (*Suppl. Table 4*)^75^ prior to use in the reaction mixture.

250 μL of 1 μM IgA1 and 500 μL of 2 μM IgA1 were incubated with 40 nM *T. ramosa* IgAP MD overnight at 37°C in 25 mM HEPES, pH 7.5. 25 μL samples were prepared by mixing 20 μL reaction volume with 5 μL 4X SDS-PAGE loading dye (0.2 M Tris, pH 6.8 + 8% (w/v) SDS + 0.4% (w/v) bromophenol blue + 40% (v/v) glycerol + 4% (v/v) β-mercaptoethanol + 50 mM EDTA). Samples were boiled for 5 min at 95°C in a dry-heat block and spun down for 1 min at 17,000xg prior to being frozen at −20°C. Aliquots were thawed as needed for each gel. 1 μM internal standards were only used for gels that contained 20 pmol IgA1 and 2 μM internal standards were used for all other gels.

### Gel-Based Kinetic Assay for Quantifying IgAP Activity

Various enzyme constructs were incubated with 1, 3, 5, 10, 20, and 38 μM IgA1 at 37°C in 25 mM HEPES, pH 7.5. 0.15, 0.5, 0.75, or 1.5 nM enzyme was used for all MD, MD+CTD1, NTD+MD, and FL reactions, respectively, with the exception of the 38 μM IgA1 reaction for MD where 0.5 nM enzyme was used. Five samples were taken from the same reaction at 30-minute time points over the course of two and a half hours. The reaction was quenched with the addition of 4X SDS-PAGE loading dye and samples were boiled for 5 min at 95°C in a dry-heat block. All samples were spun down for 1 min at 17,000xg prior to being loaded in their entirety onto the gel with gel-loading tips. All samples were prepared in 25 μL containing 20 μL reaction volume and 5 μL 4X SDS-PAGE loading dye. Reactions that contained more than 2 μM IgA1 were diluted with the reaction buffer such that the final sample contained 40 pmol of IgA1. Samples for the 1 μM IgA1 reactions were not diluted to make the sample (*i.e.*, 20 μL of the reaction was added directly to the sample).

Gels were cast with a 12% (w/v) resolving and 4% (w/v) stacking layer using a 30% (w/v) stock 37.5:1 acrylamide:bis-acrylamide solution (BioRad). The BioRad 1.0 mm-thickness 10-well Mini-PROTEAN and Tetra Cell electrophoresis systems were used for casting and running the gels, respectively. The resolving layer was cast such that it stopped 2 cm below the top of the glass short plate. Gels were initially run at 80 V before being ramped up to 160 V when the dye front encountered the separating gel (approximately 30 min). The gels were stopped when the dye front ran off the bottom of the gel (approximately 60 min from the start of the 160V phase).

100 mL of Whatman Grade 1-filtered (Cytiva) SDS-PAGE stain (0.25% (w/v) Coomassie R-250 + 50% (v/v) methanol + 10% (v/v) glacial acetic acid) was added to the gels, microwaved for 1 min, and left to stain on a rocker for 5 min. 200 mL of SDS-PAGE destain (20% (v/v) methanol + 10% (v/v) glacial acetic acid) was then added, microwaved for 1 min, and the gels were left to destain on a rocker for two hours at room temperature. The gels were then immediately imaged on a Gel Doc XR+ gel imager (BioRad) with a white-light transilluminator.

Gel images were processed with the BioRad Image Lab (version 6.1.0, build 7) program. The band intensities for the heavy-chain fragment associated with the IgA1 Fc (H1) were integrated using the program’s built-in densitometry features. The background-corrected H1 band intensity was divided by that of the H1 band in the internal standard to calculate the percentage of IgA1 cleaved at every time point. This percentage was plotted with respect to time and a simple linear regression was used to calculate the rate of the reaction (in units of molarity of IgA1 cleaved per minute) from multiplying the slope of the regression (in units of percentage of IgA1 cleaved per minute) by the concentration of substrate in the reaction. The rate was then normalized by the enzyme concentration, plotted as a function of IgA1 concentration, and fit to the Michaelis-Menten equation to extract kinetic parameters. k_cat_ and KM parameters were fit and their associated standard error were calculated using the Enzyme Kinetics module (version 1.3) in SigmaPlot (version 11.0). Standard errors for k_cat_/KM were calculated using statistical equations for the propagation of error.

### Intact Peptide Mass Spectrometry

A synthetic peptide (N-VPCPVPST-C; GenScript, Piscataway, NJ, USA) based on the *T. ramosa* IgAP IgA1 cleavage site was dissolved in 25 mM HEPES, pH 7.5 + 1 mM TCEP and frozen in small-volume aliquots at −20°C for storage. 10 nM full-length or MD *T. ramosa* IgAP was incubated with 60 μM peptide in 25 mM HEPES, pH 7.5 + 1 mM TCEP in a reaction volume of 20 μL for 18 hours at 37°C. Peptide without enzyme was also subjected to the same conditions as the negative control.

The entire reaction volume was subjected to a C_18_ ZipTip (Millipore Sigma) to prepare each sample for mass spectrometry. The sample was first acidified with trifluoroacetic acid (TFA; Millipore Sigma) to a final concentration of 0.5% (v/v). The C_18_ resin was wetted with 100% (v/v) acetonitrile and then equilibrated in 0.1% (v/v) TFA, before allowing the peptides in the sample to bind. After washing the resin with 0.1% (v/v) TFA, the bound peptides were desorbed with a 5 μL mixture of 1% (v/v) formic acid and 50% (v/v) methanol. 2 μL sample was mixed with a 2 μL of α-cyano-4-hydroxycinnamic acid (HCCA; made in-house at the University of Waterloo Mass Spectrometry Facility) as the matrix, dried onto the target plate, and subjected to MALDI-ToF mass spectrometry on a Bruker Autoflex Speed MALDI instrument (University of Waterloo, Department of Chemistry Mass Spectrometry Facility).

### Protein Crystallization

High-throughput crystallization trials for MD+CTD1 were carried out with commercially available screens in small-volume sitting-drop trays using a Crystal Gryphon LCP robot (Art Robbins Instruments; Sunnyvale, CA, USA). Drops consisting of 0.2 μL protein sample (10 mg/mL protein in Crystallization Buffer) were mixed with 0.2 μL reservoir solution and left to equilibrate at room temperature for a few weeks. The reservoir volume was 30 μL. Initial hits grew after one to two weeks in 0.1 M Bis-Tris, pH 6.5 and 25% (w/v) PEG 3350 (MCSG1 C12).^79^ As these crystals were not suitable for single-crystal diffraction, the conditions were further optimized by an additive screen (Hampton Research; Aliso Viejo, CA, USA) at a ratio of 2 μL protein to 1.8 μL mother liquor and 0.2 μL non-volatile additive. Clusters of hollow cylindrical crystals resulted from a mix of 2.0 μL protein sample (8 mg/mL), 1.8 μL reservoir solution (0.1 M Bis-Tris, pH 6.5 and 12% (v/v) PEG 3350), and 0.2 μL 0.1 M praseodymium (III) acetate. These crystal clusters were manually manipulated to acquire single crystals suitable for diffraction. All crystals were cryoprotected in their respective reservoir solutions supplemented with 20% (v/v) glycerol. Due to the fragile nature of the crystals, they were cryoprotected stepwise with increasing concentrations of glycerol, from 5% to 20% (v/v) in steps of 5%.

### Metal-Chelating and Zinc-Supplementing Crystallographic Experiments

To obtain crystal structures of the enzyme bound to only zinc, Chelex 100 resin (BioRad) prepared as described above was directly added to protein solutions containing MD+CTD1 enzyme and left to incubate overnight at room temperature. Chelex 100 resin was also added directly to all solutions used for crystallization and cryoprotection, with the exception of the crystallization additives (praseodymium (III) acetate and ZnCl_2_). All crystallization buffers and precipitant solutions were made with 18.2 MΩcm water. Chelex 100 resin that was added directly to any protein solution was further equilibrated in 50 CVs of Crystallization Buffer prior to use.

Crystal growth and manipulation were done as previously described with the exception of adding a small amount of Chelex 100 resin to the drop during crystallization. To supplement the crystals with zinc, metal-chelated single crystals were soaked in large 100 μL drops containing the mother liquor supplemented with 1 mM ZnCl_2_ (Hampton Research) overnight at room temperature. Crystallization additive was not added to the soak solutions. Zinc-soaked crystals were then cryoprotected stepwise as described above with 1 mM ZnCl_2_ added to all solutions.

### Crystallographic Data Collection and Structure Refinement

Diffraction data for all MD+CTD1 crystals were collected at the Canadian Light Source BM beamline on a Detectris Pilatus3 S 6M. To confirm the location and identity of metals in the MD+CTD1 crystal structures, full datasets were collected in succession at the same spot on each crystal at three different wavelengths. A high-resolution native dataset was collected with an exposure of 0.2°/0.4 sec/frame as the primary dataset for model building and refinement. A second dataset was collected at an energy 150-200 keV above the zinc absorption edge (9.6586 keV) and a third dataset was collected at an energy 150-200 keV below the zinc absorption edge, both with an exposure of 0.2°/0.05 sec/frame.

All data were indexed, integrated, and scaled with DIALS (version 3.8.0)^80^ and imported into the CCP4i suite (version 8.0.009)^81^ with AIMLESS (version 0.7.9).^82^ Phases were solved using molecular replacement (MOLREP version 11.9.02)^83^ with its corresponding AlphaFold2 model.^42,43^ Refinement was done using phenix.refine (version 1.20.1_4487)^84^ in conjunction with manual model building in COOT (version 0.8.9.2).^85^ B-factors were refined isotropically with automatically determined translation-libration-screw parameters for all structures. Alternate conformer and metal occupancies were also refined with phenix.refine.^84^ Model geometry was analyzed and optimized based on suggestions by MolProbity (version 4.5.2).^86^ Anomalous maps were generated with the cad^81^ and fft^87–89^ programs in the CCP4i suite using the finalized refined structures of their respective high-resolution datasets. Data collection and model statistics for all structures are summarized in *Suppl. Table 2*.

## Acknowledgements

Part of the research described in this paper was performed using beamline CMCF-BM at the Canadian Light Source, a national research facility of the University of Saskatchewan, which is supported by the Canada Foundation for Innovation (CFI), the Natural Sciences and Engineering Research Council (NSERC), the National Research Council (NRC), the Canadian Institutes of Health Research (CIHR), the Government of Saskatchewan, and the University of Saskatchewan. The SAXS data described in the paper was collected at the Center for High-Energy X-ray Sciences (CHEXS), which is supported by the National Science Foundation (BIO, ENG and MPS Directorates) under award DMR-2342336, and the Macromolecular Diffraction at CHESS (MacCHESS) facility, which is supported by award 1-P30-GM124166 from the National Institute of General Medical Sciences and the National Institutes of Health.

## Data Availability

The model coordinates and structure factors for the as-purified, metal-chelated, and zinc-supplemented *T. ramosa* IgAP MD+CTD1 structures have been deposited in the PDB (www.rcsb.org/pdb) under accession codes 9EKK, 9EKM, and 9EKN, respectively. SAXS data for the NTD, MD, MD+CTD1, CTD2, NTD+MD, CTD1-4, MD+CTD1-4, and full-length constructs have been deposited in the SASBDB (www.sasbdb.org) under accession codes SASDWH3, SASDWJ3, SASDWK3, SASDWL3, SASDWM3, SASDWN3, SASDWP3, and SASDWQ3, respectively.

## References

1. Mistry, D. & Stockley, R. A. IgA1 protease. Int J Biochem Cell Biol 38, 1244–1248 (2006).

2. Mehta, S. K., Plaut, A. G., Calvanico, N. J. & Tomasi, T. B. Human Immunoglobulin A: Production of an Fc Fragment by an Enteric Microbial Proteolytic Enzyme. J Immunol 111, 1274–1276 (1973).

3. Casadevall, A. & Pirofski, L.-A. Host-Pathogen Interactions: The Attributes of Virulence. J Infect Dis 184, 337–344 (2001).

4. Mulks, M. H., Kornfeld, S. J., Frangione, B. & Plaut, A. G. Relationship Between the Specificity of IgA Proteases and Serotypes in Haemophilus influenzae. J Infect Dis 146, 266–274 (1982).

5. Kilian, M., Mestecky, J. & Schrohenloher, R. E. Pathogenic Species of the Genus Haemophilus and Streptococcus pneumoniae Produce Immunoglobulin A1 Protease. Infect Immun 26, 143–149 (1979).

6. Mulks, M. H. & Plaut, A. G. IgA Protease Production as a Characteristic Distinguishing Pathogenic from Harmless Neisseriaceae. N Engl J Med 299, 973–976 (1978).

7. Mansa, B. & Kilian, M. Retained Antigen-Binding Activity of Fab(α) Fragments of Human Monoclonal Immunoglobulin A1 (IgA1) Cleaved by IgA1 Protease. Infect Immun 52, 171–174 (1986).

8. Male, C. J. Immunoglobulin A1 Protease Production by Haemophilus influenzae and Streptococcus pneumoniae. Infect Immun 26, 254–261 (1979).

9. Troeger, C. et al. Estimates of the global, regional, and national morbidity, mortality, and aetiologies of lower respiratory infections in 195 countries, 1990–2016: a systematic analysis for the Global Burden of Disease Study 2016. Lancet Infect Dis 18, 1191–1210 (2018).

10. Brooks, L. R. K. & Mias, G. I. Streptococcus pneumoniae’s Virulence and Host Immunity: Aging, Diagnostics, and Prevention. Front Immunol 9, (2018).

11. Liu, L. et al. Global, regional, and national causes of under-5 mortality in 2000–15: an updated systematic analysis with implications for the Sustainable Development Goals. The Lancet 388, 3027–3035 (2016).

12. Bachovchin, W. W., Plaut, A. G., Flentke, G. R., Lynch, M. & Kettner, C. A. Inhibition of IgA1 Proteinases from Neisseria gonorrhoeae and Haemophilus influenzae by Peptide Prolyl Boronic Acids. Journal of Biological Chemistry 265, 3738–3743 (1990).

13. Garner, A. L., Fullagar, J. L., Day, J. A., Cohen, S. M. & Janda, K. D. Development of a high-throughput screen and its use in the discovery of Streptococcus pneumoniae immunoglobulin A1 protease inhibitors. J Am Chem Soc 135, 10014–10017 (2013).

14. Choudary, S. K., Qiu, J., Plaut, A. G. & Kritzer, J. A. Versatile Substrates and Probes for IgA1 Protease Activity. ChemBioChem 14, 2007–2012 (2013).

15. Shehaj, L. et al. Small-Molecule Inhibitors of Haemophilus influenzae IgA1 Protease. ACS Infect Dis 5, 1129–1138 (2019).

16. Örtqvist, Å. Treatment of community-acquired lower respiratory tract infections in adults. European Respiratory Journal 20, S40–53 (2002).

17. World Health Organization. Global Antimicrobial Resistance Surveillance System (GLASS) Report Early Implementation.

18. Plaut, A. G., Genco, R. J. & Tomasi, T. B. Isolation of an Enzyme from Streptococcus sanguis Which Specifically Cleaves IgA. J Immunol 113, 289–291 (1974).

19. Johnson, T. A., Qiu, J., Plaut, A. G. & Holyoak, T. Active site gating regulates substrate selectivity in a chymotrypsin-like serine protease. The structure of Haemophilus influenzae IgA1 protease. J Mol Biol 389, 559–574 (2009).

20. 20. Redzic, J. S., et al. A substrate-induced gating mechanism is conserved among Gram-positive IgA1 metalloproteases. Commun Biol 5, (2022).

21. Fujiyama, Y. et al. A Novel IgA Protease from Clostridium sp. Capable of Cleaving IgA1 and IgA2 A2m(1) but not IgA2 A2m(2) allotype paraproteins. The Journal of Immunology 134, 573–576 (1985).

22. Kosowska, K. et al. The Clostridium ramosum IgA Proteinase Represents a Novel Type of Metalloendopeptidase. J Biol Chem 277, 11987–11994 (2002).

23. Qiu, J., Brackee, G. P. & Plaut, A. G. Analysis of the Specificity of Bacterial Immunoglobulin A (IgA) Proteases by a Comparative Study of Ape Serum IgAs as Substrates. Infect Immun 64, 933–937 (1996).

24. Xie, L.-S., Huang, J., Qin, W. & Fan, J.-M. Immunoglobulin A1 protease: A new therapeutic candidate for immunoglobulin A nephropathy. Nephrology 15, 584–586 (2010).

25. Kintzing, J. R., Filsinger Interrante, M. V. & Cochran, J. R. Emerging Strategies for Developing Next-Generation Protein Therapeutics for Cancer Treatment. Trends Pharmacol Sci 37, 993–1008 (2016).

26. Julian, B. A., Waldo, F. B., Rifai, A. & Mestecky, J. IgA Nephropathy, the Most Common Glomerulonephritis Worldwide - A Neglected Disease in the United States? Am J Med 84, 129– 132 (1988).

27. Lamm, M. E. et al. Microbial IgA protease removes IgA immune complexes from mouse glomeruli in vivo: Potential therapy for IgA nephropathy. Am J Pathol 172, 31–36 (2008).

28. Lechner, S. M. et al. IgA1 Protease Treatment Reverses Mesangial Deposits and Hematuria in a Model of IgA Nephropathy. Journal of the American Society of Nephrology 27, 2622–2629 (2016).

29. Wang, L. et al. Bacterial IgA protease-mediated degradation of agIgA1 and agIgA1 immune complexes as a potential therapy for IgA Nephropathy. Sci Rep 6, (2016).

30. Kosowska, K. et al. The Clostridium ramosum IgA proteinase represents a novel type of metalloendopeptidase. Journal of Biological Chemistry 277, 11987–11994 (2002).

31. Plaut, A. G., Gilbert, J. V, Artenstein, M. S. & Capra, J. D. Neisseria gonorrhoeae and Neisseria meningitidis: Extracellular Enzyme Cleaves Human Immunoglobulin A. Science (1979) 190, 1103– 1105 (1975).

32. Mortensen, S. B. & Kilian, M. Purification and Characterization of an Immunoglobulin A1 Protease from Bacteroides melaninogenicus. Infect Immun 45, 550–557 (1984).

33. Kilian, M., Thomsen, B., Petersen, T. E. & Bleeg, H. Molecular Biology of Haemophilus influenzae IgA1 Proteases. Mol Immunol 20, 1051–1058 (1983).

34. Spooner, R. K., Russell, W. C. & Thirkell, D. Characterization of the Immunoglobulin A Protease of Ureaplasma urealyticum. Infect Immun 60, 2544–2546 (1992).

35. Fujiyama, Y., Masayoshi, I., Hodohara, K., Hosoda, S. & Kobayashi, K. The Site of Cleavage in Human Alpha Chains of IgA1 and IgA2:A2m(1) Allotype Paraproteins by the Clostridial IgA Protease. Mol Immunol 23, 147–150 (1986).

36. Rawlings, N. D., Barrett, A. J. & Bateman, A. MEROPS: The database of proteolytic enzymes, their substrates and inhibitors. Nucleic Acids Res 40, (2012).

37. Rawlings, N. D. & Bateman, A. How to use the MEROPS database and website to help understand peptidase specificity. Protein Science 30, 83–92 (2021).

38. Johnson, T. A., Qiu, J., Plaut, A. G. & Holyoak, T. Active-Site Gating Regulates Substrate Selectivity in a Chymotrypsin-Like Serine Protease. The Structure of Haemophilus influenzae Immunoglobulin A1 Protease. J Mol Biol 389, 559–574 (2009).

39. Wang, Z. et al. Mechanism and inhibition of Streptococcus pneumoniae IgA1 protease. Nat Commun 11, (2020).

40. Lawson, P. A., Perez, L. S. & Sankaranarayanan, K. Reclassification of Clostridium cocleatum, Clostridium ramosum, Clostridium spiroforme and Clostridium saccharogumia as Thomasclavelia cocleata gen. nov., comb. nov., Thomasclavelia ramosa comb. nov., gen. nov., Thomasclavelia spiroformis comb. nov. and Thomasclavelia saccharogumia comb. nov. Int J Syst Evol Microbiol 73, (2023).

41. Xie, X. et al. Chimeric Fusion between Clostridium Ramosum IgA Protease and IgG Fc Provides Long-Lasting Clearance of IgA Deposits in Mouse Models of IgA Nephropathy. Journal of the American Society of Nephrology 33, 918–935 (2022).

42. Jumper, J. et al. Highly accurate protein structure prediction with AlphaFold. Nature 596, 583– 589 (2021).

43. Mirdita, M. et al. ColabFold: making protein folding accessible to all. Nat Methods 19, 679–682 (2022).

44. Krishna, S. S., Majumdar, I. & Grishin, N. V. Structural classification of zinc fingers. Nucleic Acids Res 31, 532–550 (2003).

45. Krissinel, E. & Henrick, K. Inference of Macromolecular Assemblies from Crystalline State. J Mol Biol 372, 774–797 (2007).

46. Baek, M. et al. Accurate prediction of protein structures and interactions using a three-track neural network. Science (1979) 373, 871–876 (2021).

47. Jacques, D. A. & Trewhella, J. Small-angle scattering for structural biology - Expanding the frontier while avoiding the pitfalls. Protein Science 19, 642–657 (2010).

48. Petoukhov, M. V. et al. New developments in the ATSAS program package for small-angle scattering data analysis. J Appl Crystallogr 45, 342–350 (2012).

49. Schneidman-Duhovny, D., Hammel, M., Tainer, J. A. & Sali, A. Accurate SAXS Profile Computation and its Assessment by Contrast Variation Experiments. Biophys J 105, 962–974 (2013).

50. Doolittle, R. F. The Multiplicity of Domains in Proteins. AnnlL Rev. Biochem 64, 287–314 (1995).

51. Krissinel, E. & Henrick, K. Secondary-structure matching (SSM), a new tool for fast protein structure alignment in three dimensions. Acta Crystallogr D Biol Crystallogr 60, 2256–2268 (2004).

52. Ebbes, M. et al. Fold and Function of the InlB B-repeat. Journal of Biological Chemistry 286, 15496–15506 (2011).

53. Copp, J., Marino, M., Banerjee, M., Ghosh, P. & Van der Geer, P. Multiple Regions of Internalin B Contribute to Its Ability to Turn on the Ras-Mitogen-activated Protein Kinase Pathway. Journal of Biological Chemistry 278, 7783–7789 (2003).

54. Bogomolovas, J. et al. Exploration of pathomechanisms triggered by a single-nucleotide polymorphism in titin’s I-band: The cardiomyopathy-linked mutation T2580I. Open Biol 6, (2016).

55. Whelan, F. The long and the short of Periscope Proteins. Biochem Soc Trans 50, 1293–1302 (2022).

56. Bateman, A., Holden, M. T. G. & Yeats, C. The G5 domain: a potential N-acetylglucosamine recognition domain involved in biofilm formation. Bioinformatics 21, 1301–1303 (2005).

57. Roche, F. M., Meehan, M. & Foster, T. J. The Staphylococcus aureus surface protein SasG and its homologues promote bacterial adherence to human desquamated nasal epithelial cells. Microbiology 149, 2759–2767 (2003).

58. Paukovich, N. et al. Streptococcus pneumoniae G5 domains bind different ligands. Protein Science 28, 1797–1805 (2019).

59. Conrady, D. G., Wilson, J. J. & Herr, A. B. Structural basis for Zn2+-dependent intercellular adhesion in staphylococcal biofilms. Proc Natl Acad Sci U S A 110, E202–E211 (2013).

60. Whelan, F. et al. Periscope Proteins are variable-length regulators of bacterial cell surface interactions. Proc Natl Acad Sci U S A 118, (2021).

61. Chaton, C. T. & Herr, A. B. Defining the metal specificity of a multifunctional biofilm adhesion protein. Protein Science 26, 1964–1973 (2017).

62. Mertens, H. D. T. & Svergun, D. I. Structural characterization of proteins and complexes using small-angle X-ray solution scattering. J Struct Biol 172, 128–141 (2010).

63. Paysan-Lafosse, T. et al. InterPro in 2022. Nucleic Acids Res 51, D418–427 (2023).

64. Baykov, I. K. et al. Tentaclins—A Novel Family of Phage Receptor-Binding Proteins That Can Be Hypermutated by DGR Systems. Int J Mol Sci 24, (2023).

65. Steinrücke, P. et al. Design of helical proteins for real-time endoprotease assays. Anal Biochem 286, 26–34 (2000).

66. Wood, S. G. & Burton, J. Synthetic Peptide Substrates for the Immunoglobulin A1 Protease from Neisseria gonorrhoeae (Type 2). Infect Immun 59, 1818–1822 (1991).

67. Northrop, D. B. On the Meaning of Km and V/K in Enzyme Kinetics. J Chem Educ 75, 1153 (1998).

68. Cerdà-Costa, N. & Gomis-Rüth, F. X. Architecture and function of metallopeptidase catalytic domains. Protein Science 23, 123–144 (2014).

69. Bertini, I. et al. Snapshots of the Reaction Mechanism of Matrix Metalloproteinases. Angew Chem Int Ed 45, 7952–7955 (2006).

70. Gomis-Rüth, F.-X., Kress, L. F. & Bode, W. First structure of a snake venom metalloproteinase: a prototype for matrix metalloproteinases/collagenases. EMBO J 12, 4151–4157 (1993).

71. Bode, W., Gomis-Rüth, F. X., Zwilling, R. & Stöcker, W. Structure of astacin and implications for activation of astacins and zinc-ligation of collagenases. Nature 358, 164–167 (1992).

72. Handing, K. B. et al. Characterizing metal-binding sites in proteins with X-ray crystallography. Nat Protoc 13, 1062–1090 (2018).

73. Studier, F. W. Protein production by auto-induction in high density shaking cultures. Protein Expr Purif 41, 207–234 (2005).

74. Lau, Y.-T. K. et al. Discovery and engineering of enhanced SUMO protease enzymes. Journal of Biological Chemistry 293, 13224–13233 (2018).

75. Gasteiger, E. et al. Protein Analysis Tools on the ExPASy Server. in The Proteomics Protocols Handbook (ed. Walker, J. M.) 571–607 (Humana Press Inc., Totowa, 2005).

76. Acerbo, A. S., Cook, M. J. & Gillilan, R. E. Upgrade of MacCHESS facility for X-ray scattering of biological macromolecules in solution. J Synchrotron Radiat 22, 180–186 (2015).

77. Hopkins, J. B., Gillilan, R. E. & Skou, S. BioXTAS RAW: improvements to a free open-source program for small-angle X-ray scattering data reduction and analysis. J Appl Crystallogr 50, 1545– 1553 (2017).

78. Svergun, D. I. Determination of the Regularization Parameter in Indirect-Transform Methods Using Perceptual Criteria. J. Appl. Cryst 25, 495–503 (1992).

79. Fazio, V. J., Peat, T. S. & Newman, J. A drunken search in crystallization space. Acta Crystallogr F70, 1303–1311 (2014).

80. Winter, G. et al. DIALS: Implementation and evaluation of a new integration package. Acta Crystallogr D74, 85–97 (2018).

81. Winn, M. D. et al. Overview of the CCP4 suite and current developments. Acta Crystallogr D67, 235–242 (2011).

82. Evans, P. R. & Murshudov, G. N. How good are my data and what is the resolution? Acta Crystallogr D69, 1204–1214 (2013).

83. Vagin, A. & Teplyakov, A. MOLREP: an Automated Program for Molecular Replacement. J Appl Crystallogr 30, 1022–1025 (1997).

84. Afonine, P. V. et al. Towards automated crystallographic structure refinement with phenix.refine. Acta Crystallogr D68, 352–367 (2012).

85. Emsley, P., Lohkamp, B., Scott, W. G. & Cowtan, K. Features and development of Coot. Acta Crystallogr D66, 486–501 (2010).

86. Williams, C. J. et al. MolProbity: More and better reference data for improved all-atom structure validation. Protein Science 27, 293–315 (2018).

87. Ten Eyck, L. F. Crystallographic Fast Fourier Transforms. Acta Cryst A29, 183–191 (1973).

88. Read, R. J. & Schierbeek, A. J. A phased translation function. J. Appl. Cryst 21, 490–495 (1988).

89. Immirzi, A. Crystallographic Computing Techniques. in (ed. Ahmed, F. R.) 399 (Munksgaard, 1966).

