## Supplemental information for "Structure of the *Thomasclavelia ramosa* immunoglobulin A protease reveals a modular and minimizable architecture distinct from other immunoglobulin A proteases"

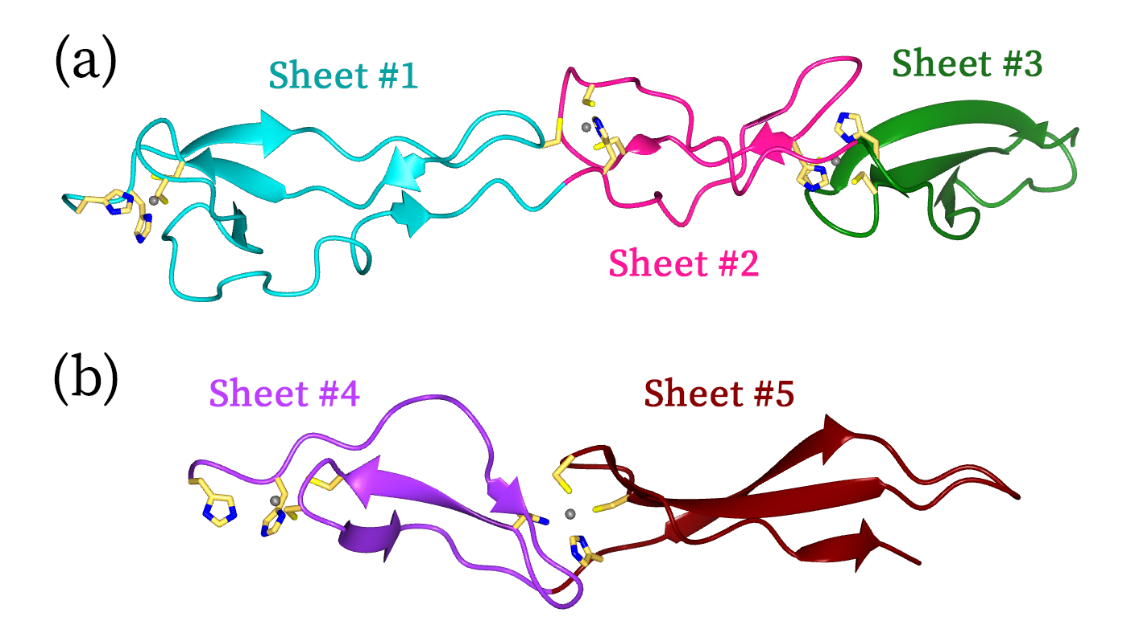

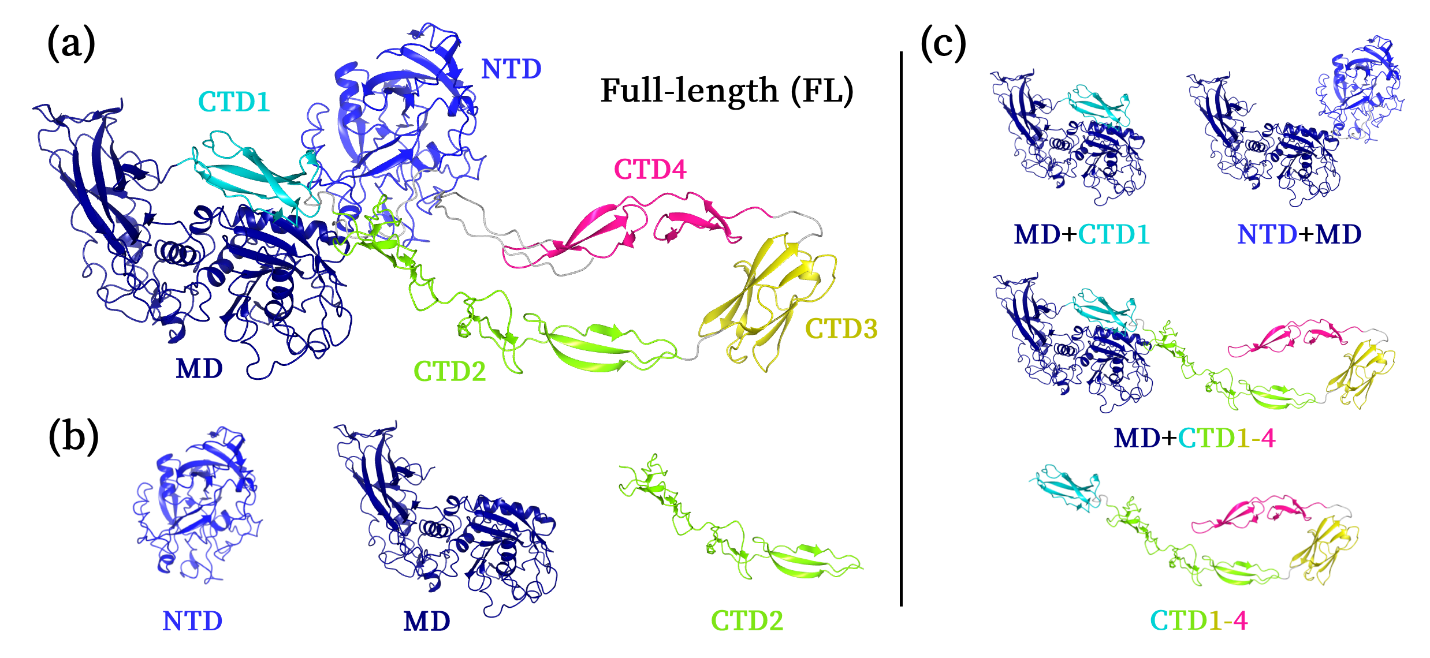

Supplementary Figure 2 – AlphaFold2 Model for the C-terminal Putative Zinc-Binding Domains. The CTD2 **(a)** and CTD4 **(b)** domains of *T. ramosa* IgAP contain several putative zinc-binding sites as a cluster of cysteine and histidine residues are modelled to be in close proximity. The side chains of these metal-binding residues are shown in their predicted conformation and zinc ions (grey spheres) were added post-hoc purely for visualization purposes. CTD2 contains two Cys_2_His_2_ and one Cys_3_His_1_ putative zinc-binding sites and CTD4 contains two Cys_2_His_2_ putative zinc-binding sites.

Supplementary Figure 1 – Putative Domains and Protein Constructs of the *T. ramosa* IgAP. (a) The full-length (FL) AlphaFold2 model was split into six putative domains linked together by flexible regions (grey): N-terminal domain (NTD; blue), middle domain or protease domain (MD; dark blue), C-terminal domain #1 (CTD1; cyan), C-terminal domain #2 (CTD2; green), C-terminal domain #3 (CTD3; yellow), and C-terminal domain #4 (CTD4; pink). These predictions formed the basis for the design of all *T. ramosa* IgAP **(b)** single-domain and **(c)** multi-domain protein constructs.H

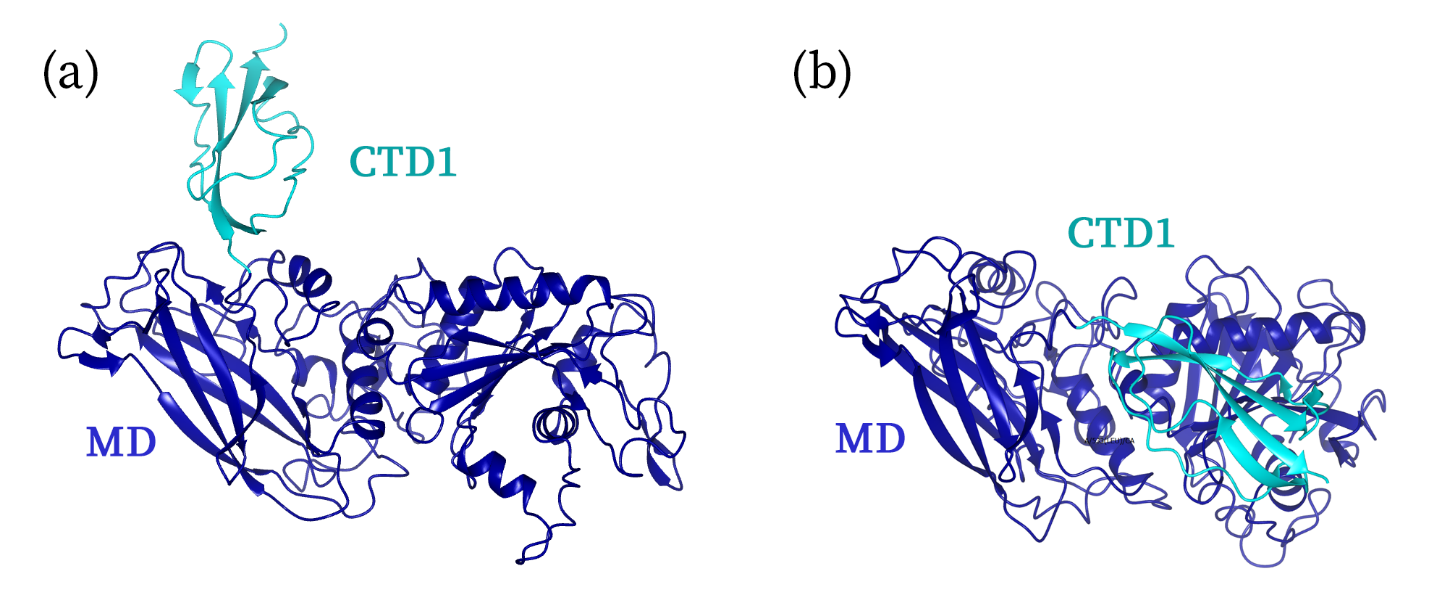

Supplementary Figure 3 – RosettaFold Model for the Protease Domain and Its Adjacent C-terminal Domain. The CTD1 of the *T. ramosa* IgAP is predicted to be a separate domain in the RosettaFold model **(a)** but not in the AlphaFold2 model **(b)**. This suggests that CTD1 might be a true domain despite the predicted confidence in the AlphaFold2 model.

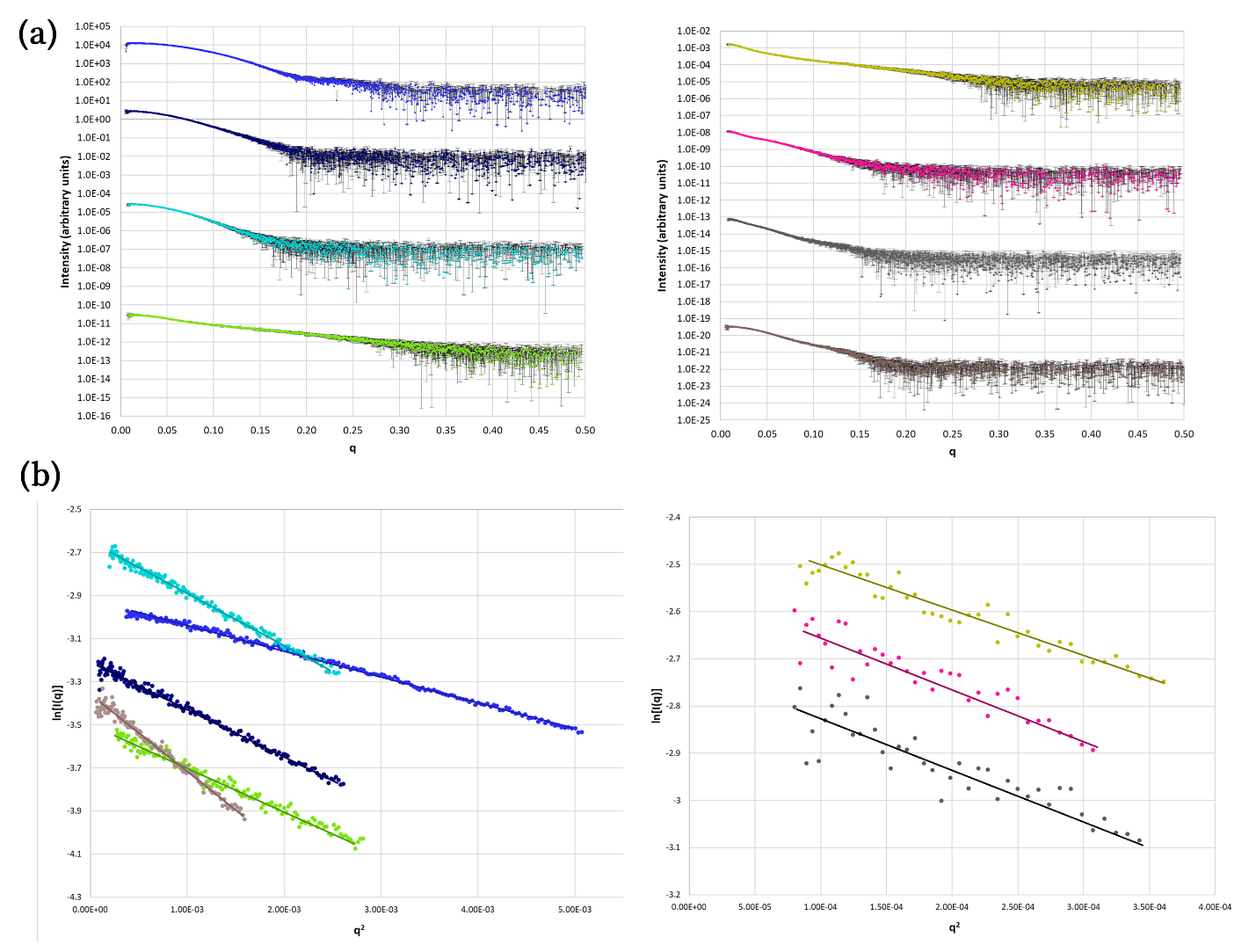

Supplementary Figure 4 – SAXS Scattering Curves and Guinier Plots for All Protein Constructs. **(a)** The scattering profiles of the *T. ramosa* IgAP NTD (light blue), MD (dark blue), MD+CTD1 (cyan), CTD2 (green), CTD1-4 (yellow), MD+CTD1-4 (pink), full-length (grey), and NTD+MD (brown) are shown here arbitrarily offset in I(q) to aid with visual comparison. Data are presented as mean values ± one standard deviation across 15 azimuthally averaged detector images. **(b)** Guinier analyses for the same SAXS data are shown with the line of best fit overlayed onto the processed data.

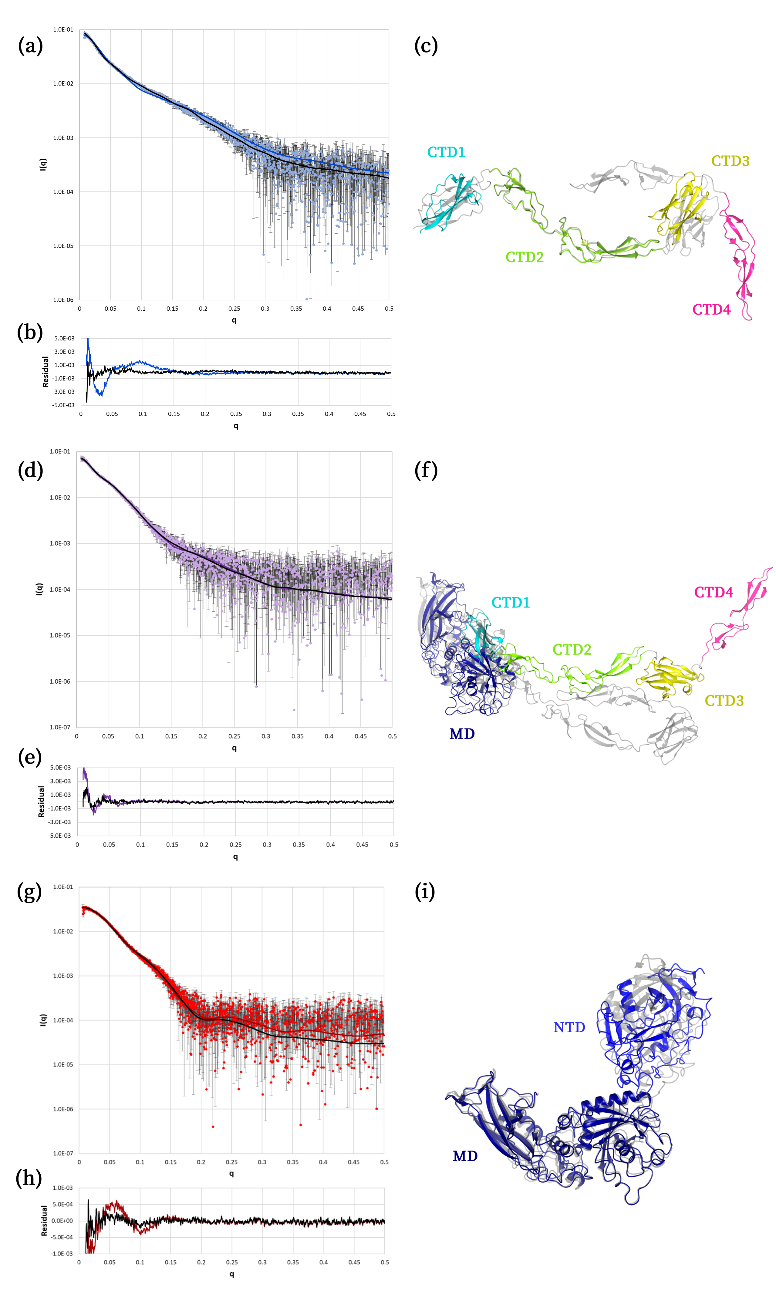

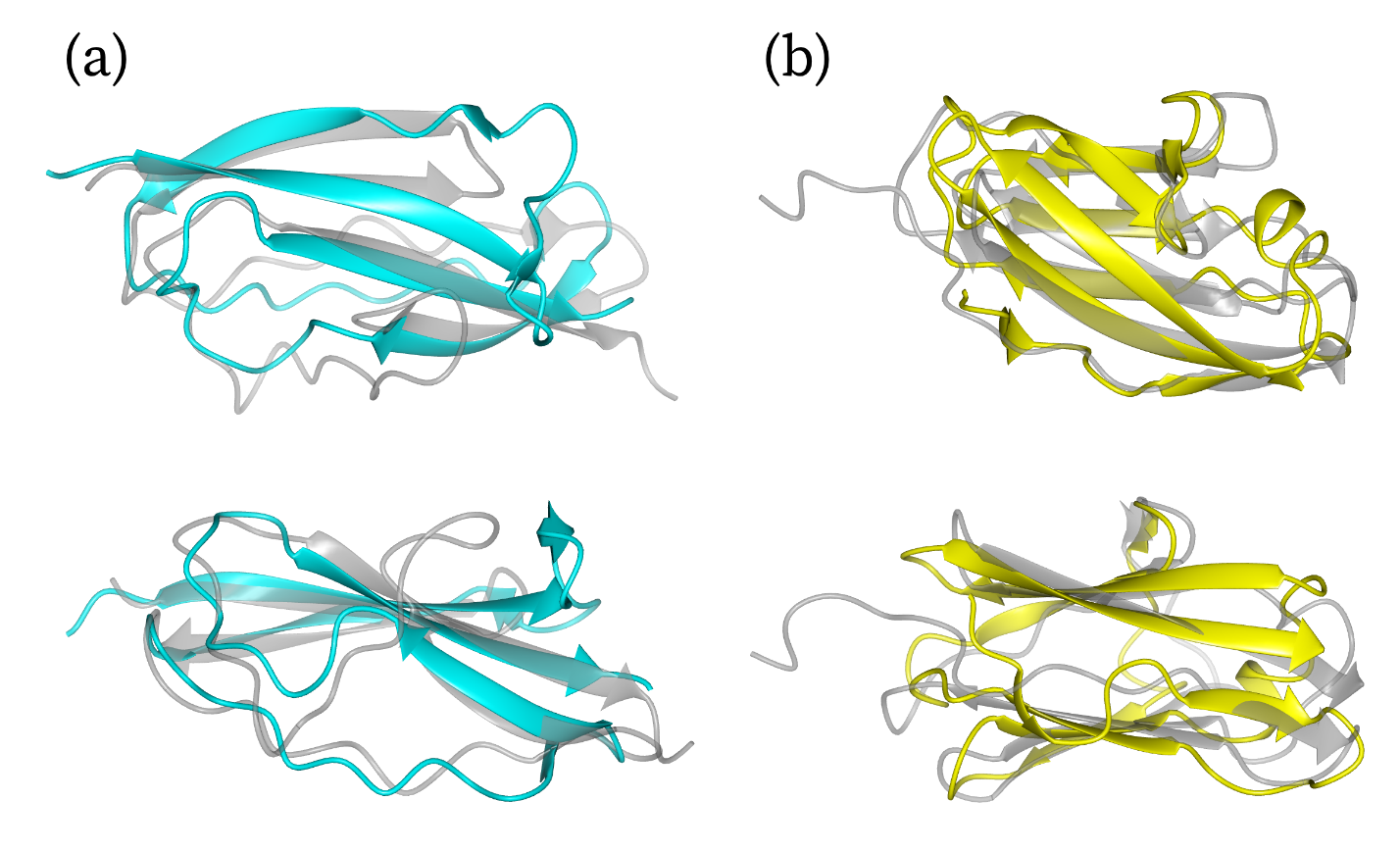

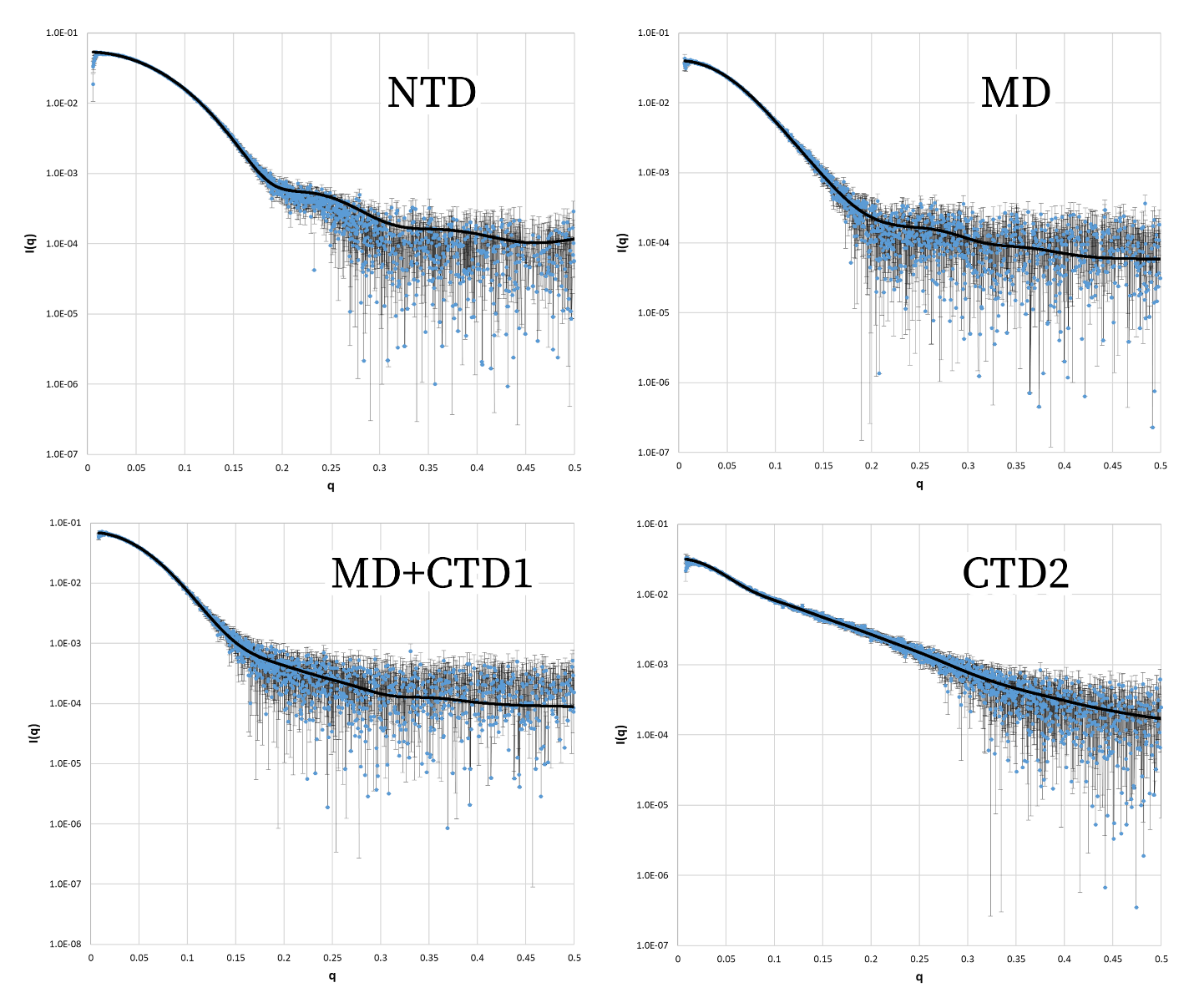

Supplementary Figure 5 – Simulated Scatter of CORAL-Relaxed AlphaFold2 Models Suggest the Enzyme Exists in an Extended Conformation. FoXS was used to generate the simulated solution scattering intensities of the CORAL-relaxed (black) and AlphaFold2-predicted **(a & c)** CTD1-4 (dark blue), and **(d & f)** MD+CTD1-4 (dark purple), and **(g & i)** NTD+MD (red) models. **(a,d,g)** Theoretical scattering curves from FoXS for each protein were compared to real data and **(b,e,h)** residuals were calculated for each measured angle. Data in the scattering curves are presented as mean values ± one standard deviation across 15 azimuthally averaged detector images. The simulated profiles for the CORAL-relaxed models had better data-to-model fits relative to their AlphaFold2-predicted models. The FoXS χ2 values of the CORAL-relaxed and AlphaFold2-predicted fits, respectively, were 0.79 and 2.98 for CTD1-4, 0.74 and 0.95 for MD+CTD1-4, and 1.00 and 1.60 for NTD+MD **(c,f,i).** The CORAL-relaxed models (colored) were always in an extended conformation relative to the AlphaFold2 model (transparent grey).

Supplementary Figure 6 – **CTD1 and CTD3 Are Structurally Homologous to Known Folds of Periscope Domains**. **(a)** CTD1 (cyan) is structurally homologous (2.07Å Cα-RMSD) to the B-repeat of *Listeria monocytogenes* InlB (PDB 2Y5P, chain A). **(b)** CTD3 (yellow) is structurally homologous (2.01Å Cα-RMSD) to the I10 domain of human titin (PDB 5JDJ, chain D). The AlphaFold2 models of the CTDs are superimposed onto their structural homologues (grey) and shown at two perpendicular viewing angles.

Supplementary Figure 7 – Simulated Scatter of AlphaFold2 Models Fit Their Scattering Curves. FoXS was used to generate the simulated solution scattering intensity (black line) of the NTD, MD+CTD1, and CTD2 AlphaFold2 models with manually added zinc ions where applicable. Data in the scattering curves are presented as mean values ± one standard deviation across 15 azimuthally averaged detector images. All simulated profiles showed good data-to-model fits, with FoXS χ^2^ values of 1.94, 1.14, 0.74, 0.90 for NTD, MD, MD+CTD1, and CTD2, respectively.

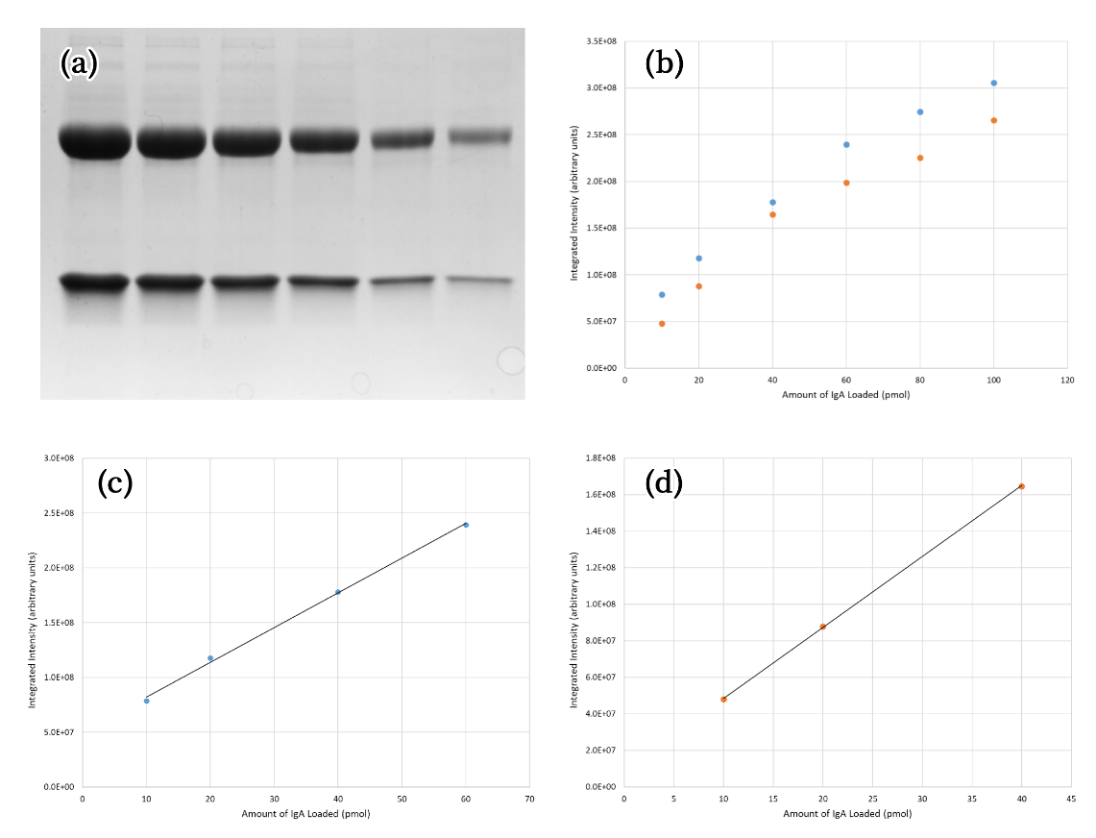

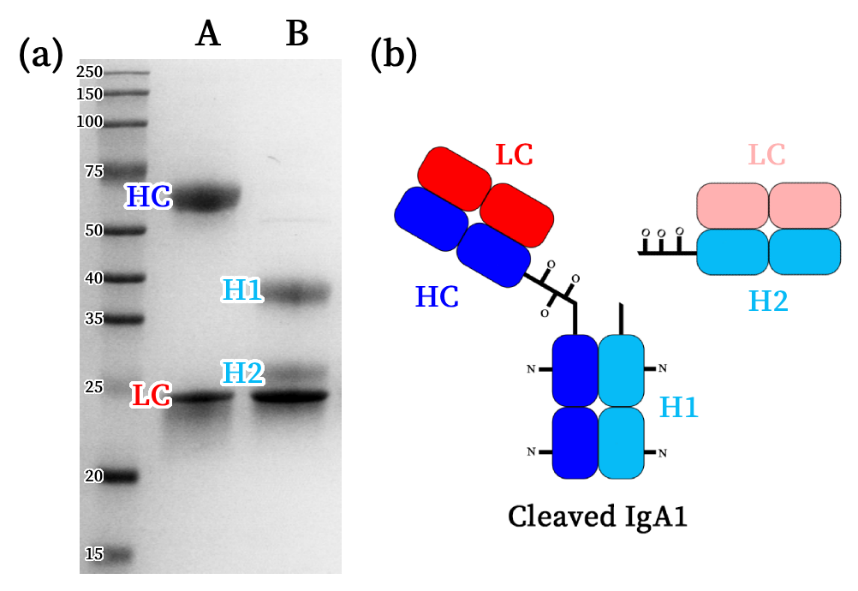

Supplementary Figure 9 – Testing Linearity of the Gel-Based Kinetic Assay Using a Substrate Standard Curve. **(a)** Intact IgA1 was titrated from 10-100 pmol and loaded on an SDS-PAGE gel. **(b)** The bands for the IgA1 heavy chain (blue) and light chain (orange) were integrated and their intensities were plotted as a function of amount of IgA loaded. **(c)** The IgA1 heavy chain intensity remains linear (R^2^ = 0.998) up to 60 pmol while **(d)** the intensity for the light chain remains linear (R^2^ = 0.999) up to 40 pmol.

Supplementary Figure 8 – Cleavage Pattern of IgA1 on an SDS-PAGE Gel. **(a)** The SDS-PAGE banding pattern of intact IgA1 (A) and IgA1 cleaved by the *T. ramosa* IgAP (B) are distinct and can be separately analyzed via densitometry. **(B)** The intact heavy chain (HC; blue) is split into two fragments following proteolysis while the light chain (LC; red and light red) remains unchanged. The HC fragment associated with the Fc (H1; light blue) is larger compared to the fragment associated with the Fab (H2; light blue).

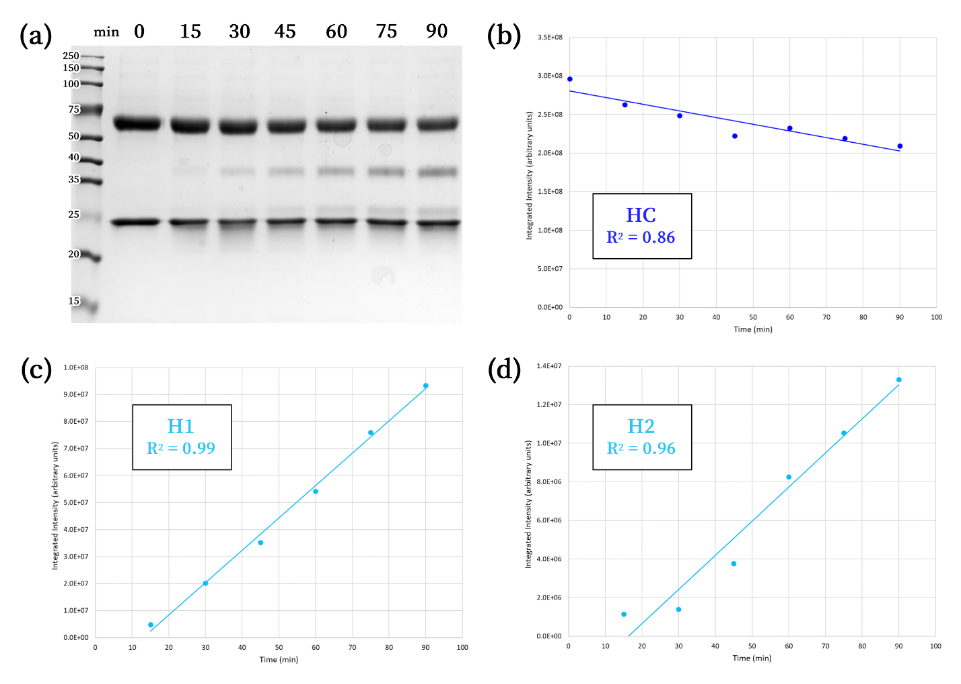

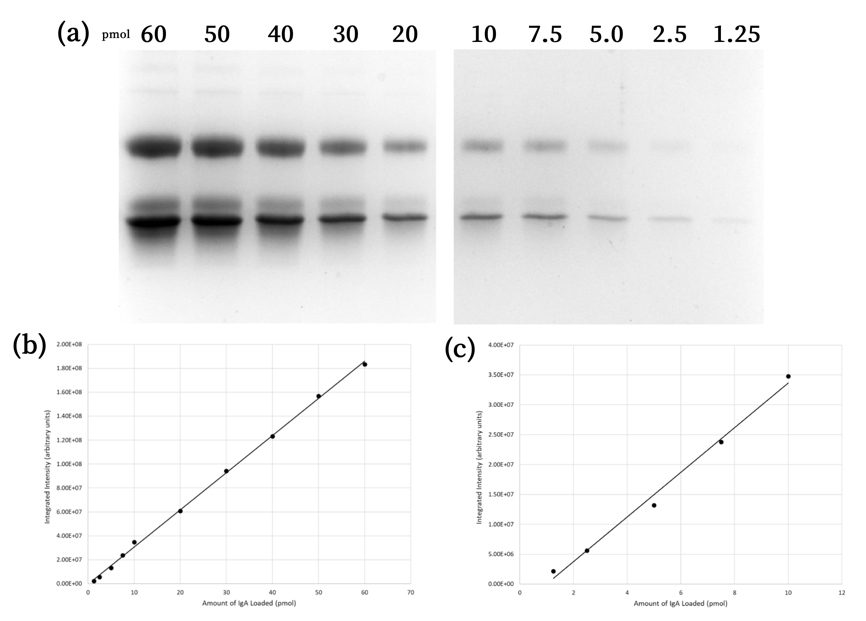

Supplementary Figure 11 – The Cleaved Heavy-Chain Fragment Associated with the Fc Provided the Most Reliable Readout for the Gel-Based Assay. **(a)** 0.5 nM *T. ramosa* IgAP MD was incubated with 3 μM IgA1 at 37 ^o^C and sampled at 15-minute intervals. The integrated band intensities for the **(b)** heavy chain, **(c)** H1 fragment, and **(c)** H2 fragment were plotted with respect to time.

Supplementary Figure 10 – Testing Linearity of the Gel-Based Kinetic Assay Using a Product Standard Curve. **(a)** IgA1 that was fully cleaved with *T. ramosa* IgAP MD was titrated from 1.25-60 pmol and loaded on an SDS-PAGE gel. **(b)** The band for the IgA1 H1 fragment was integrated and its intensities were plotted as a function of amount of IgA loaded. This standard curve shows a strong linear correlation (R^2^ = 0.999) for all amounts loaded. **(c)** The same data for 1.25-10 pmol cleaved IgA1 was replotted in a separate standard curve that is more representative of intensities that will be measured within the initial-rate region of the progress curve. This curve also shows a strong linear correlation (R^2^ = 0.992) for all amounts loaded.

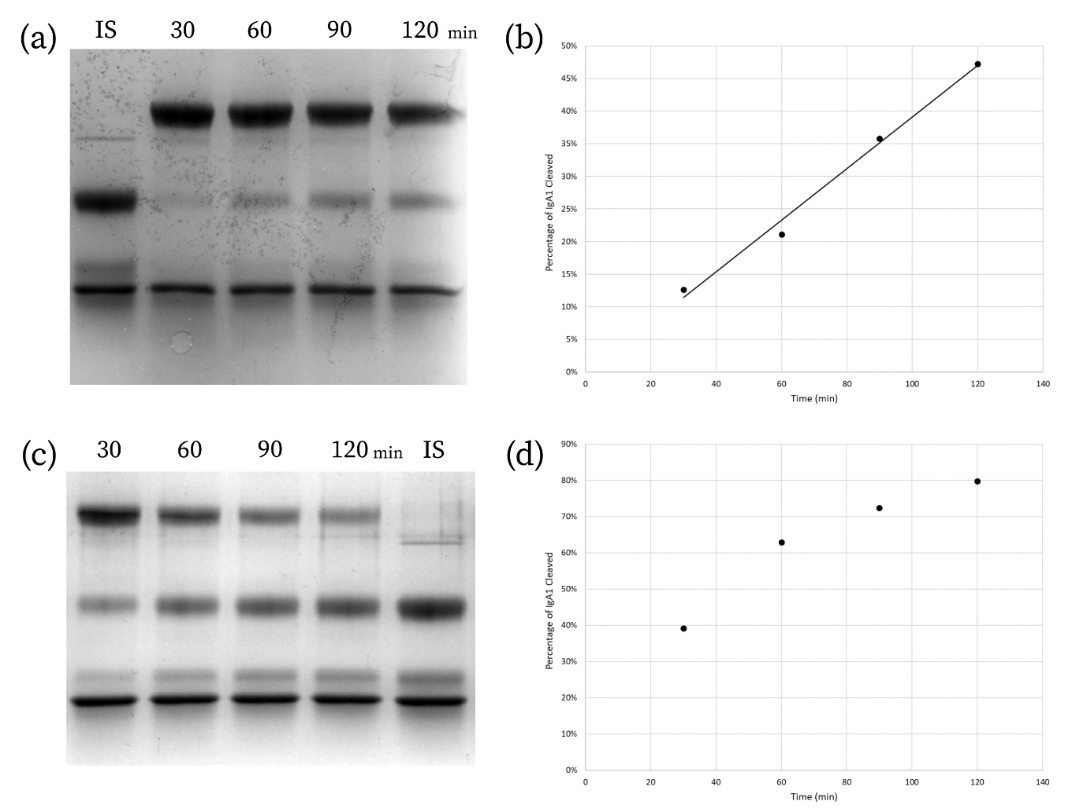

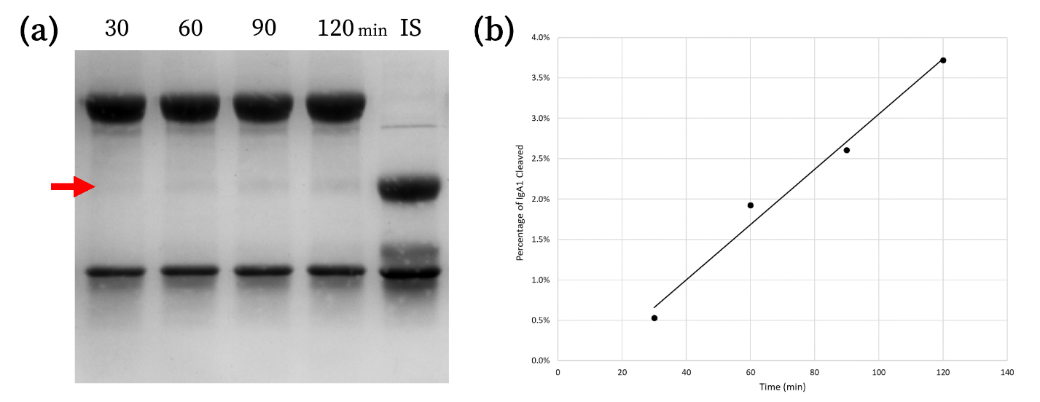

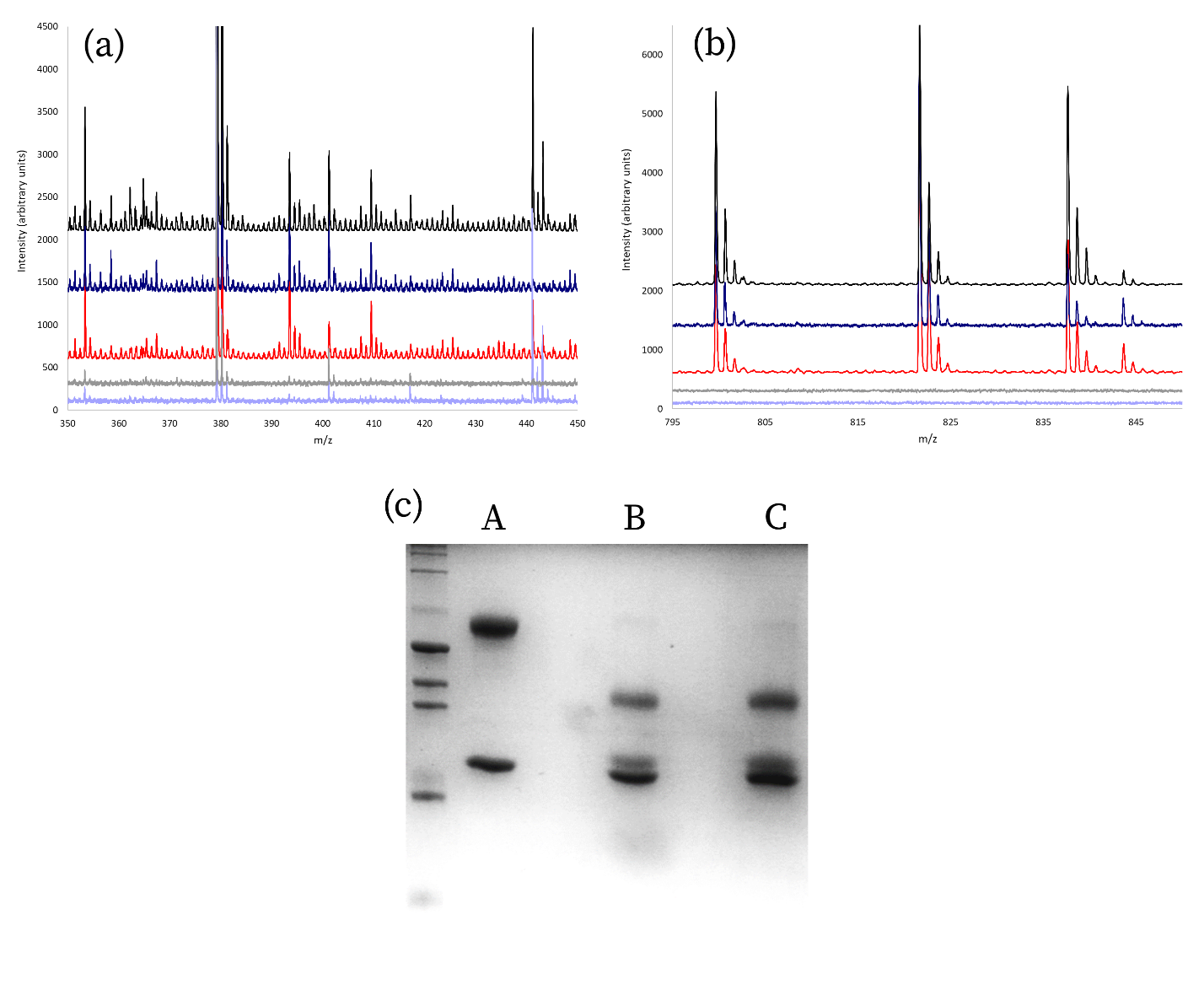

Supplementary Figure 12 – The Gel-Based Assay is Capable of Determining Initial Rates Under Michaelis-Menten Conditions. **(a)** 0.75 nM *T. ramosa* IgAP NTD+MD was incubated with 50 μM IgA1 at 37 ^o^C and sampled at 30-minute intervals. These samples were run alongside a fully cleaved 2 μM IgA1 internal standard (IS) to determine the percentage of IgA1 cleaved in each lane. The H1 band is indicated by the red arrow. **(b)** The integrated band intensities for the H1 fragment were converted to the percentage of IgA1 cleaved and plotted with respect to time. Under initial-rate conditions (<5% substrate consumed), the H1 band intensity shows a strong linear correlation with time (R^2^ = 0.98).

Supplementary Figure 13 – The *Thomasclavelia ramosa* Immunoglobulin A Protease is Robust Against Product Inhibition. **(a)** 1.5 nM full-length *T. ramosa* IgAP was incubated with 1 μM IgA1 at 37 ^o^C and sampled at 30-minute intervals. These samples were run alongside a fully cleaved 2 μM IgA1 internal standard (IS) to determine the percentage of IgA1 cleaved in each lane. **(b)** The integrated band intensities for the H1 fragment were converted to the percentage of IgA1 cleaved and plotted with respect to time. The H1 band intensity shows a strong linear correlation with time (R^2^ = 0.99) despite the reaction progressing significantly past what is normally considered to be the initial-rate range. **(c)** 0.5 nM *T. ramosa* IgAP MD was incubated with 1 μM IgA1 at 37 ^o^C and sampled at 30-minute intervals. **(d)** Unlike (a) and (b), the reaction progressed too far and is slowing down due to effects of product inhibition and/or substrate depletion.

Supplementary Figure 14 – **The *T. ramosa* IgAP MD Does Not Cleave the IgA1 Hinge Peptide In Isolation.** MALDI-ToF mass spectrometry was used to detect the masses of intact (798.95 g/mol) and cleaved peptide (402.5 and 414.5 g/mol) after incubation with various *T. ramosa* IgAP enzyme constructs. **(a)** No peaks corresponding to the cleaved peptide were seen in the samples when the peptide was mixed with the full-length enzyme (black) or the MD (dark blue). All other peaks could be accounted for in the enzyme controls (MD – light blue; full-length – grey) or peptide-only control (red). **(b)** Peaks corresponding to the intact peptide were seen in all samples but the enzyme controls. The intact peptide also showed peaks in complex with Na^+^ or K^+^. **(d)** 2 μM IgA1 (A) was incubated with ~1.3 nM MD (B) or full-length enzyme (C) at the same molar ratio of substrate to enzyme and identical reaction conditions as the mass-spectrometry samples. All of the IgA1 was cleaved under these conditions.

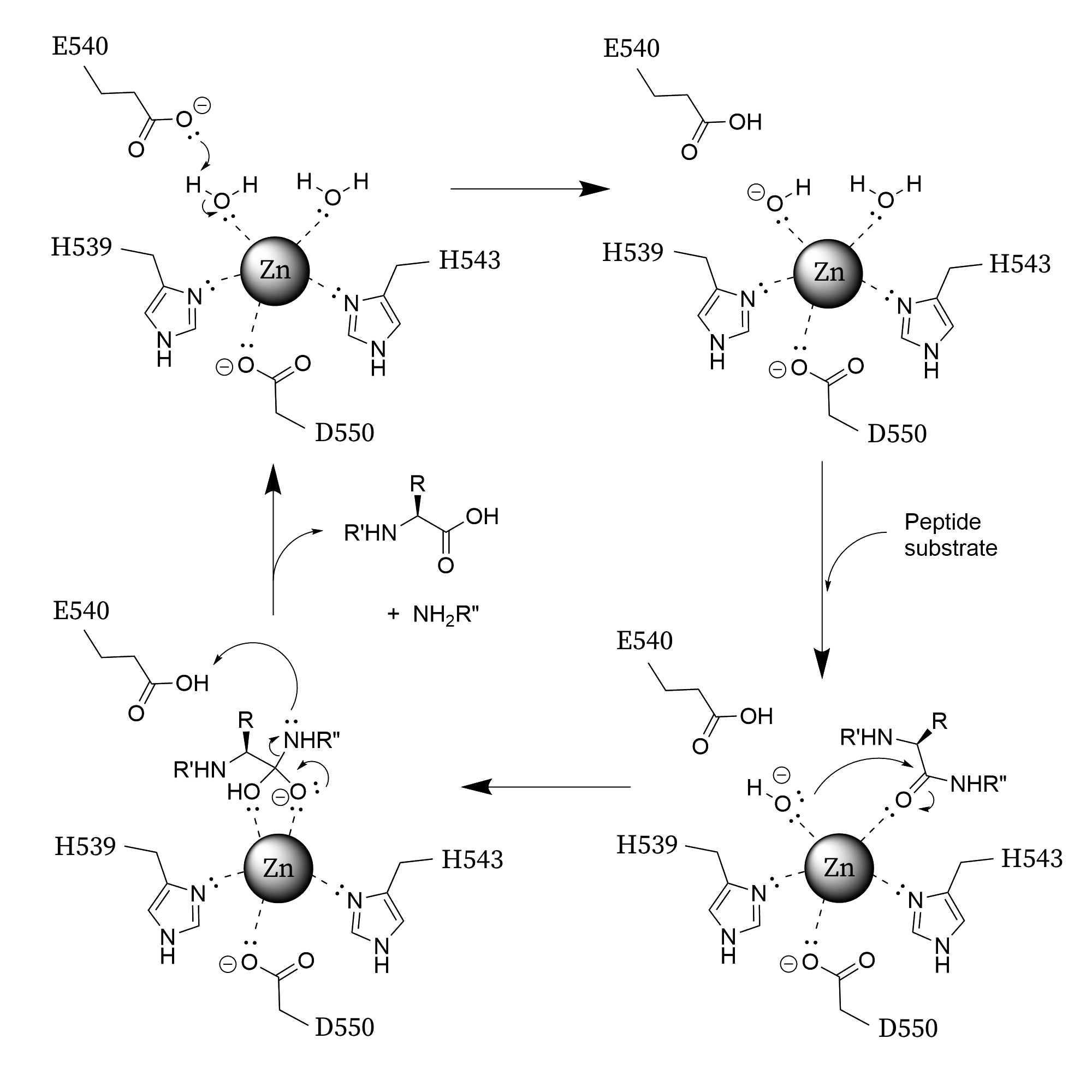

Supplementary Figure 15 – Catalytic Mechanism of Zinc Metalloproteases. The conserved chemical mechanism used by zinc metalloproteases is shown as a four-step cycle. R represents the side chain of the residue with the scissile peptide bond. For the *T. ramosa* IgAP, this is a proline side chain. R’ represents the portion of the immunoglobulin molecule found N-terminal to the scissile residue (Fab). R” represents the portion of the immunoglobulin molecule found C-terminal to the scissile residue (Fc). **(top left)** The active-site zinc is coordinated by H539, H543, D550, and two water molecules. **(top right)** The water molecule closest to the catalytic base, E540, is deprotonated to form the nucleophilic zinc-hydroxide complex. **(bottom right)** The peptide substrate binds to the active-site groove with the carbonyl of the scissile bond being stabilized by the zinc ion. The nucleophilic hydroxide then attacks the peptide bond. **(bottom left)** A tetrahedral intermediate is formed before breaking down into the two proteolysis products.

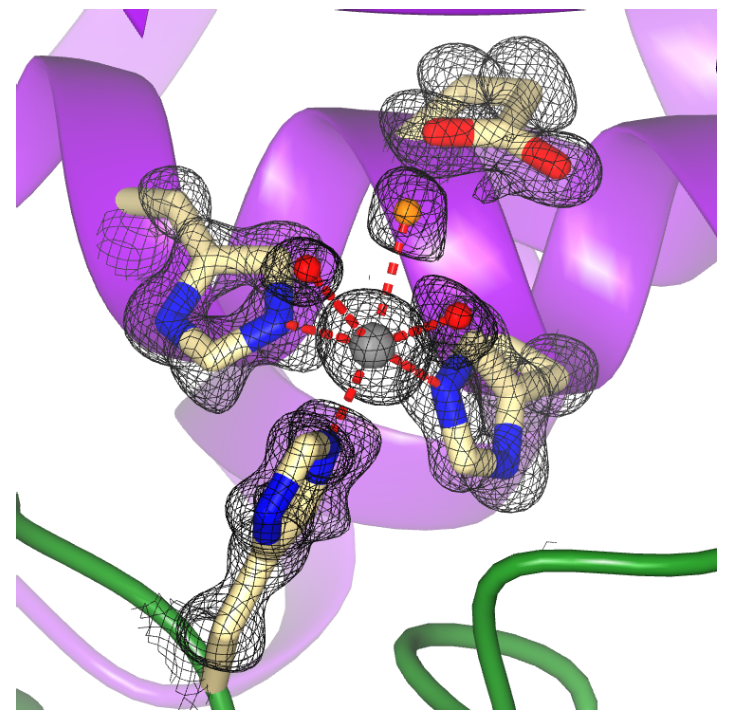

Supplementary Figure 16 – **Crystal Structure of the Human Matrix Metalloprotease-12 Active Site**. Compared to the holoenzyme structures of human matrix metalloprotease-12 (PDB 2OXU; black 2F_o_-F_c_ density contoured at 1.5 σ), the structure of the MD+CTD1 holoenzyme (*Fig. 6*) is conformationally heterogeneous at the active site.

##### Supplementary Table 1 – Small-Angle X-ray Scattering Data Details and Statistics.

| Protein construct | NTD | MD | MD+CTD1 | CTD2 |
| --- | --- | --- | --- | --- |
| SASBDB Entry | *SASDWH3* | *SASDWJ3* | *SASDWK3* | *SASDWL3* |
| Concentration (mg/mL) | 3.16 | 1.83 | 1.60 | 3.0 |
| q-measurement range | 0.00584 – 0.50081 | 0.00584 – 0.50081 | 0.00821 – 0.51346 | 0.00821 – 0.51346 |
| Guinier analysis |  |  |  |  |
| I(0) (arbitrary units) | 0.0541 | 0.0405 | 0.0705 | 0.0305 |
| R_g_ (Å) | 18.94 | 25.66 | 26.67 | 24.99 |
| q-range (Å^-1^) | 0.0193 – 0.0712 | 0.0091 – 0.0511 | 0.0141 – 0.0506 | 0.0165 – 0.0531 |
| q_min_*R_g_ | 0.365 | 0.233 | 0.376 | 0.389 |
| q_max_*R_g_ | 1.349 | 1.311 | 1.349 | 1.007 |
| R^2^ fit | 0.996 | 0.989 | 0.992 | 0.916 |
| P(r) analysis |  |  |  |  |
| I(0) (arbitrary units) | 0.0541 | 0.0407 | 0.0707 | 0.0312 |
| R_g_ (Å) | 18.91 | 26.12 | 26.97 | 26.93 |
| D_max_ (Å) | 60 | 88 | 92 | 96 |
| R^2^ fit | 0.951 | 0.882 | 0.653 | 0.690 |
| q-range (Å^-1^) | 0.0193 – 0.4010 | 0.0091 – 0.5008 | 0.0141 – 0.4135 | 0.0156 – 0.5135 |
| Volume analysis (kDa) |  |  |  |  |
| Theoretical | 31.63 | 54.78 | 62.53 | 13.59 |
| Porod volume (V_p_) | 34.0 | 54.5 | 65.7 | 9.1 |
| Correl. volume (V_c_) | 25.0 | 46.3 | 56.0 | 9.2 |

| Protein construct | NTD+MD | CTD1-4 | MD+CTD1-4 | FL |
| --- | --- | --- | --- | --- |
| SASBDB Entry | *SASDWM3* | *SASDWN3* | *SASDWP3* | *SASDWQ3* |
| Concentration (mg/mL) | 1.14 | 3.05 | 1.10 | 0.74 |
| q-measurement range | 0.00584 – 0.50081 | 0.00821 – 0.51346 | 0.00821 – 0.51346 | 0.00821 – 0.51346 |
| Guinier analysis |  |  |  |  |
| I(0) (arbitrary units) | 0.0350 | 0.0900 | 0.0791 | 0.0656 |
| R_g_ (Å) | 32.91 | 53.23 | 57.48 | 55.28 |
| q-range (Å^-1^) | 0.0078 – 0.0400 | 0.0092 – 0.0190 | 0.0089 – 0.0175 | 0.0089 – 0.0185 |
| q_min_*R_g_ | 0.258 | 0.490 | 0.514 | 0.494 |
| q_max_*R_g_ | 1.305 | 1.011 | 1.007 | 1.023 |
| R^2^ fit | 0.978 | 0.931 | 0.862 | 0.808 |
| P(r) analysis |  |  |  |  |
| I(0) (arbitrary units) | 0.0352 | 0.0917 | 0.0824 | 0.0680 |
| R_g_ (Å) | 33.48 | 57.09 | 65.07 | 62.63 |
| D_max_ (Å) | 108 | 215 | 240 | 245 |
| R^2^ fit | 0.944 | 0.635 | 0.672 | 0.619 |
| q-range (Å^-1^) | 0.0078 – 0.3009 | 0.0092 – 0.5135 | 0.0111 – 0.5135 | 0.0089 – 0.5135 |
| Volume analysis (kDa) |  |  |  |  |
| Theoretical | 88.02 | 39.27 | 95.00 | 130.85 |
| Porod volume (V_p_) | 82.7 | 74.3 | 125.6 | 147.3 |
| Correl. volume (V_c_) | 71.1 | 30.7 | 74.9 | 98.7 |

##### Supplementary Table 2 – Michaelis-Menten Parameters for Various Domain Constructs.^a^

| Protein construct | MD | MD+CTD1 | NTD+MD | FL |
| --- | --- | --- | --- | --- |
| k_cat_/K_M_ (M^-1^s^-1^) | 2x10^-7^ ± 2x10^-8^ | 2x10^-7^ ± 5x10^-9^ | 8x10^-8^ ± 9x10^-9^ | 4x10^-8^ ± 3x10^-9^ |
| k_cat_ (s^-1^) | 1.4 ± 0.04 | 1.4 ± 0.02 | 0.70 ± 0.03 | 0.62 ± 0.02 |
| K_M_ (μM) | 7.2 ± 0.6 | 9.2 ± 0.3 | 8.9 ± 0.9 | 15 ± 1 |

^a^ These data represent kinetic parameters obtained from best-fits of the Michaelis-Menten equation to the curves shown in *Fig. 4*. The error values only represent variation in the mathematical fit and not in the experimental methodology itself, due to the impracticalities of repeating the assay in replicates. Thus, the error values likely underestimate the true error of the parameters.

Supplementary Table 3 – Data Collection and Refinement Statistics. A summary of data and refinement statistics for all crystallographic datasets. Statistics for the highest-resolution bin are in parentheses.

| Description of structure | Protease domain  (native dataset; PDB 9EKK) | Protease domain  (above Zn edge) | Protease domain  (below Zn edge) |
| --- | --- | --- | --- |
| Wavelength (Å) | 1.1808 | 1.2705 | 1.2970 |
| Resolution range | 91.45 – 1.65  (1.67 – 1.65) | 91.35 – 1.85  (1.88 – 1.85) | 91.34 – 1.85  (1.88 – 1.85) |
| Space group | P2_1_ | P2_1_ | P2_1_ |
| Unit cell (Å) | 70.69 91.27 86.86  β = 91.09^°^ | 70.66 91.23 86.81  β = 91.10^°^ | 70.69 91.27 86.86  β = 91.0° |
| Total reflections | 803949 (21824) | 591061 (17258) | 589909 (17004) |
| Unique reflections | 122442 (3852) | 88545 (3003) | 88345 (2963) |
| Multiplicity | 6.57 (5.67) | 6.68 (5.75) | 6.68 (5.74) |
| Completeness (%) | 92.39 (58.61) | 94.24 (64.55) | 94.04 (63.75) |
| Mean I/σ(I) | 14.4 (0.6) | 12.0 (0.7) | 11.9 (0.6) |
| Wilson B-factor | 21.10 | 24.65 | 24.42 |
| R_merge_ | 0.083 (1.410) | 0.120 (1.469) | 0.119 (1.454) |
| R_meas_ | 0.090 (1.554) | 0.131 (1.617) | 0.130 (1.601) |
| R_pim_ | 0.035 (0.639) | 0.050 (0.661) | 0.050 (0.653) |
| CC_1/2_ | 0.997 (0.591) | 0.995 (0.645) | 0.995 (0.661) |
| Number of reflections used in refinement | 115075 (2329) |  |  |
| Number of reflections used for R_free_ | 6315 (150) |  |  |
| R_work_ | 0.1959 (0.3069) |  |  |
| R_free_ | 0.2307 (0.3442) |  |  |
| Molecules per asymmetric unit | 2 |  |  |
| Number of atoms | 10205 |  |  |
| Protein | 8931 |  |  |
| Ligands | 10 |  |  |
| Water | 1264 |  |  |
| Average B-factors (Å^2^) |  |  |  |
| Overall | 27.36 |  |  |
| Protein | 26.36 |  |  |
| Ligands | 37.39 |  |  |
| Water | 34.36 |  |  |
| Bond RMSD (Å) | 0.006 |  |  |
| Angle RMSD (^o^) | 0.80 |  |  |
| Ramachandran (%) |  |  |  |
| Favored | 97.52 |  |  |
| Allowed | 2.48 |  |  |
| Outliers | 0 |  |  |
| Rotamer outliers (%) | 0.10 |  |  |
| Clashscore (percentile) | 2.62 (99^th^) |  |  |
| MolProbity score (percentile) | 1.15 (99^th^) |  |  |

Supplementary Table 3 **Continued –** *Data Collection and Refinement Statistics*.

| Description of structure | Metal-chelated  protease domain  (native dataset; PDB 9EKM) | Metal-chelated  protease domain  (above Zn edge) | Metal-chelated  protease domain  (below Zn edge) |
| --- | --- | --- | --- |
| Wavelength (Å) | 1.1808 | 1.2651 | 1.3051 |
| Resolution range | 91.70 – 1.50  (1.53 – 1.50) | 91.71 – 1.80  (1.83 – 1.80) | 91.72 – 1.85  (1.88 – 1.85) |
| Space group | P2_1_ | P2_1_ | P2_1_ |
| Unit cell (Å) | 70.74 91.55 86.85  β = 90.96^o^ | 70.80 91.63 86.90  β = 90.96^o^ | 70.83 91.66 86.94  β = 90.96^o^ |
| Total reflections | 1111333 (33717) | 620884 (14669) | 594648 (17082) |
| Unique reflections | 174570 (7224) | 98765 (3397) | 93148 (3706) |
| Multiplicity | 6.37 (4.67) | 6.29 (4.32) | 6.38 (4.61) |
| Completeness (%) | 98.91 (82.51) | 96.22 (66.32) | 98.46 (79.07) |
| Mean I/σ(I) | 6.3 (0.2) | 11.1 (0.6) | 11.4 (0.8) |
| Wilson B-factor | 18.31 | 23.53 | 24.31 |
| R_merge_ | 0.089 (1.174) | 0.103 (1.243) | 0.103 (0.997) |
| R_meas_ | 0.097 (1.326) | 0.112 (1.418) | 0.112 (1.128) |
| R_pim_ | 0.037 (0.594) | 0.044 (0.658) | 0.043 (0.509) |
| CC_1/2_ | 0.998 (0.499) | 0.996 (0.478) | 0.996 (0.604) |
| Number of reflections used in refinement | 165774 (4365) |  |  |
| Number of reflections used for R_free_ | 8718 (240) |  |  |
| R_work_ | 0.1388 (0.2845) |  |  |
| R_free_ | 0.1810 (0.3214) |  |  |
| Molecules per asymmetric unit | 2 |  |  |
| Number of atoms | 10467 |  |  |
| Protein | 9009 |  |  |
| Ligands | 6 |  |  |
| Water | 1452 |  |  |
| Average B-factors (Å^2^) |  |  |  |
| Overall | 22.36 |  |  |
| Protein | 20.54 |  |  |
| Ligands | 30.50 |  |  |
| Water | 33.61 |  |  |
| Bond RMSD (Å) | 0.006 |  |  |
| Angle RMSD (^o^) | 0.79 |  |  |
| Ramachandran (%) |  |  |  |
| Favored | 98.07 |  |  |
| Allowed | 1.93 |  |  |
| Outliers | 0 |  |  |
| Rotamer outliers (%) | 0.10 |  |  |
| Clashscore (percentile) | 2.37 (99^th^) |  |  |
| MolProbity score (percentile) | 1.02 (99^th^) |  |  |

Supplementary Table 3 **Continued –** *Data Collection and Refinement Statistics*.

| Description of structure | Metal-chelated protease domain with 1 mM ZnCl_2_  (native dataset; PDB 9EKN) | Metal-chelated protease domain with 1 mM ZnCl_2_  (above Zn edge) | Metal-chelated protease domain with 1 mM ZnCl_2_  (below Zn edge) |
| --- | --- | --- | --- |
| Wavelength (Å) | 1.1808 | 1.2651 | 1.3051 |
| Resolution range | 92.01 – 1.78  (1.81 – 1.78) | 92.01 – 2.05  (2.09 – 2.05) | 92.01 – 2.05  (2.09 – 2.05) |
| Space group | P2_1_2_1_2_1_ | P2_1_2_1_2_1_ | P2_1_2_1_2_1_ |
| Unit cell (Å) | 71.13 85.64 92.12 | 71.25 85.75 92.22 | 71.24 85.76 92.23 |
| Total reflections | 689077 (31395) | 458850 (23381) | 458158 (23257) |
| Unique reflections | 54207 (2637) | 35820 (1789) | 35826 (1788) |
| Multiplicity | 12.71 (11.91) | 12.81 (13.07) | 12.79 (13.01) |
| Completeness (%) | 100 (100) | 100 (100) | 100 (100) |
| Mean I/σ(I) | 11.0 (0.3) | 10.1 (0.6) | 9.9 (0.6) |
| Wilson B-factor | 28.35 | 36.84 | 36.93 |
| R_merge_ | 0.102 (2.166) | 0.170 (2.430) | 0.174 (2.604) |
| R_meas_ | 0.107 (2.264) | 0.177 (2.529) | 0.182 (2.710) |
| R_pim_ | 0.030 (0.654) | 0.049 (0.697) | 0.050 (0.748) |
| CC_1/2_ | 0.999 (0.586) | 0.996 (0.538) | 0.998 (0.501) |
| Number of reflections used in refinement | 51393 (2605) |  |  |
| Number of reflections used for R_free_ | 2713 (143) |  |  |
| R_work_ | 0.1707 (0.3448) |  |  |
| R_free_ | 0.2072 (0.3737) |  |  |
| Molecules per asymmetric unit | 1 |  |  |
| Number of atoms | 4999 |  |  |
| Protein | 4502 |  |  |
| Ligands | 13 |  |  |
| Water | 490 |  |  |
| Average B-factors (Å^2^) |  |  |  |
| Overall | 36.69 |  |  |
| Protein | 36.18 |  |  |
| Ligands | 45.63 |  |  |
| Water | 41.18 |  |  |
| Bond RMSD (Å) | 0.006 |  |  |
| Angle RMSD (^o^) | 0.73 |  |  |
| Ramachandran (%) |  |  |  |
| Favored | 96.88 |  |  |
| Allowed | 3.12 |  |  |
| Outliers | 0 |  |  |
| Rotamer outliers (%) | 0.20 |  |  |
| Clashscore (percentile) | 2.71 (99^th^) |  |  |
| MolProbity score (percentile) | 1.25 (99^th^) |  |  |

**Supplementary Table 4 – Vector and Open-Reading Frame Sequences.** The full sequences of the expression vectors and their associated construct as described in the Online Methods section are presented. The primary sequences of the proteins after purification (*i.e.,* after affinity tag cleavage) are also detailed below.

| Protein construct | Full Vector Sequence | Purified Protein Sequence |
| --- | --- | --- |
| NTD | gatggtgtccgggatctcgacgctctcccttatgcgactcctgcattaggaagcagcccagtagtaggttgaggccgttgagcaccgccgccgcaaggaatggtgcatgcaaggagatggcgcccaacagtcccccggccacggggcctgccaccatacccacgccgaaacaagcgctcatgagcccgaagtggcgagcccgatcttccccatcggtgatgtcggcgatataggcgccagcaaccgcacctgtggcgccggtgatgccggccacgatgcgtccggcgtagaggatcgagatctcgatcccgcgaaattaatacgactcactataggggaattgtgagcggataacaattcccctctagaaataattttgtttaactttaagaaggagatataccatgggtcatcaccatcatcatcacgggtcggactcagaagtcaatcaagaagctaagccagaggtcaagccagaagtcaagcctgagactcacatcaatttaaaggtgtccgatggatcttcagagatcttcttcaagatcaaaaagaccactcctttaagaaggctgatggaagcgttcgctaaaagacagggtaaggaaatggactccttaagattcttgtacgacggtattagaattcaagctgatcaggcccctgaagatttggacatggaggataacgatattattgaggctcaccgcgaacagattggaggtgcttcaaaacccgacataaaggtaggagattacgttaaaatgggtgtttataataacgcgagcattctgtggcgttgcgtttctatcgataacaacggtccgttaatgctggcagacaagattgtggataccctcgcatacgacgcgaaaaccaacgacaactctaacagcaagtcccacagccgtagctacaaacgtgatgattatggctccaactactggaaggactcaaacatgcgtagctggctcaatagcacggctgcggaaggtaaagtagactggctgtgcggtaacccgcccaaagatggctatgttagcggtgtgggtgcgtacaacgaaaaagccggctttctgaacgcatttagcaagtctgaaatcgcagcaatgaaaaccgttacccaacgttctctggtttcgcatccggagtacaacaaaggtatcgttgacggcgacgccaattctgatctgttgtactacacagacatcagtgaggcggttgcgaactatgactctagctacttcgaaactactacggagaaggtgtttctgttggacgttaaacaagcaaatgccgtgtggaaaaacctgaagggctattacgtggcgtataacaacgacggtatggcgtggccgtattggctgcgtaccccggtgaccgactgtaaccacgatatgcgttatatcagttccagcggccaggtcggtcgttatgctccgtggtatagcgacctgggcgtgagaccagcattttatctggactctgagtattttgttactacgagcggctctggctcccaaagcagtccgtatatcggctccgcgccgaacaaacaagaagatgactacacctgactcgagcaccaccaccaccaccactgagatccggctgctaacaaagcccgaaaggaagctgagttggctgctgccaccgctgagcaataactagcataaccccttggggcctctaaacgggtcttgaggggttttttgctgaaaggaggaactatatccggattggcgaatgggacgcgccctgtagcggcgcattaagcgcggcgggtgtggtggttacgcgcagcgtgaccgctacacttgccagcgccctagcgcccgctcctttcgctttcttcccttcctttctcgccacgttcgccggctttccccgtcaagctctaaatcgggggctccctttagggttccgatttagtgctttacggcacctcgaccccaaaaaacttgattagggtgatggttcacgtagtgggccatcgccctgatagacggtttttcgccctttgacgttggagtccacgttctttaatagtggactcttgttccaaactggaacaacactcaaccctatctcggtctattcttttgatttataagggattttgccgatttcggcctattggttaaaaaatgagctgatttaacaaaaatttaacgcgaattttaacaaaatattaacgcttacaatttaggtggcacttttcggggaaatgtgcgcggaacccctatttgtttatttttctaaatacattcaaatatgtatccgctcatgaattaattcttagaaaaactcatcgagcatcaaatgaaactgcaatttattcatatcaggattatcaataccatatttttgaaaaagccgtttctgtaatgaaggagaaaactcaccgaggcagttccataggatggcaagatcctggtatcggtctgcgattccgactcgtccaacatcaatacaacctattaatttcccctcgtcaaaaataaggttatcaagtgagaaatcaccatgagtgacgactgaatccggtgagaatggcaaaagtttatgcatttctttccagacttgttcaacaggccagccattacgctcgtcatcaaaatcactcgcatcaaccaaaccgttattcattcgtgattgcgcctgagcgagacgaaatacgcgatcgctgttaaaaggacaattacaaacaggaatcgaatgcaaccggcgcaggaacactgccagcgcatcaacaatattttcacctgaatcaggatattcttctaatacctggaatgctgttttcccggggatcgcagtggtgagtaaccatgcatcatcaggagtacggataaaatgcttgatggtcggaagaggcataaattccgtcagccagtttagtctgaccatctcatctgtaacatcattggcaacgctacctttgccatgtttcagaaacaactctggcgcatcgggcttcccatacaatcgatagattgtcgcacctgattgcccgacattatcgcgagcccatttatacccatataaatcagcatccatgttggaatttaatcgcggcctagagcaagacgtttcccgttgaatatggctcataacaccccttgtattactgtttatgtaagcagacagttttattgttcatgaccaaaatcccttaacgtgagttttcgttccactgagcgtcagaccccgtagaaaagatcaaaggatcttcttgagatcctttttttctgcgcgtaatctgctgcttgcaaacaaaaaaaccaccgctaccagcggtggtttgtttgccggatcaagagctaccaactctttttccgaaggtaactggcttcagcagagcgcagataccaaatactgtccttctagtgtagccgtagttaggccaccacttcaagaactctgtagcaccgcctacatacctcgctctgctaatcctgttaccagtggctgctgccagtggcgataagtcgtgtcttaccgggttggactcaagacgatagttaccggataaggcgcagcggtcgggctgaacggggggttcgtgcacacagcccagcttggagcgaacgacctacaccgaactgagatacctacagcgtgagctatgagaaagcgccacgcttcccgaagggagaaaggcggacaggtatccggtaagcggcagggtcggaacaggagagcgcacgagggagcttccagggggaaacgcctggtatctttatagtcctgtcgggtttcgccacctctgacttgagcgtcgatttttgtgatgctcgtcaggggggcggagcctatggaaaaacgccagcaacgcggcctttttacggttcctggccttttgctggccttttgctcacatgttctttcctgcgttatcccctgattctgtggataaccgtattaccgcctttgagtgagctgataccgctcgccgcagccgaacgaccgagcgcagcgagtcagtgagcgaggaagcggaagagcgcctgatgcggtattttctccttacgcatctgtgcggtatttcacaccgcatatatggtgcactctcagtacaatctgctctgatgccgcatagttaagccagtatacactccgctatcgctacgtgactgggtcatggctgcgccccgacacccgccaacacccgctgacgcgccctgacgggcttgtctgctcccggcatccgcttacagacaagctgtgaccgtctccgggagctgcatgtgtcagaggttttcaccgtcatcaccgaaacgcgcgaggcagctgcggtaaagctcatcagcgtggtcgtgaagcgattcacagatgtctgcctgttcatccgcgtccagctcgttgagtttctccagaagcgttaatgtctggcttctgataaagcgggccatgttaagggcggttttttcctgtttggtcactgatgcctccgtgtaagggggatttctgttcatgggggtaatgataccgatgaaacgagagaggatgctcacgatacgggttactgatgatgaacatgcccggttactggaacgttgtgagggtaaacaactggcggtatggatgcggcgggaccagagaaaaatcactcagggtcaatgccagcgcttcgttaatacagatgtaggtgttccacagggtagccagcagcatcctgcgatgcagatccggaa | ASKPDIKVGDYVKMGVYNNASILWRCVSIDNNGPLMLADKIVDTLAYDAKTNDNSNSKSHSRSYKRDDYGSNYWKDSNMRSWLNSTAAEGKVDWLCGNPPKDGYVSGVGAYNEKAGFLNAFSKSEIAAMKTVTQRSLVSHPEYNKGIVDGDANSDLLYYTDISEAVANYDSSYFETTTEKVFLLDVKQANAVWKNLKGYYVAYNNDGMAWPYWLRTPVTDCNHDMRYISSSGQVGRYAPWYSDLGVRPAFYLDSEYFVTTSGSGSQSSPYIGSAPNKQEDDYT |
| NTD+MD | gatggtgtccgggatctcgacgctctcccttatgcgactcctgcattaggaagcagcccagtagtaggttgaggccgttgagcaccgccgccgcaaggaatggtgcatgcaaggagatggcgcccaacagtcccccggccacggggcctgccaccatacccacgccgaaacaagcgctcatgagcccgaagtggcgagcccgatcttccccatcggtgatgtcggcgatataggcgccagcaaccgcacctgtggcgccggtgatgccggccacgatgcgtccggcgtagaggatcgagatctcgatcccgcgaaattaatacgactcactataggggaattgtgagcggataacaattcccctctagaaataattttgtttaactttaagaaggagatataccatgggtcatcaccatcatcatcacgggtcggactcagaagtcaatcaagaagctaagccagaggtcaagccagaagtcaagcctgagactcacatcaatttaaaggtgtccgatggatcttcagagatcttcttcaagatcaaaaagaccactcctttaagaaggctgatggaagcgttcgctaaaagacagggtaaggaaatggactccttaagattcttgtacgacggtattagaattcaagctgatcaggcccctgaagatttggacatggaggataacgatattattgaggctcaccgcgaacagattggaggtgcttcaaaacccgacataaaggtaggagattacgttaaaatgggtgtttataataacgcgagcattctgtggcgttgcgtttctatcgataacaacggtccgttaatgctggcagacaagattgtggataccctcgcatacgacgcgaaaaccaacgacaactctaacagcaagtcccacagccgtagctacaaacgtgatgattatggctccaactactggaaggactcaaacatgcgtagctggctcaatagcacggctgcggaaggtaaagtagactggctgtgcggtaacccgcccaaagatggctatgttagcggtgtgggtgcgtacaacgaaaaagccggctttctgaacgcatttagcaagtctgaaatcgcagcaatgaaaaccgttacccaacgttctctggtttcgcatccggagtacaacaaaggtatcgttgacggcgacgccaattctgatctgttgtactacacagacatcagtgaggcggttgcgaactatgactctagctacttcgaaactactacggagaaggtgtttctgttggacgttaaacaagcaaatgccgtgtggaaaaacctgaagggctattacgtggcgtataacaacgacggtatggcgtggccgtattggctgcgtaccccggtgaccgactgtaaccacgatatgcgttatatcagttccagcggccaggtcggtcgttatgctccgtggtatagcgacctgggcgtgagaccagcattttatctggactctgagtattttgttactacgagcggctctggctcccaaagcagtccgtatatcggctccgcgccgaacaaacaagaagatgactacaccatttccgagccagcagaagacgccaacccggattggaacgttagtaccgagcagagcatccaactgaccctcggcccttggtattccaacgacggtaaatacagcaacccgaccatcccggtttacaccatccaaaaaacgcgttccgacaccgagaatatggtggtggtcgtgtgcggtgagggctacactaagagccagcagggtaaattcatcaacgacgtaaagcgcctgtggcaggatgctatgaagtatgagccgtaccgctcctacgcggatcgtttcaatgtgtacgctctgtgtaccgcgtcagagagcaccttcgacaacggtggttctacgttctttgatgtgatagttgacaaatacaactccccggttatttccaacaacctgcatggttcccaatggaagaaccacatttttgagcgttgtattggtccggagtttatcgaaaagattcatgatgcgcatattaaaaaaaagtgcgatccgaataccattccgtcggggagcgagtatgaaccgtactattacgtgcatgactacattgcgcagtttgctatggttgtgaataccaaaagtgacttcggcggtgcgtacaataaccgtgaatacggcttccattatttcatcagcccgagcgacagctatcgtgcaagcaaaaccttcgcgcacgaattcggtcacggcctgcttggcttgggtgatgaatactccaacgggtacctgttggatgacaaggaactgaagagcctgaacctgagcagcgtcgaggacccggaaaaaatcaagtggcgtcaattgctgggttttcgcaacacatatacgtgccgtaatgcgtacggctctaaaatgctggtttctagctacgagtgcatcatgcgcgacacgaattatcagttctgcgaagtttgccgcctgcaaggtttcaaacgcatgagccagttggtgaaggatgtcgacttgtatgtcgccaccccagaggttaaagaatacaccggcgcatacagcaagccgtccgactttaccgatttggaaacctccagctactacaactacacctataaccgcaatgaccgcttgcttagcggtaattcgaagagccgtttcaataccaatatgaacggtaaaaaaatcgagttgcgtaccgttatacagaatattagcgacaagaatgcgcgtcaactgaaatttaagatgtggattaaacacagcgatggtagcgtggcaacggactcgtccggcaacccgctgcagaccgtccagaccttcgatattccggtttggaacgataaggctaacttttggcctctgggtgcgttggaccacatcaagagcgacttcaacagcggtctgaagagctgtagcctgatttatcagattccgtctgatgcccaactaaagtccggcgacaccgttgcgttccaggtcctggacgaaaatggcaatgtgttggctgacgataacaccgaaacccagtgactcgagcaccaccaccaccaccactgagatccggctgctaacaaagcccgaaaggaagctgagttggctgctgccaccgctgagcaataactagcataaccccttggggcctctaaacgggtcttgaggggttttttgctgaaaggaggaactatatccggattggcgaatgggacgcgccctgtagcggcgcattaagcgcggcgggtgtggtggttacgcgcagcgtgaccgctacacttgccagcgccctagcgcccgctcctttcgctttcttcccttcctttctcgccacgttcgccggctttccccgtcaagctctaaatcgggggctccctttagggttccgatttagtgctttacggcacctcgaccccaaaaaacttgattagggtgatggttcacgtagtgggccatcgccctgatagacggtttttcgccctttgacgttggagtccacgttctttaatagtggactcttgttccaaactggaacaacactcaaccctatctcggtctattcttttgatttataagggattttgccgatttcggcctattggttaaaaaatgagctgatttaacaaaaatttaacgcgaattttaacaaaatattaacgcttacaatttaggtggcacttttcggggaaatgtgcgcggaacccctatttgtttatttttctaaatacattcaaatatgtatccgctcatgaattaattcttagaaaaactcatcgagcatcaaatgaaactgcaatttattcatatcaggattatcaataccatatttttgaaaaagccgtttctgtaatgaaggagaaaactcaccgaggcagttccataggatggcaagatcctggtatcggtctgcgattccgactcgtccaacatcaatacaacctattaatttcccctcgtcaaaaataaggttatcaagtgagaaatcaccatgagtgacgactgaatccggtgagaatggcaaaagtttatgcatttctttccagacttgttcaacaggccagccattacgctcgtcatcaaaatcactcgcatcaaccaaaccgttattcattcgtgattgcgcctgagcgagacgaaatacgcgatcgctgttaaaaggacaattacaaacaggaatcgaatgcaaccggcgcaggaacactgccagcgcatcaacaatattttcacctgaatcaggatattcttctaatacctggaatgctgttttcccggggatcgcagtggtgagtaaccatgcatcatcaggagtacggataaaatgcttgatggtcggaagaggcataaattccgtcagccagtttagtctgaccatctcatctgtaacatcattggcaacgctacctttgccatgtttcagaaacaactctggcgcatcgggcttcccatacaatcgatagattgtcgcacctgattgcccgacattatcgcgagcccatttatacccatataaatcagcatccatgttggaatttaatcgcggcctagagcaagacgtttcccgttgaatatggctcataacaccccttgtattactgtttatgtaagcagacagttttattgttcatgaccaaaatcccttaacgtgagttttcgttccactgagcgtcagaccccgtagaaaagatcaaaggatcttcttgagatcctttttttctgcgcgtaatctgctgcttgcaaacaaaaaaaccaccgctaccagcggtggtttgtttgccggatcaagagctaccaactctttttccgaaggtaactggcttcagcagagcgcagataccaaatactgtccttctagtgtagccgtagttaggccaccacttcaagaactctgtagcaccgcctacatacctcgctctgctaatcctgttaccagtggctgctgccagtggcgataagtcgtgtcttaccgggttggactcaagacgatagttaccggataaggcgcagcggtcgggctgaacggggggttcgtgcacacagcccagcttggagcgaacgacctacaccgaactgagatacctacagcgtgagctatgagaaagcgccacgcttcccgaagggagaaaggcggacaggtatccggtaagcggcagggtcggaacaggagagcgcacgagggagcttccagggggaaacgcctggtatctttatagtcctgtcgggtttcgccacctctgacttgagcgtcgatttttgtgatgctcgtcaggggggcggagcctatggaaaaacgccagcaacgcggcctttttacggttcctggccttttgctggccttttgctcacatgttctttcctgcgttatcccctgattctgtggataaccgtattaccgcctttgagtgagctgataccgctcgccgcagccgaacgaccgagcgcagcgagtcagtgagcgaggaagcggaagagcgcctgatgcggtattttctccttacgcatctgtgcggtatttcacaccgcatatatggtgcactctcagtacaatctgctctgatgccgcatagttaagccagtatacactccgctatcgctacgtgactgggtcatggctgcgccccgacacccgccaacacccgctgacgcgccctgacgggcttgtctgctcccggcatccgcttacagacaagctgtgaccgtctccgggagctgcatgtgtcagaggttttcaccgtcatcaccgaaacgcgcgaggcagctgcggtaaagctcatcagcgtggtcgtgaagcgattcacagatgtctgcctgttcatccgcgtccagctcgttgagtttctccagaagcgttaatgtctggcttctgataaagcgggccatgttaagggcggttttttcctgtttggtcactgatgcctccgtgtaagggggatttctgttcatgggggtaatgataccgatgaaacgagagaggatgctcacgatacgggttactgatgatgaacatgcccggttactggaacgttgtgagggtaaacaactggcggtatggatgcggcgggaccagagaaaaatcactcagggtcaatgccagcgcttcgttaatacagatgtaggtgttccacagggtagccagcagcatcctgcgatgcagatccggaa | ASKPDIKVGDYVKMGVYNNASILWRCVSIDNNGPLMLADKIVDTLAYDAKTNDNSNSKSHSRSYKRDDYGSNYWKDSNMRSWLNSTAAEGKVDWLCGNPPKDGYVSGVGAYNEKAGFLNAFSKSEIAAMKTVTQRSLVSHPEYNKGIVDGDANSDLLYYTDISEAVANYDSSYFETTTEKVFLLDVKQANAVWKNLKGYYVAYNNDGMAWPYWLRTPVTDCNHDMRYISSSGQVGRYAPWYSDLGVRPAFYLDSEYFVTTSGSGSQSSPYIGSAPNKQEDDYTISEPAEDANPDWNVSTEQSIQLTLGPWYSNDGKYSNPTIPVYTIQKTRSDTENMVVVVCGEGYTKSQQGKFINDVKRLWQDAMKYEPYRSYADRFNVYALCTASESTFDNGGSTFFDVIVDKYNSPVISNNLHGSQWKNHIFERCIGPEFIEKIHDAHIKKKCDPNTIPSGSEYEPYYYVHDYIAQFAMVVNTKSDFGGAYNNREYGFHYFISPSDSYRASKTFAHEFGHGLLGLGDEYSNGYLLDDKELKSLNLSSVEDPEKIKWRQLLGFRNTYTCRNAYGSKMLVSSYECIMRDTNYQFCEVCRLQGFKRMSQLVKDVDLYVATPEVKEYTGAYSKPSDFTDLETSSYYNYTYNRNDRLLSGNSKSRFNTNMNGKKIELRTVIQNISDKNARQLKFKMWIKHSDGSVATDSSGNPLQTVQTFDIPVWNDKANFWPLGALDHIKSDFNSGLKSCSLIYQIPSDAQLKSGDTVAFQVLDENGNVLADDNTETQ |
| MD | gatggtgtccgggatctcgacgctctcccttatgcgactcctgcattaggaagcagcccagtagtaggttgaggccgttgagcaccgccgccgcaaggaatggtgcatgcaaggagatggcgcccaacagtcccccggccacggggcctgccaccatacccacgccgaaacaagcgctcatgagcccgaagtggcgagcccgatcttccccatcggtgatgtcggcgatataggcgccagcaaccgcacctgtggcgccggtgatgccggccacgatgcgtccggcgtagaggatcgagatctcgatcccgcgaaattaatacgactcactataggggaattgtgagcggataacaattcccctctagaaataattttgtttaactttaagaaggagatataccatgggtcatcaccatcatcatcacgggtcggactcagaagtcaatcaagaagctaagccagaggtcaagccagaagtcaagcctgagactcacatcaatttaaaggtgtccgatggatcttcagagatcttcttcaagatcaaaaagaccactcctttaagaaggctgatggaagcgttcgctaaaagacagggtaaggaaatggactccttaagattcttgtacgacggtattagaattcaagctgatcaggcccctgaagatttggacatggaggataacgatattattgaggctcaccgcgaacagattggaggtaccgagcagagcatccaactgaccctcggcccttggtattccaacgacggtaaatacagcaacccgaccatcccggtttacaccatccaaaaaacgcgttccgacaccgagaatatggtggtggtcgtgtgcggtgagggctacactaagagccagcagggtaaattcatcaacgacgtaaagcgcctgtggcaggatgctatgaagtatgagccgtaccgctcctacgcggatcgtttcaatgtgtacgctctgtgtaccgcgtcagagagcaccttcgacaacggtggttctacgttctttgatgtgatagttgacaaatacaactccccggttatttccaacaacctgcatggttcccaatggaagaaccacatttttgagcgttgtattggtccggagtttatcgaaaagattcatgatgcgcatattaaaaaaaagtgcgatccgaataccattccgtcggggagcgagtatgaaccgtactattacgtgcatgactacattgcgcagtttgctatggttgtgaataccaaaagtgacttcggcggtgcgtacaataaccgtgaatacggcttccattatttcatcagcccgagcgacagctatcgtgcaagcaaaaccttcgcgcacgaattcggtcacggcctgcttggcttgggtgatgaatactccaacgggtacctgttggatgacaaggaactgaagagcctgaacctgagcagcgtcgaggacccggaaaaaatcaagtggcgtcaattgctgggttttcgcaacacatatacgtgccgtaatgcgtacggctctaaaatgctggtttctagctacgagtgcatcatgcgcgacacgaattatcagttctgcgaagtttgccgcctgcaaggtttcaaacgcatgagccagttggtgaaggatgtcgacttgtatgtcgccaccccagaggttaaagaatacaccggcgcatacagcaagccgtccgactttaccgatttggaaacctccagctactacaactacacctataaccgcaatgaccgcttgcttagcggtaattcgaagagccgtttcaataccaatatgaacggtaaaaaaatcgagttgcgtaccgttatacagaatattagcgacaagaatgcgcgtcaactgaaatttaagatgtggattaaacacagcgatggtagcgtggcaacggactcgtccggcaacccgctgcagaccgtccagaccttcgatattccggtttggaacgataaggctaacttttggcctctgggtgcgttggaccacatcaagagcgacttcaacagcggtctgaagagctgtagcctgatttatcagattccgtctgatgcccaactaaagtccggcgacaccgttgcgttccaggtcctggacgaaaatggcaatgtgttggctgacgataacaccgaaacccagtgactcgagcaccaccaccaccaccactgagatccggctgctaacaaagcccgaaaggaagctgagttggctgctgccaccgctgagcaataactagcataaccccttggggcctctaaacgggtcttgaggggttttttgctgaaaggaggaactatatccggattggcgaatgggacgcgccctgtagcggcgcattaagcgcggcgggtgtggtggttacgcgcagcgtgaccgctacacttgccagcgccctagcgcccgctcctttcgctttcttcccttcctttctcgccacgttcgccggctttccccgtcaagctctaaatcgggggctccctttagggttccgatttagtgctttacggcacctcgaccccaaaaaacttgattagggtgatggttcacgtagtgggccatcgccctgatagacggtttttcgccctttgacgttggagtccacgttctttaatagtggactcttgttccaaactggaacaacactcaaccctatctcggtctattcttttgatttataagggattttgccgatttcggcctattggttaaaaaatgagctgatttaacaaaaatttaacgcgaattttaacaaaatattaacgcttacaatttaggtggcacttttcggggaaatgtgcgcggaacccctatttgtttatttttctaaatacattcaaatatgtatccgctcatgaattaattcttagaaaaactcatcgagcatcaaatgaaactgcaatttattcatatcaggattatcaataccatatttttgaaaaagccgtttctgtaatgaaggagaaaactcaccgaggcagttccataggatggcaagatcctggtatcggtctgcgattccgactcgtccaacatcaatacaacctattaatttcccctcgtcaaaaataaggttatcaagtgagaaatcaccatgagtgacgactgaatccggtgagaatggcaaaagtttatgcatttctttccagacttgttcaacaggccagccattacgctcgtcatcaaaatcactcgcatcaaccaaaccgttattcattcgtgattgcgcctgagcgagacgaaatacgcgatcgctgttaaaaggacaattacaaacaggaatcgaatgcaaccggcgcaggaacactgccagcgcatcaacaatattttcacctgaatcaggatattcttctaatacctggaatgctgttttcccggggatcgcagtggtgagtaaccatgcatcatcaggagtacggataaaatgcttgatggtcggaagaggcataaattccgtcagccagtttagtctgaccatctcatctgtaacatcattggcaacgctacctttgccatgtttcagaaacaactctggcgcatcgggcttcccatacaatcgatagattgtcgcacctgattgcccgacattatcgcgagcccatttatacccatataaatcagcatccatgttggaatttaatcgcggcctagagcaagacgtttcccgttgaatatggctcataacaccccttgtattactgtttatgtaagcagacagttttattgttcatgaccaaaatcccttaacgtgagttttcgttccactgagcgtcagaccccgtagaaaagatcaaaggatcttcttgagatcctttttttctgcgcgtaatctgctgcttgcaaacaaaaaaaccaccgctaccagcggtggtttgtttgccggatcaagagctaccaactctttttccgaaggtaactggcttcagcagagcgcagataccaaatactgtccttctagtgtagccgtagttaggccaccacttcaagaactctgtagcaccgcctacatacctcgctctgctaatcctgttaccagtggctgctgccagtggcgataagtcgtgtcttaccgggttggactcaagacgatagttaccggataaggcgcagcggtcgggctgaacggggggttcgtgcacacagcccagcttggagcgaacgacctacaccgaactgagatacctacagcgtgagctatgagaaagcgccacgcttcccgaagggagaaaggcggacaggtatccggtaagcggcagggtcggaacaggagagcgcacgagggagcttccagggggaaacgcctggtatctttatagtcctgtcgggtttcgccacctctgacttgagcgtcgatttttgtgatgctcgtcaggggggcggagcctatggaaaaacgccagcaacgcggcctttttacggttcctggccttttgctggccttttgctcacatgttctttcctgcgttatcccctgattctgtggataaccgtattaccgcctttgagtgagctgataccgctcgccgcagccgaacgaccgagcgcagcgagtcagtgagcgaggaagcggaagagcgcctgatgcggtattttctccttacgcatctgtgcggtatttcacaccgcatatatggtgcactctcagtacaatctgctctgatgccgcatagttaagccagtatacactccgctatcgctacgtgactgggtcatggctgcgccccgacacccgccaacacccgctgacgcgccctgacgggcttgtctgctcccggcatccgcttacagacaagctgtgaccgtctccgggagctgcatgtgtcagaggttttcaccgtcatcaccgaaacgcgcgaggcagctgcggtaaagctcatcagcgtggtcgtgaagcgattcacagatgtctgcctgttcatccgcgtccagctcgttgagtttctccagaagcgttaatgtctggcttctgataaagcgggccatgttaagggcggttttttcctgtttggtcactgatgcctccgtgtaagggggatttctgttcatgggggtaatgataccgatgaaacgagagaggatgctcacgatacgggttactgatgatgaacatgcccggttactggaacgttgtgagggtaaacaactggcggtatggatgcggcgggaccagagaaaaatcactcagggtcaatgccagcgcttcgttaatacagatgtaggtgttccacagggtagccagcagcatcctgcgatgcagatccggaa | TEQSIQLTLGPWYSNDGKYSNPTIPVYTIQKTRSDTENMVVVVCGEGYTKSQQGKFINDVKRLWQDAMKYEPYRSYADRFNVYALCTASESTFDNGGSTFFDVIVDKYNSPVISNNLHGSQWKNHIFERCIGPEFIEKIHDAHIKKKCDPNTIPSGSEYEPYYYVHDYIAQFAMVVNTKSDFGGAYNNREYGFHYFISPSDSYRASKTFAHEFGHGLLGLGDEYSNGYLLDDKELKSLNLSSVEDPEKIKWRQLLGFRNTYTCRNAYGSKMLVSSYECIMRDTNYQFCEVCRLQGFKRMSQLVKDVDLYVATPEVKEYTGAYSKPSDFTDLETSSYYNYTYNRNDRLLSGNSKSRFNTNMNGKKIELRTVIQNISDKNARQLKFKMWIKHSDGSVATDSSGNPLQTVQTFDIPVWNDKANFWPLGALDHIKSDFNSGLKSCSLIYQIPSDAQLKSGDTVAFQVLDENGNVLADDNTETQ |
| MD+CTD1 | cgcattgcgcccagcgccatctgatcgttggcaaccagcatcgcagtgggaacgatgccctcattcagcatttgcatggtttgttgaaaaccggacatggcactccagtcgccttcccgttccgctatcggctgaatttgattgcgagtgagatatttatgccagccagccagacgcagacgcgccgagacagaacttaatgggcccgctaacagcgcgatttgctggtgacccaatgcgaccagatgctccacgcccagtcgcgtaccgtcttcatgggagaaaataatactgttgatgggtgtctggtcagagacatcaagaaataacgccggaacattagtgcaggcagcttccacagcaatggcatcctggtcatccagcggatagttaatgatcagcccactgacgcgttgcgcgagaagattgtgcaccgccgctttacaggcttcgacgccgcttcgttctaccatcgacaccaccacgctggcacccagttgatcggcgcgagatttaatcgccgcgacaatttgcgacggcgcgtgcagggccagactggaggtggcaacgccaatcagcaacgactgtttgcccgccagttgttgtgccacgcggttgggaatgtaattcagctccgccatcgccgcttccactttttcccgcgttttcgcagaaacgtggctggcctggttcaccacgcgggaaacggtctgataagagacaccggcatactctgcgacatcgtataacgttactggtttcacattcaccaccctgaattgactctcttccgggcgctatcatgccataccgcgaaaggttttgcgccattcgatggtgtccgggatctcgacgctctcccttatgcgactcctgcattaggaagcagcccagtagtaggttgaggccgttgagcaccgccgccgcaaggaatggtgcatgcaaggagatggcgcccaacagtcccccggccacggggcctgccaccatacccacgccgaaacaagcgctcatgagcccgaagtggcgagcccgatcttccccatcggtgatgtcggcgatataggcgccagcaaccgcacctgtggcgccggtgatgccggccacgatgcgtccggcgtagaggatcgagatctcgatcccgcgaaattaatacgactcactataggggaattgtgagcggataacaattcccctctagaaataattttgtttaactttaagaaggagatataccatgggtcatcaccatcatcatcacgggtcggactcagaagtcaatcaagaagctaagccagaggtcaagccagaagtcaagcctgagactcacatcaatttaaaggtgtccgatggatcttcagagatcttcttcaagatcaaaaagaccactcctttaagaaggctgatggaagcgttcgctaaaagacagggtaaggaaatggactccttaagattcttgtacgacggtattagaattcaagctgatcaggcccctgaagatttggacatggaggataacgatattattgaggctcaccgcgaacagattggaggtaccgagcagagcatccaactgaccctcggcccttggtattccaacgacggtaaatacagcaacccgaccatcccggtttacaccatccaaaaaacgcgttccgacaccgagaatatggtggtggtcgtgtgcggtgagggctacactaagagccagcagggtaaattcatcaacgacgtaaagcgcctgtggcaggatgctatgaagtatgagccgtaccgctcctacgcggatcgtttcaatgtgtacgctctgtgtaccgcgtcagagagcaccttcgacaacggtggttctacgttctttgatgtgatagttgacaaatacaactccccggttatttccaacaacctgcatggttcccaatggaagaaccacatttttgagcgttgtattggtccggagtttatcgaaaagattcatgatgcgcatattaaaaaaaagtgcgatccgaataccattccgtcggggagcgagtatgaaccgtactattacgtgcatgactacattgcgcagtttgctatggttgtgaataccaaaagtgacttcggcggtgcgtacaataaccgtgaatacggcttccattatttcatcagcccgagcgacagctatcgtgcaagcaaaaccttcgcgcacgaattcggtcacggcctgcttggcttgggtgatgaatactccaacgggtacctgttggatgacaaggaactgaagagcctgaacctgagcagcgtcgaggacccggaaaaaatcaagtggcgtcaattgctgggttttcgcaacacatatacgtgccgtaatgcgtacggctctaaaatgctggtttctagctacgagtgcatcatgcgcgacacgaattatcagttctgcgaagtttgccgcctgcaaggtttcaaacgcatgagccagttggtgaaggatgtcgacttgtatgtcgccaccccagaggttaaagaatacaccggcgcatacagcaagccgtccgactttaccgatttggaaacctccagctactacaactacacctataaccgcaatgaccgcttgcttagcggtaattcgaagagccgtttcaataccaatatgaacggtaaaaaaatcgagttgcgtaccgttatacagaatattagcgacaagaatgcgcgtcaactgaaatttaagatgtggattaaacacagcgatggtagcgtggcaacggactcgtccggcaacccgctgcagaccgtccagaccttcgatattccggtttggaacgataaggctaacttttggcctctgggtgcgttggaccacatcaagagcgacttcaacagcggtctgaagagctgtagcctgatttatcagattccgtctgatgcccaactaaagtccggcgacaccgttgcgttccaggtcctggacgaaaatggcaatgtgttggctgacgataacaccgaaacccagcgctatactacggttagcattcaatataaattcgaagacggttcggagattccgaataccgcgggtggcaccttcaccgttccgtacggtacgaagctggatttgaccccagccaaaacgctgtatgattatgaatttatcaaagttgatggcttaaataagccgattgtctccgacggcactgtggtgacgtactactacaagaactaactcgagcaccaccaccaccaccactgagatccggctgctaacaaagcccgaaaggaagctgagttggctgctgccaccgctgagcaataactagcataaccccttggggcctctaaacgggtcttgaggggttttttgctgaaaggaggaactatatccggattggcgaatgggacgcgccctgtagcggcgcattaagcgcggcgggtgtggtggttacgcgcagcgtgaccgctacacttgccagcgccctagcgcccgctcctttcgctttcttcccttcctttctcgccacgttcgccggctttccccgtcaagctctaaatcgggggctccctttagggttccgatttagtgctttacggcacctcgaccccaaaaaacttgattagggtgatggttcacgtagtgggccatcgccctgatagacggtttttcgccctttgacgttggagtccacgttctttaatagtggactcttgttccaaactggaacaacactcaaccctatctcggtctattcttttgatttataagggattttgccgatttcggcctattggttaaaaaatgagctgatttaacaaaaatttaacgcgaattttaacaaaatattaacgcttacaatttaggtggcacttttcggggaaatgtgcgcggaacccctatttgtttatttttctaaatacattcaaatatgtatccgctcatgaattaattcttagaaaaactcatcgagcatcaaatgaaactgcaatttattcatatcaggattatcaataccatatttttgaaaaagccgtttctgtaatgaaggagaaaactcaccgaggcagttccataggatggcaagatcctggtatcggtctgcgattccgactcgtccaacatcaatacaacctattaatttcccctcgtcaaaaataaggttatcaagtgagaaatcaccatgagtgacgactgaatccggtgagaatggcaaaagtttatgcatttctttccagacttgttcaacaggccagccattacgctcgtcatcaaaatcactcgcatcaaccaaaccgttattcattcgtgattgcgcctgagcgagacgaaatacgcgatcgctgttaaaaggacaattacaaacaggaatcgaatgcaaccggcgcaggaacactgccagcgcatcaacaatattttcacctgaatcaggatattcttctaatacctggaatgctgttttcccggggatcgcagtggtgagtaaccatgcatcatcaggagtacggataaaatgcttgatggtcggaagaggcataaattccgtcagccagtttagtctgaccatctcatctgtaacatcattggcaacgctacctttgccatgtttcagaaacaactctggcgcatcgggcttcccatacaatcgatagattgtcgcacctgattgcccgacattatcgcgagcccatttatacccatataaatcagcatccatgttggaatttaatcgcggcctagagcaagacgtttcccgttgaatatggctcataacaccccttgtattactgtttatgtaagcagacagttttattgttcatgaccaaaatcccttaacgtgagttttcgttccactgagcgtcagaccccgtagaaaagatcaaaggatcttcttgagatcctttttttctgcgcgtaatctgctgcttgcaaacaaaaaaaccaccgctaccagcggtggtttgtttgccggatcaagagctaccaactctttttccgaaggtaactggcttcagcagagcgcagataccaaatactgtccttctagtgtagccgtagttaggccaccacttcaagaactctgtagcaccgcctacatacctcgctctgctaatcctgttaccagtggctgctgccagtggcgataagtcgtgtcttaccgggttggactcaagacgatagttaccggataaggcgcagcggtcgggctgaacggggggttcgtgcacacagcccagcttggagcgaacgacctacaccgaactgagatacctacagcgtgagctatgagaaagcgccacgcttcccgaagggagaaaggcggacaggtatccggtaagcggcagggtcggaacaggagagcgcacgagggagcttccagggggaaacgcctggtatctttatagtcctgtcgggtttcgccacctctgacttgagcgtcgatttttgtgatgctcgtcaggggggcggagcctatggaaaaacgccagcaacgcggcctttttacggttcctggccttttgctggccttttgctcacatgttctttcctgcgttatcccctgattctgtggataaccgtattaccgcctttgagtgagctgataccgctcgccgcagccgaacgaccgagcgcagcgagtcagtgagcgaggaagcggaagagcgcctgatgcggtattttctccttacgcatctgtgcggtatttcacaccgcaatggtgcactctcagtacaatctgctctgatgccgcatagttaagccagtatacactccgctatcgctacgtgactgggtcatggctgcgccccgacacccgccaacacccgctgacgcgccctgacgggcttgtctgctcccggcatccgcttacagacaagctgtgaccgtctccgggagctgcatgtgtcagaggttttcaccgtcatcaccgaaacgcgcgaggcagctgcggtaaagctcatcagcgtggtcgtgaagcgattcacagatgtctgcctgttcatccgcgtccagctcgttgagtttctccagaagcgttaatgtctggcttctgataaagcgggccatgttaagggcggttttttcctgtttggtcactgatgcctccgtgtaagggggatttctgttcatgggggtaatgataccgatgaaacgagagaggatgctcacgatacgggttactgatgatgaacatgcccggttactggaacgttgtgagggtaaacaactggcggtatggatgcggcgggaccagagaaaaatcactcagggtcaatgccagcgcttcgttaatacagatgtaggtgttccacagggtagccagcagcatcctgcgatgcagatccggaacataatggtgcagggcgctgacttccgcgtttccagactttacgaaacacggaaaccgaagaccattcatgttgttgctcaggtcgcagacgttttgcagcagcagtcgcttcacgttcgctcgcgtatcggtgattcattctgctaaccagtaaggcaaccccgccagcctagccgggtcctcaacgacaggagcacgatcatgcgcacccgtggggccgccatgccggcgataatggcctgcttctcgccgaaacgtttggtggcgggaccagtgacgaaggcttgagcgagggcgtgcaagattccgaataccgcaagcgacaggccgatcatcgtcgcgctccagcgaaagcggtcctcgccgaaaatgacccagagcgctgccggcacctgtcctacgagttgcatgataaagaagacagtcataagtgcggcgacgatagtcatgccccgcgcccaccggaaggagctgactgggttgaaggctctcaagggcatcggtcgagatcccggtgcctaatgagtgagctaacttacattaattgcgttgcgctcactgcccgctttccagtcgggaaacctgtcgtgccagctgcattaatgaatcggccaacgcgcggggagaggcggtttgcgtattgggcgccagggtggtttttcttttcaccagtgagacgggcaacagctgattgcccttcaccgcctggccctgagagagttgcagcaagcggtccacgctggtttgccccagcaggcgaaaatcctgtttgatggtggttaacggcgggatataacatgagctgtcttcggtatcgtcgtatcccactaccgagatatccgcaccaacgcgcagcccggactcggtaatggcg | TEQSIQLTLGPWYSNDGKYSNPTIPVYTIQKTRSDTENMVVVVCGEGYTKSQQGKFINDVKRLWQDAMKYEPYRSYADRFNVYALCTASESTFDNGGSTFFDVIVDKYNSPVISNNLHGSQWKNHIFERCIGPEFIEKIHDAHIKKKCDPNTIPSGSEYEPYYYVHDYIAQFAMVVNTKSDFGGAYNNREYGFHYFISPSDSYRASKTFAHEFGHGLLGLGDEYSNGYLLDDKELKSLNLSSVEDPEKIKWRQLLGFRNTYTCRNAYGSKMLVSSYECIMRDTNYQFCEVCRLQGFKRMSQLVKDVDLYVATPEVKEYTGAYSKPSDFTDLETSSYYNYTYNRNDRLLSGNSKSRFNTNMNGKKIELRTVIQNISDKNARQLKFKMWIKHSDGSVATDSSGNPLQTVQTFDIPVWNDKANFWPLGALDHIKSDFNSGLKSCSLIYQIPSDAQLKSGDTVAFQVLDENGNVLADDNTETQRYTTVSIQYKFEDGSEIPNTAGGTFTVPYGTKLDLTPAKTLYDYEFIKVDGLNKPIVSDGTVVTYYYKN |
| CTD2 | cgcattgcgcccagcgccatctgatcgttggcaaccagcatcgcagtgggaacgatgccctcattcagcatttgcatggtttgttgaaaaccggacatggcactccagtcgccttcccgttccgctatcggctgaatttgattgcgagtgagatatttatgccagccagccagacgcagacgcgccgagacagaacttaatgggcccgctaacagcgcgatttgctggtgacccaatgcgaccagatgctccacgcccagtcgcgtaccgtcttcatgggagaaaataatactgttgatgggtgtctggtcagagacatcaagaaataacgccggaacattagtgcaggcagcttccacagcaatggcatcctggtcatccagcggatagttaatgatcagcccactgacgcgttgcgcgagaagattgtgcaccgccgctttacaggcttcgacgccgcttcgttctaccatcgacaccaccacgctggcacccagttgatcggcgcgagatttaatcgccgcgacaatttgcgacggcgcgtgcagggccagactggaggtggcaacgccaatcagcaacgactgtttgcccgccagttgttgtgccacgcggttgggaatgtaattcagctccgccatcgccgcttccactttttcccgcgttttcgcagaaacgtggctggcctggttcaccacgcgggaaacggtctgataagagacaccggcatactctgcgacatcgtataacgttactggtttcacattcaccaccctgaattgactctcttccgggcgctatcatgccataccgcgaaaggttttgcgccattcgatggtgtccgggatctcgacgctctcccttatgcgactcctgcattaggaagcagcccagtagtaggttgaggccgttgagcaccgccgccgcaaggaatggtgcatgcaaggagatggcgcccaacagtcccccggccacggggcctgccaccatacccacgccgaaacaagcgctcatgagcccgaagtggcgagcccgatcttccccatcggtgatgtcggcgatataggcgccagcaaccgcacctgtggcgccggtgatgccggccacgatgcgtccggcgtagaggatcgagatctcgatcccgcgaaattaatacgactcactataggggaattgtgagcggataacaattcccctctagaaataattttgtttaactttaagaaggagatataccatgggtcatcaccatcatcatcacgggtcggactcagaagtcaatcaagaagctaagccagaggtcaagccagaagtcaagcctgagactcacatcaatttaaaggtgtccgatggatcttcagagatcttcttcaagatcaaaaagaccactcctttaagaaggctgatggaagcgttcgctaaaagacagggtaaggaaatggactccttaagattcttgtacgacggtattagaattcaagctgatcaggcccctgaagatttggacatggaggataacgatattattgaggctcaccgcgaacagattggaggtgaacatacccataatctgacgctggtggcggcgaaggctgctacctgcacaaccgcgggcaactcagcctactacacgtgcgatggctgtgataaatggtttgcggacgcgacaggctccgtcgaaatcaccgacaaaaccagcgtgaagatcccggcgccgggtcataccgcgggaaccgagtggaaaagcgacgataccaatcattggcacgagtgcactgtggcgggttgcggtgtgattatcgagagcactaagagtgcgcacaccgcgggtgaatggatcgtggataccccagctaccgcgactactgctggcaccaagcacaaagagtgtactgtgtgtcatcgtgttcttgagacgcaaccgattccgagcaccggtacctaactcgagcaccaccaccaccaccactgagatccggctgctaacaaagcccgaaaggaagctgagttggctgctgccaccgctgagcaataactagcataaccccttggggcctctaaacgggtcttgaggggttttttgctgaaaggaggaactatatccggattggcgaatgggacgcgccctgtagcggcgcattaagcgcggcgggtgtggtggttacgcgcagcgtgaccgctacacttgccagcgccctagcgcccgctcctttcgctttcttcccttcctttctcgccacgttcgccggctttccccgtcaagctctaaatcgggggctccctttagggttccgatttagtgctttacggcacctcgaccccaaaaaacttgattagggtgatggttcacgtagtgggccatcgccctgatagacggtttttcgccctttgacgttggagtccacgttctttaatagtggactcttgttccaaactggaacaacactcaaccctatctcggtctattcttttgatttataagggattttgccgatttcggcctattggttaaaaaatgagctgatttaacaaaaatttaacgcgaattttaacaaaatattaacgcttacaatttaggtggcacttttcggggaaatgtgcgcggaacccctatttgtttatttttctaaatacattcaaatatgtatccgctcatgaattaattcttagaaaaactcatcgagcatcaaatgaaactgcaatttattcatatcaggattatcaataccatatttttgaaaaagccgtttctgtaatgaaggagaaaactcaccgaggcagttccataggatggcaagatcctggtatcggtctgcgattccgactcgtccaacatcaatacaacctattaatttcccctcgtcaaaaataaggttatcaagtgagaaatcaccatgagtgacgactgaatccggtgagaatggcaaaagtttatgcatttctttccagacttgttcaacaggccagccattacgctcgtcatcaaaatcactcgcatcaaccaaaccgttattcattcgtgattgcgcctgagcgagacgaaatacgcgatcgctgttaaaaggacaattacaaacaggaatcgaatgcaaccggcgcaggaacactgccagcgcatcaacaatattttcacctgaatcaggatattcttctaatacctggaatgctgttttcccggggatcgcagtggtgagtaaccatgcatcatcaggagtacggataaaatgcttgatggtcggaagaggcataaattccgtcagccagtttagtctgaccatctcatctgtaacatcattggcaacgctacctttgccatgtttcagaaacaactctggcgcatcgggcttcccatacaatcgatagattgtcgcacctgattgcccgacattatcgcgagcccatttatacccatataaatcagcatccatgttggaatttaatcgcggcctagagcaagacgtttcccgttgaatatggctcataacaccccttgtattactgtttatgtaagcagacagttttattgttcatgaccaaaatcccttaacgtgagttttcgttccactgagcgtcagaccccgtagaaaagatcaaaggatcttcttgagatcctttttttctgcgcgtaatctgctgcttgcaaacaaaaaaaccaccgctaccagcggtggtttgtttgccggatcaagagctaccaactctttttccgaaggtaactggcttcagcagagcgcagataccaaatactgtccttctagtgtagccgtagttaggccaccacttcaagaactctgtagcaccgcctacatacctcgctctgctaatcctgttaccagtggctgctgccagtggcgataagtcgtgtcttaccgggttggactcaagacgatagttaccggataaggcgcagcggtcgggctgaacggggggttcgtgcacacagcccagcttggagcgaacgacctacaccgaactgagatacctacagcgtgagctatgagaaagcgccacgcttcccgaagggagaaaggcggacaggtatccggtaagcggcagggtcggaacaggagagcgcacgagggagcttccagggggaaacgcctggtatctttatagtcctgtcgggtttcgccacctctgacttgagcgtcgatttttgtgatgctcgtcaggggggcggagcctatggaaaaacgccagcaacgcggcctttttacggttcctggccttttgctggccttttgctcacatgttctttcctgcgttatcccctgattctgtggataaccgtattaccgcctttgagtgagctgataccgctcgccgcagccgaacgaccgagcgcagcgagtcagtgagcgaggaagcggaagagcgcctgatgcggtattttctccttacgcatctgtgcggtatttcacaccgcaatggtgcactctcagtacaatctgctctgatgccgcatagttaagccagtatacactccgctatcgctacgtgactgggtcatggctgcgccccgacacccgccaacacccgctgacgcgccctgacgggcttgtctgctcccggcatccgcttacagacaagctgtgaccgtctccgggagctgcatgtgtcagaggttttcaccgtcatcaccgaaacgcgcgaggcagctgcggtaaagctcatcagcgtggtcgtgaagcgattcacagatgtctgcctgttcatccgcgtccagctcgttgagtttctccagaagcgttaatgtctggcttctgataaagcgggccatgttaagggcggttttttcctgtttggtcactgatgcctccgtgtaagggggatttctgttcatgggggtaatgataccgatgaaacgagagaggatgctcacgatacgggttactgatgatgaacatgcccggttactggaacgttgtgagggtaaacaactggcggtatggatgcggcgggaccagagaaaaatcactcagggtcaatgccagcgcttcgttaatacagatgtaggtgttccacagggtagccagcagcatcctgcgatgcagatccggaacataatggtgcagggcgctgacttccgcgtttccagactttacgaaacacggaaaccgaagaccattcatgttgttgctcaggtcgcagacgttttgcagcagcagtcgcttcacgttcgctcgcgtatcggtgattcattctgctaaccagtaaggcaaccccgccagcctagccgggtcctcaacgacaggagcacgatcatgcgcacccgtggggccgccatgccggcgataatggcctgcttctcgccgaaacgtttggtggcgggaccagtgacgaaggcttgagcgagggcgtgcaagattccgaataccgcaagcgacaggccgatcatcgtcgcgctccagcgaaagcggtcctcgccgaaaatgacccagagcgctgccggcacctgtcctacgagttgcatgataaagaagacagtcataagtgcggcgacgatagtcatgccccgcgcccaccggaaggagctgactgggttgaaggctctcaagggcatcggtcgagatcccggtgcctaatgagtgagctaacttacattaattgcgttgcgctcactgcccgctttccagtcgggaaacctgtcgtgccagctgcattaatgaatcggccaacgcgcggggagaggcggtttgcgtattgggcgccagggtggtttttcttttcaccagtgagacgggcaacagctgattgcccttcaccgcctggccctgagagagttgcagcaagcggtccacgctggtttgccccagcaggcgaaaatcctgtttgatggtggttaacggcgggatataacatgagctgtcttcggtatcgtcgtatcccactaccgagatatccgcaccaacgcgcagcccggactcggtaatggcg | EHTHNLTLVAAKAATCTTAGNSAYYTCDGCDKWFADATGSVEITDKTSVKIPAPGHTAGTEWKSDDTNHWHECTVAGCGVIIESTKSAHTAGEWIVDTPATATTAGTKHKECTVCHRVLETQPIPSTGT |
| CTD1-4 | cgcattgcgcccagcgccatctgatcgttggcaaccagcatcgcagtgggaacgatgccctcattcagcatttgcatggtttgttgaaaaccggacatggcactccagtcgccttcccgttccgctatcggctgaatttgattgcgagtgagatatttatgccagccagccagacgcagacgcgccgagacagaacttaatgggcccgctaacagcgcgatttgctggtgacccaatgcgaccagatgctccacgcccagtcgcgtaccgtcttcatgggagaaaataatactgttgatgggtgtctggtcagagacatcaagaaataacgccggaacattagtgcaggcagcttccacagcaatggcatcctggtcatccagcggatagttaatgatcagcccactgacgcgttgcgcgagaagattgtgcaccgccgctttacaggcttcgacgccgcttcgttctaccatcgacaccaccacgctggcacccagttgatcggcgcgagatttaatcgccgcgacaatttgcgacggcgcgtgcagggccagactggaggtggcaacgccaatcagcaacgactgtttgcccgccagttgttgtgccacgcggttgggaatgtaattcagctccgccatcgccgcttccactttttcccgcgttttcgcagaaacgtggctggcctggttcaccacgcgggaaacggtctgataagagacaccggcatactctgcgacatcgtataacgttactggtttcacattcaccaccctgaattgactctcttccgggcgctatcatgccataccgcgaaaggttttgcgccattcgatggtgtccgggatctcgacgctctcccttatgcgactcctgcattaggaagcagcccagtagtaggttgaggccgttgagcaccgccgccgcaaggaatggtgcatgcaaggagatggcgcccaacagtcccccggccacggggcctgccaccatacccacgccgaaacaagcgctcatgagcccgaagtggcgagcccgatcttccccatcggtgatgtcggcgatataggcgccagcaaccgcacctgtggcgccggtgatgccggccacgatgcgtccggcgtagaggatcgagatctcgatcccgcgaaattaatacgactcactataggggaattgtgagcggataacaattcccctctagaaataattttgtttaactttaagaaggagatataccatgggtcatcaccatcatcatcacgggtcggactcagaagtcaatcaagaagctaagccagaggtcaagccagaagtcaagcctgagactcacatcaatttaaaggtgtccgatggatcttcagagatcttcttcaagatcaaaaagaccactcctttaagaaggctgatggaagcgttcgctaaaagacagggtaaggaaatggactccttaagattcttgtacgacggtattagaattcaagctgatcaggcccctgaagatttggacatggaggataacgatattattgaggctcaccgcgaacagattggaggtacccagcgctatactacggttagcattcaatataaattcgaagacggttcggagattccgaataccgcgggtggcaccttcaccgttccgtacggtacgaagctggatttgaccccagccaaaacgctgtatgattatgaatttatcaaagttgatggcttaaataagccgattgtctccgacggcactgtggtgacgtactactacaagaacaagaacgaggaacatacccataatctgacgctggtggcggcgaaggctgctacctgcacaaccgcgggcaactcagcctactacacgtgcgatggctgtgataaatggtttgcggacgcgacaggctccgtcgaaatcaccgacaaaaccagcgtgaagatcccggcgccgggtcataccgcgggaaccgagtggaaaagcgacgataccaatcattggcacgagtgcactgtggcgggttgcggtgtgattatcgagagcactaagagtgcgcacaccgcgggtgaatggatcgtggataccccagctaccgcgactactgctggcaccaagcacaaagagtgtactgtgtgtcatcgtgttcttgagacgcaaccgattccgagcaccggtaccgagctgaagatcatagccggcgacaaccagatttataacaaagccagcggtagcgatgtaacgatcacctgtaatggtgattttgcaaaattcaccggtatcaaagttgacggcagtgtcgtggacagcagcaattacaccgcggtgtccggttctacggttctgacactgaaagcgagttatctgggcactctgactgacggttctcacaccatcaccttcgtttataccgacggcgaggcgaacgctaatttaacggtacgcaccgctggcagcgggcacatccacgattatggcactgagtggaaatccaatgcagataaccattggcacgaatgcaactgcggcgataagaaggatgaggcggctcactcgttcaagtgggtagtcgataaagaagccaccgctaccaaaaagggttctaagcacgaagagtgcaaaatctgcggttataagcgctctgcggtggaaatcccggcgaccggttaactcgagcaccaccaccaccaccactgagatccggctgctaacaaagcccgaaaggaagctgagttggctgctgccaccgctgagcaataactagcataaccccttggggcctctaaacgggtcttgaggggttttttgctgaaaggaggaactatatccggattggcgaatgggacgcgccctgtagcggcgcattaagcgcggcgggtgtggtggttacgcgcagcgtgaccgctacacttgccagcgccctagcgcccgctcctttcgctttcttcccttcctttctcgccacgttcgccggctttccccgtcaagctctaaatcgggggctccctttagggttccgatttagtgctttacggcacctcgaccccaaaaaacttgattagggtgatggttcacgtagtgggccatcgccctgatagacggtttttcgccctttgacgttggagtccacgttctttaatagtggactcttgttccaaactggaacaacactcaaccctatctcggtctattcttttgatttataagggattttgccgatttcggcctattggttaaaaaatgagctgatttaacaaaaatttaacgcgaattttaacaaaatattaacgcttacaatttaggtggcacttttcggggaaatgtgcgcggaacccctatttgtttatttttctaaatacattcaaatatgtatccgctcatgaattaattcttagaaaaactcatcgagcatcaaatgaaactgcaatttattcatatcaggattatcaataccatatttttgaaaaagccgtttctgtaatgaaggagaaaactcaccgaggcagttccataggatggcaagatcctggtatcggtctgcgattccgactcgtccaacatcaatacaacctattaatttcccctcgtcaaaaataaggttatcaagtgagaaatcaccatgagtgacgactgaatccggtgagaatggcaaaagtttatgcatttctttccagacttgttcaacaggccagccattacgctcgtcatcaaaatcactcgcatcaaccaaaccgttattcattcgtgattgcgcctgagcgagacgaaatacgcgatcgctgttaaaaggacaattacaaacaggaatcgaatgcaaccggcgcaggaacactgccagcgcatcaacaatattttcacctgaatcaggatattcttctaatacctggaatgctgttttcccggggatcgcagtggtgagtaaccatgcatcatcaggagtacggataaaatgcttgatggtcggaagaggcataaattccgtcagccagtttagtctgaccatctcatctgtaacatcattggcaacgctacctttgccatgtttcagaaacaactctggcgcatcgggcttcccatacaatcgatagattgtcgcacctgattgcccgacattatcgcgagcccatttatacccatataaatcagcatccatgttggaatttaatcgcggcctagagcaagacgtttcccgttgaatatggctcataacaccccttgtattactgtttatgtaagcagacagttttattgttcatgaccaaaatcccttaacgtgagttttcgttccactgagcgtcagaccccgtagaaaagatcaaaggatcttcttgagatcctttttttctgcgcgtaatctgctgcttgcaaacaaaaaaaccaccgctaccagcggtggtttgtttgccggatcaagagctaccaactctttttccgaaggtaactggcttcagcagagcgcagataccaaatactgtccttctagtgtagccgtagttaggccaccacttcaagaactctgtagcaccgcctacatacctcgctctgctaatcctgttaccagtggctgctgccagtggcgataagtcgtgtcttaccgggttggactcaagacgatagttaccggataaggcgcagcggtcgggctgaacggggggttcgtgcacacagcccagcttggagcgaacgacctacaccgaactgagatacctacagcgtgagctatgagaaagcgccacgcttcccgaagggagaaaggcggacaggtatccggtaagcggcagggtcggaacaggagagcgcacgagggagcttccagggggaaacgcctggtatctttatagtcctgtcgggtttcgccacctctgacttgagcgtcgatttttgtgatgctcgtcaggggggcggagcctatggaaaaacgccagcaacgcggcctttttacggttcctggccttttgctggccttttgctcacatgttctttcctgcgttatcccctgattctgtggataaccgtattaccgcctttgagtgagctgataccgctcgccgcagccgaacgaccgagcgcagcgagtcagtgagcgaggaagcggaagagcgcctgatgcggtattttctccttacgcatctgtgcggtatttcacaccgcaatggtgcactctcagtacaatctgctctgatgccgcatagttaagccagtatacactccgctatcgctacgtgactgggtcatggctgcgccccgacacccgccaacacccgctgacgcgccctgacgggcttgtctgctcccggcatccgcttacagacaagctgtgaccgtctccgggagctgcatgtgtcagaggttttcaccgtcatcaccgaaacgcgcgaggcagctgcggtaaagctcatcagcgtggtcgtgaagcgattcacagatgtctgcctgttcatccgcgtccagctcgttgagtttctccagaagcgttaatgtctggcttctgataaagcgggccatgttaagggcggttttttcctgtttggtcactgatgcctccgtgtaagggggatttctgttcatgggggtaatgataccgatgaaacgagagaggatgctcacgatacgggttactgatgatgaacatgcccggttactggaacgttgtgagggtaaacaactggcggtatggatgcggcgggaccagagaaaaatcactcagggtcaatgccagcgcttcgttaatacagatgtaggtgttccacagggtagccagcagcatcctgcgatgcagatccggaacataatggtgcagggcgctgacttccgcgtttccagactttacgaaacacggaaaccgaagaccattcatgttgttgctcaggtcgcagacgttttgcagcagcagtcgcttcacgttcgctcgcgtatcggtgattcattctgctaaccagtaaggcaaccccgccagcctagccgggtcctcaacgacaggagcacgatcatgcgcacccgtggggccgccatgccggcgataatggcctgcttctcgccgaaacgtttggtggcgggaccagtgacgaaggcttgagcgagggcgtgcaagattccgaataccgcaagcgacaggccgatcatcgtcgcgctccagcgaaagcggtcctcgccgaaaatgacccagagcgctgccggcacctgtcctacgagttgcatgataaagaagacagtcataagtgcggcgacgatagtcatgccccgcgcccaccggaaggagctgactgggttgaaggctctcaagggcatcggtcgagatcccggtgcctaatgagtgagctaacttacattaattgcgttgcgctcactgcccgctttccagtcgggaaacctgtcgtgccagctgcattaatgaatcggccaacgcgcggggagaggcggtttgcgtattgggcgccagggtggtttttcttttcaccagtgagacgggcaacagctgattgcccttcaccgcctggccctgagagagttgcagcaagcggtccacgctggtttgccccagcaggcgaaaatcctgtttgatggtggttaacggcgggatataacatgagctgtcttcggtatcgtcgtatcccactaccgagatatccgcaccaacgcgcagcccggactcggtaatggcg | TQRYTTVSIQYKFEDGSEIPNTAGGTFTVPYGTKLDLTPAKTLYDYEFIKVDGLNKPIVSDGTVVTYYYKNKNEEHTHNLTLVAAKAATCTTAGNSAYYTCDGCDKWFADATGSVEITDKTSVKIPAPGHTAGTEWKSDDTNHWHECTVAGCGVIIESTKSAHTAGEWIVDTPATATTAGTKHKECTVCHRVLETQPIPSTGTELKIIAGDNQIYNKASGSDVTITCNGDFAKFTGIKVDGSVVDSSNYTAVSGSTVLTLKASYLGTLTDGSHTITFVYTDGEANANLTVRTAGSGHIHDYGTEWKSNADNHWHECNCGDKKDEAAHSFKWVVDKEATATKKGSKHEECKICGYKRSAVEIPATG |
| MD+  CTD1-4 | atgctgaatgagggcatcgttcccactgcgatgctggttgccaacgatcagatggcgctgggcgcaatgcgcgccattaccgagtccgggctgcgcgttggtgcggatatctcggtagtgggatacgacgataccgaagatagctcatgttatatcccgccgttaaccaccatcaaacaggattttcgcctgctggggcaaaccagcgtggaccgcttgctgcaactctctcagggccaggcggtgaagggcaatcagctgttgccagtctcactggtgaaaagaaaaaccaccctggcgcccaatacgcaaaccgcctctccccgcgcgttggccgattcattaatgcagctggcacgacaggtttcccgactggaaagcgggcagtgactcatgaccaaaatcccttaacgtgagttacgcgcgcgtcgttccactgagcgtcagaccccgtagaaaagatcaaaggatcttcttgagatcctttttttctgcgcgtaatctgctgcttgcaaacaaaaaaaccaccgctaccagcggtggtttgtttgccggatcaagagctaccaactctttttccgaaggtaactggcttcagcagagcgcagataccaaatactgttcttctagtgtagccgtagttagcccaccacttcaagaactctgtagcaccgcctacatacctcgctctgctaatcctgttaccagtggctgctgccagtggcgataagtcgtgtcttaccgggttggactcaagacgatagttaccggataaggcgcagcggtcgggctgaacggggggttcgtgcacacagcccagcttggagcgaacgacctacaccgaactgagatacctacagcgtgagctatgagaaagcgccacgcttcccgaagggagaaaggcggacaggtatccggtaagcggcagggtcggaacaggagagcgcacgagggagcttccagggggaaacgcctggtatctttatagtcctgtcgggtttcgccacctctgacttgagcgtcgatttttgtgatgctcgtcaggggggcggagcctatggaaaaacgccagcaacgcggcctttttacggttcctggccttttgctggccttttgctcacatgttctttcctgcgttatcccctgattctgtggataaccgtattaccgcctttgagtgagctgataccgctcgccgcagccgaacgaccgagcgcagcgagtcagtgagcgaggaagcggaaggcgagagtagggaactgccaggcatcaaactaagcagaaggcccctgacggatggcctttttgcgtttctacaaactctttctgtgttgtaaaacgacggccagtcttaagctcgggccccctgggcggttctgataacgagtaatcgttaatccgcaaataacgtaaaaacccgcttcggcgggtttttttatggggggagtttagggaaagagcatttgtcagaatatttaagggcgcctgtcactttgcttgatatatgagaattatttaaccttataaatgagaaaaaagcaacgcactttaaataagatacgttgctttttcgattgatgaacacctataattaaactattcatctattatttatgattttttgtatatacaatatttctagtttgttaaagagaattaagaaaataaatctcgaaaataataaagggaaaatcagtttttgatatcaaaattatacatgtcaacgataatacaaaatataatacaaactataagatgttatcagtatttattatgcatttagaataaattttgtgtcgcccttaattgtgagcggataacaattacgagcttcatgcacagtgaaatcatgaaaaatttatttgctttgtgagcggataacaattataatatgtggaattgtgagcgctcacaattccacaacggtttccctctagaaataattttgtttaacttttaggaggtaaaacatatggcgaagtattacggcagtatgaaaatcgaagaaggtaaactggtaatctggattaacggcgataaaggctataacggtctcgctgaagtcggtaagaaattcgagaaagataccggaattaaagtcaccgttgagcatccggataaactggaagagaaattcccacaggttgcggcaactggcgatggccctgacattatcttctgggcacacgaccgctttggtggctacgctcaatctggcctgttggctgaaatcaccccggacaaagcgttccaggacaagctgtatccgtttacctgggatgccgtacgttacaacggcaagctgattgcttacccgatcgctgttgaagcgttatcgctgatttataacaaagatctgctgccgaacccgccaaaaacctgggaagagatcccggcgctggataaagaactgaaagcgaaaggtaagagcgcgctgatgttcaacctgcaagaaccgtacttcacctggccgctgattgctgctgacgggggttatgcgttcaagtatgaaaacggcaagtacgacattaaagacgtgggcgtggataacgctggcgcgaaagcgggtctgaccttcctggttgacctgattaaaaacaaacacatgaatgcagacaccgattactccatcgcagaagctgcctttaataaaggcgaaacagcgatgaccatcaacggcccgtgggcatggtccaacatcgacaccagcaaagtgaattatggtgtaacggtactgccgaccttcaagggtcaaccatccaaaccgttcgttggcgtgctgagcgcaggtattaacgccgccagtccgaacaaagagctggcaaaagagttcctcgaaaactatctgctgactgatgaaggtctggaagcggttaataaagacaaaccgctgggtgccgtagcgctgaagtcttacgaggaagagttggcgaaagatccacgtattgccgccactatggaaaacgcccagaaaggtgaaatcatgccgaacatcccgcagatgtccgctttctggtatgccgtgcgtactgcggtgatcaacgccgccagcggtcgtcagactgtcgatgaagccctgaaagacgcgcagactcgtatcactaaaagcggaagcggagagaacctgtacttccaatccctggtgccgcgcggcagcggatccaccgagcagagcatccaactgaccctcggcccttggtattccaacgacggtaaatacagcaacccgaccatcccggtttacaccatccaaaaaacgcgttccgacaccgagaatatggtggtggtcgtgtgcggtgagggctacactaagagccagcagggtaaattcatcaacgacgtaaagcgcctgtggcaggatgctatgaagtatgagccgtaccgctcctacgcggatcgtttcaatgtgtacgctctgtgtaccgcgtcagagagcaccttcgacaacggtggttctacgttctttgatgtgatagttgacaaatacaactccccggttatttccaacaacctgcatggttcccaatggaagaaccacatttttgagcgttgtattggtccggagtttatcgaaaagattcatgatgcgcatattaaaaaaaagtgcgatccgaataccattccgtcggggagcgagtatgaaccgtactattacgtgcatgactacattgcgcagtttgctatggttgtgaataccaaaagtgacttcggcggtgcgtacaataaccgtgaatacggcttccattatttcatcagcccgagcgacagctatcgtgcaagcaaaaccttcgcgcacgaattcggtcacggcctgcttggcttgggtgatgaatactccaacgggtacctgttggatgacaaggaactgaagagcctgaacctgagcagcgtcgaggacccggaaaaaatcaagtggcgtcaattgctgggttttcgcaacacatatacgtgccgtaatgcgtacggctctaaaatgctggtttctagctacgagtgcatcatgcgcgacacgaattatcagttctgcgaagtttgccgcctgcaaggtttcaaacgcatgagccagttggtgaaggatgtcgacttgtatgtcgccaccccagaggttaaagaatacaccggcgcatacagcaagccgtccgactttaccgatttggaaacctccagctactacaactacacctataaccgcaatgaccgcttgcttagcggtaattcgaagagccgtttcaataccaatatgaacggtaaaaaaatcgagttgcgtaccgttatacagaatattagcgacaagaatgcgcgtcaactgaaatttaagatgtggattaaacacagcgatggtagcgtggcaacggactcgtccggcaacccgctgcagaccgtccagaccttcgatattccggtttggaacgataaggctaacttttggcctctgggtgcgttggaccacatcaagagcgacttcaacagcggtctgaagagctgtagcctgatttatcagattccgtctgatgcccaactaaagtccggcgacaccgttgcgttccaggtcctggacgaaaatggcaatgtgttggctgacgataacaccgaaacccagcgctatactacggttagcattcaatataaattcgaagacggttcggagattccgaataccgcgggtggcaccttcaccgttccgtacggtacgaagctggatttgaccccagccaaaacgctgtatgattatgaatttatcaaagttgatggcttaaataagccgattgtctccgacggcactgtggtgacgtactactacaagaacaagaacgaggaacatacccataatctgacgctggtggcggcgaaggctgctacctgcacaaccgcgggcaactcagcctactacacgtgcgatggctgtgataaatggtttgcggacgcgacaggctccgtcgaaatcaccgacaaaaccagcgtgaagatcccggcgccgggtcataccgcgggaaccgagtggaaaagcgacgataccaatcattggcacgagtgcactgtggcgggttgcggtgtgattatcgagagcactaagagtgcgcacaccgcgggtgaatggatcgtggataccccagctaccgcgactactgctggcaccaagcacaaagagtgtactgtgtgtcatcgtgttcttgagacgcaaccgattccgagcaccggtaccgagctgaagatcatagccggcgacaaccagatttataacaaagccagcggtagcgatgtaacgatcacctgtaatggtgattttgcaaaattcaccggtatcaaagttgacggcagtgtcgtggacagcagcaattacaccgcggtgtccggttctacggttctgacactgaaagcgagttatctgggcactctgactgacggttctcacaccatcaccttcgtttataccgacggcgaggcgaacgctaatttaacggtacgcaccgctggcagcgggcacatccacgattatggcactgagtggaaatccaatgcagataaccattggcacgaatgcaactgcggcgataagaaggatgaggcggctcactcgttcaagtgggtagtcgataaagaagccaccgctaccaaaaagggttctaagcacgaagagtgcaaaatctgcggttataagcgctctgcggtggaaatcccggcgaccggatcccaccatcaccatcaccactaagactacaaagacgatgacgacaagtaactcgagccccaagggcgacaccccctaattagcccgggcgaaaggcccagtctttcgactgagcctttcgttttatttgatgcctggcagttccctactctcgcatggggagtccccacactaccatcggcgctacggcgtttcacttctgagttcggcatggggtcaggtgggaccaccgcgctactgccgccaggcaaacaaggggtgttatgagccatattcaggtataaatgggctcgcgataatgttcagaattggttaattggttgtaacactgacccctatttgtttatttttctaaatacattcaaatatgtatccgctcatgagacaataaccctgataaatgcttcaataatattgaaaaaggaagaatatgagccatattcaacgggaaacgtcgaggccgcgattaaattccaacatggatgctgatttatatgggtataaatgggctcgcgataatgtcgggcaatcaggtgcgacaatctatcgcttgtatgggaagcccgatgcgccagagttgtttctgaaacatggcaaaggtagcgttgccaatgatgttacagatgagatggtcagactaaactggctgacggaatttatgccacttccgaccatcaagcattttatccgtactcctgatgatgcatggttactcaccactgcgatccccggaaaaacagcgttccaggtattagaagaatatcctgattcaggtgaaaatattgttgatgcgctggcagtgttcctgcgccggttgcactcgattcctgtttgtaattgtccttttaacagcgatcgcgtatttcgcctcgctcaggcgcaatcacgaatgaataacggtttggttgatgcgagtgattttgatgacgagcgtaatggctggcctgttgaacaagtctggaaagaaatgcataaacttttgccattctcaccggattcagtcgtcactcatggtgatttctcacttgataaccttatttttgacgaggggaaattaataggttgtattgatgttggacgagtcggaatcgcagaccgataccaggatcttgccatcctatggaactgcctcggtgagttttctccttcattacagaaacggctttttcaaaaatatggtattgataatcctgatatgaataaattgcaatttcatttgatgctcgatgagtttttctaagcggcgcgccatcgaatggcgcaaaacctttcgcggtatggcatgatagcgcccggaagagagtcaattcagggtggtgaatatgaaaccagtaacgttatacgatgtcgcagagtatgccggtgtctcttatcagaccgtttcccgcgtggtgaaccaggccagccacgtttctgcgaaaacgcgggaaaaagtggaagcggcgatggcggagctgaattacattcccaaccgcgtggcacaacaactggcgggcaaacagtcgttgctgattggcgttgccacctccagtctggccctgcacgcgccgtcgcaaattgtcgcggcgattaaatctcgcgccgatcaactgggtgccagcgtggtggtgtcgatggtagaacgaagcggcgtcgaagcctgtaaagcggcggtgcacaatcttctcgcgcaacgcgtcagtgggctgatcattaactatccgctggatgaccaggatgccattgctgtggaagctgcctgcactaatgttccggcgttatttcttgatgtctctgaccagacacccatcaacagtattattttctcccatgaggacggtacgcgactgggcgtggagcatctggtcgcattgggtcaccagcaaatcgcgctgttagcgggcccattaagttctgtctcggcgcgtctgcgtctggctggctggcataaatatctcactcgcaatcaaattcagccgatagcggaacgggaaggcgactggagtgccatgtccggttttcaacaaaccatgcaa | GSGSTEQSIQLTLGPWYSNDGKYSNPTIPVYTIQKTRSDTENMVVVVCGEGYTKSQQGKFINDVKRLWQDAMKYEPYRSYADRFNVYALCTASESTFDNGGSTFFDVIVDKYNSPVISNNLHGSQWKNHIFERCIGPEFIEKIHDAHIKKKCDPNTIPSGSEYEPYYYVHDYIAQFAMVVNTKSDFGGAYNNREYGFHYFISPSDSYRASKTFAHEFGHGLLGLGDEYSNGYLLDDKELKSLNLSSVEDPEKIKWRQLLGFRNTYTCRNAYGSKMLVSSYECIMRDTNYQFCEVCRLQGFKRMSQLVKDVDLYVATPEVKEYTGAYSKPSDFTDLETSSYYNYTYNRNDRLLSGNSKSRFNTNMNGKKIELRTVIQNISDKNARQLKFKMWIKHSDGSVATDSSGNPLQTVQTFDIPVWNDKANFWPLGALDHIKSDFNSGLKSCSLIYQIPSDAQLKSGDTVAFQVLDENGNVLADDNTETQRYTTVSIQYKFEDGSEIPNTAGGTFTVPYGTKLDLTPAKTLYDYEFIKVDGLNKPIVSDGTVVTYYYKNKNEEHTHNLTLVAAKAATCTTAGNSAYYTCDGCDKWFADATGSVEITDKTSVKIPAPGHTAGTEWKSDDTNHWHECTVAGCGVIIESTKSAHTAGEWIVDTPATATTAGTKHKECTVCHRVLETQPIPSTGTELKIIAGDNQIYNKASGSDVTITCNGDFAKFTGIKVDGSVVDSSNYTAVSGSTVLTLKASYLGTLTDGSHTITFVYTDGEANANLTVRTAGSGHIHDYGTEWKSNADNHWHECNCGDKKDEAAHSFKWVVDKEATATKKGSKHEECKICGYKRSAVEIPATGSHHHHHH |
| FL | atgctgaatgagggcatcgttcccactgcgatgctggttgccaacgatcagatggcgctgggcgcaatgcgcgccattaccgagtccgggctgcgcgttggtgcggatatctcggtagtgggatacgacgataccgaagatagctcatgttatatcccgccgttaaccaccatcaaacaggattttcgcctgctggggcaaaccagcgtggaccgcttgctgcaactctctcagggccaggcggtgaagggcaatcagctgttgccagtctcactggtgaaaagaaaaaccaccctggcgcccaatacgcaaaccgcctctccccgcgcgttggccgattcattaatgcagctggcacgacaggtttcccgactggaaagcgggcagtgactcatgaccaaaatcccttaacgtgagttacgcgcgcgtcgttccactgagcgtcagaccccgtagaaaagatcaaaggatcttcttgagatcctttttttctgcgcgtaatctgctgcttgcaaacaaaaaaaccaccgctaccagcggtggtttgtttgccggatcaagagctaccaactctttttccgaaggtaactggcttcagcagagcgcagataccaaatactgttcttctagtgtagccgtagttagcccaccacttcaagaactctgtagcaccgcctacatacctcgctctgctaatcctgttaccagtggctgctgccagtggcgataagtcgtgtcttaccgggttggactcaagacgatagttaccggataaggcgcagcggtcgggctgaacggggggttcgtgcacacagcccagcttggagcgaacgacctacaccgaactgagatacctacagcgtgagctatgagaaagcgccacgcttcccgaagggagaaaggcggacaggtatccggtaagcggcagggtcggaacaggagagcgcacgagggagcttccagggggaaacgcctggtatctttatagtcctgtcgggtttcgccacctctgacttgagcgtcgatttttgtgatgctcgtcaggggggcggagcctatggaaaaacgccagcaacgcggcctttttacggttcctggccttttgctggccttttgctcacatgttctttcctgcgttatcccctgattctgtggataaccgtattaccgcctttgagtgagctgataccgctcgccgcagccgaacgaccgagcgcagcgagtcagtgagcgaggaagcggaaggcgagagtagggaactgccaggcatcaaactaagcagaaggcccctgacggatggcctttttgcgtttctacaaactctttctgtgttgtaaaacgacggccagtcttaagctcgggccccctgggcggttctgataacgagtaatcgttaatccgcaaataacgtaaaaacccgcttcggcgggtttttttatggggggagtttagggaaagagcatttgtcagaatatttaagggcgcctgtcactttgcttgatatatgagaattatttaaccttataaatgagaaaaaagcaacgcactttaaataagatacgttgctttttcgattgatgaacacctataattaaactattcatctattatttatgattttttgtatatacaatatttctagtttgttaaagagaattaagaaaataaatctcgaaaataataaagggaaaatcagtttttgatatcaaaattatacatgtcaacgataatacaaaatataatacaaactataagatgttatcagtatttattatgcatttagaataaattttgtgtcgcccttaattgtgagcggataacaattacgagcttcatgcacagtgaaatcatgaaaaatttatttgctttgtgagcggataacaattataatatgtggaattgtgagcgctcacaattccacaacggtttccctctagaaataattttgtttaacttttaggaggtaaaacatatggcgaagtattacggcagtatgaaaatcgaagaaggtaaactggtaatctggattaacggcgataaaggctataacggtctcgctgaagtcggtaagaaattcgagaaagataccggaattaaagtcaccgttgagcatccggataaactggaagagaaattcccacaggttgcggcaactggcgatggccctgacattatcttctgggcacacgaccgctttggtggctacgctcaatctggcctgttggctgaaatcaccccggacaaagcgttccaggacaagctgtatccgtttacctgggatgccgtacgttacaacggcaagctgattgcttacccgatcgctgttgaagcgttatcgctgatttataacaaagatctgctgccgaacccgccaaaaacctgggaagagatcccggcgctggataaagaactgaaagcgaaaggtaagagcgcgctgatgttcaacctgcaagaaccgtacttcacctggccgctgattgctgctgacgggggttatgcgttcaagtatgaaaacggcaagtacgacattaaagacgtgggcgtggataacgctggcgcgaaagcgggtctgaccttcctggttgacctgattaaaaacaaacacatgaatgcagacaccgattactccatcgcagaagctgcctttaataaaggcgaaacagcgatgaccatcaacggcccgtgggcatggtccaacatcgacaccagcaaagtgaattatggtgtaacggtactgccgaccttcaagggtcaaccatccaaaccgttcgttggcgtgctgagcgcaggtattaacgccgccagtccgaacaaagagctggcaaaagagttcctcgaaaactatctgctgactgatgaaggtctggaagcggttaataaagacaaaccgctgggtgccgtagcgctgaagtcttacgaggaagagttggcgaaagatccacgtattgccgccactatggaaaacgcccagaaaggtgaaatcatgccgaacatcccgcagatgtccgctttctggtatgccgtgcgtactgcggtgatcaacgccgccagcggtcgtcagactgtcgatgaagccctgaaagacgcgcagactcgtatcactaaaagcggaagcggagagaacctgtacttccaatccctggtgccgcgcggcagcggatccgcttcaaaacccgacataaaggtaggagattacgttaaaatgggtgtttataataacgcgagcattctgtggcgttgcgtttctatcgataacaacggtccgttaatgctggcagacaagattgtggataccctcgcatacgacgcgaaaaccaacgacaactctaacagcaagtcccacagccgtagctacaaacgtgatgattatggctccaactactggaaggactcaaacatgcgtagctggctcaatagcacggctgcggaaggtaaagtagactggctgtgcggtaacccgcccaaagatggctatgttagcggtgtgggtgcgtacaacgaaaaagccggctttctgaacgcatttagcaagtctgaaatcgcagcaatgaaaaccgttacccaacgttctctggtttcgcatccggagtacaacaaaggtatcgttgacggcgacgccaattctgatctgttgtactacacagacatcagtgaggcggttgcgaactatgactctagctacttcgaaactactacggagaaggtgtttctgttggacgttaaacaagcaaatgccgtgtggaaaaacctgaagggctattacgtggcgtataacaacgacggtatggcgtggccgtattggctgcgtaccccggtgaccgactgtaaccacgatatgcgttatatcagttccagcggccaggtcggtcgttatgctccgtggtatagcgacctgggcgtgagaccagcattttatctggactctgagtattttgttactacgagcggctctggctcccaaagcagtccgtatatcggctccgcgccgaacaaacaagaagatgactacaccatttccgagccagcagaagacgccaacccggattggaacgttagtaccgagcagagcatccaactgaccctcggcccttggtattccaacgacggtaaatacagcaacccgaccatcccggtttacaccatccaaaaaacgcgttccgacaccgagaatatggtggtggtcgtgtgcggtgagggctacactaagagccagcagggtaaattcatcaacgacgtaaagcgcctgtggcaggatgctatgaagtatgagccgtaccgctcctacgcggatcgtttcaatgtgtacgctctgtgtaccgcgtcagagagcaccttcgacaacggtggttctacgttctttgatgtgatagttgacaaatacaactccccggttatttccaacaacctgcatggttcccaatggaagaaccacatttttgagcgttgtattggtccggagtttatcgaaaagattcatgatgcgcatattaaaaaaaagtgcgatccgaataccattccgtcggggagcgagtatgaaccgtactattacgtgcatgactacattgcgcagtttgctatggttgtgaataccaaaagtgacttcggcggtgcgtacaataaccgtgaatacggcttccattatttcatcagcccgagcgacagctatcgtgcaagcaaaaccttcgcgcacgaattcggtcacggcctgcttggcttgggtgatgaatactccaacgggtacctgttggatgacaaggaactgaagagcctgaacctgagcagcgtcgaggacccggaaaaaatcaagtggcgtcaattgctgggttttcgcaacacatatacgtgccgtaatgcgtacggctctaaaatgctggtttctagctacgagtgcatcatgcgcgacacgaattatcagttctgcgaagtttgccgcctgcaaggtttcaaacgcatgagccagttggtgaaggatgtcgacttgtatgtcgccaccccagaggttaaagaatacaccggcgcatacagcaagccgtccgactttaccgatttggaaacctccagctactacaactacacctataaccgcaatgaccgcttgcttagcggtaattcgaagagccgtttcaataccaatatgaacggtaaaaaaatcgagttgcgtaccgttatacagaatattagcgacaagaatgcgcgtcaactgaaatttaagatgtggattaaacacagcgatggtagcgtggcaacggactcgtccggcaacccgctgcagaccgtccagaccttcgatattccggtttggaacgataaggctaacttttggcctctgggtgcgttggaccacatcaagagcgacttcaacagcggtctgaagagctgtagcctgatttatcagattccgtctgatgcccaactaaagtccggcgacaccgttgcgttccaggtcctggacgaaaatggcaatgtgttggctgacgataacaccgaaacccagcgctatactacggttagcattcaatataaattcgaagacggttcggagattccgaataccgcgggtggcaccttcaccgttccgtacggtacgaagctggatttgaccccagccaaaacgctgtatgattatgaatttatcaaagttgatggcttaaataagccgattgtctccgacggcactgtggtgacgtactactacaagaacaagaacgaggaacatacccataatctgacgctggtggcggcgaaggctgctacctgcacaaccgcgggcaactcagcctactacacgtgcgatggctgtgataaatggtttgcggacgcgacaggctccgtcgaaatcaccgacaaaaccagcgtgaagatcccggcgccgggtcataccgcgggaaccgagtggaaaagcgacgataccaatcattggcacgagtgcactgtggcgggttgcggtgtgattatcgagagcactaagagtgcgcacaccgcgggtgaatggatcgtggataccccagctaccgcgactactgctggcaccaagcacaaagagtgtactgtgtgtcatcgtgttcttgagacgcaaccgattccgagcaccggtaccgagctgaagatcatagccggcgacaaccagatttataacaaagccagcggtagcgatgtaacgatcacctgtaatggtgattttgcaaaattcaccggtatcaaagttgacggcagtgtcgtggacagcagcaattacaccgcggtgtccggttctacggttctgacactgaaagcgagttatctgggcactctgactgacggttctcacaccatcaccttcgtttataccgacggcgaggcgaacgctaatttaacggtacgcaccgctggcagcgggcacatccacgattatggcactgagtggaaatccaatgcagataaccattggcacgaatgcaactgcggcgataagaaggatgaggcggctcactcgttcaagtgggtagtcgataaagaagccaccgctaccaaaaagggttctaagcacgaagagtgcaaaatctgcggttataagcgctctgcggtggaaatcccggcgaccggtacttcaaccgcgccaaccgataccacgaaaccgaatgacaccaccaagccgggtaacacgaacggctctgaaaagggatcccaccatcaccatcaccactaagactacaaagacgatgacgacaagtaactcgagccccaagggcgacaccccctaattagcccgggcgaaaggcccagtctttcgactgagcctttcgttttatttgatgcctggcagttccctactctcgcatggggagtccccacactaccatcggcgctacggcgtttcacttctgagttcggcatggggtcaggtgggaccaccgcgctactgccgccaggcaaacaaggggtgttatgagccatattcaggtataaatgggctcgcgataatgttcagaattggttaattggttgtaacactgacccctatttgtttatttttctaaatacattcaaatatgtatccgctcatgagacaataaccctgataaatgcttcaataatattgaaaaaggaagaatatgagccatattcaacgggaaacgtcgaggccgcgattaaattccaacatggatgctgatttatatgggtataaatgggctcgcgataatgtcgggcaatcaggtgcgacaatctatcgcttgtatgggaagcccgatgcgccagagttgtttctgaaacatggcaaaggtagcgttgccaatgatgttacagatgagatggtcagactaaactggctgacggaatttatgccacttccgaccatcaagcattttatccgtactcctgatgatgcatggttactcaccactgcgatccccggaaaaacagcgttccaggtattagaagaatatcctgattcaggtgaaaatattgttgatgcgctggcagtgttcctgcgccggttgcactcgattcctgtttgtaattgtccttttaacagcgatcgcgtatttcgcctcgctcaggcgcaatcacgaatgaataacggtttggttgatgcgagtgattttgatgacgagcgtaatggctggcctgttgaacaagtctggaaagaaatgcataaacttttgccattctcaccggattcagtcgtcactcatggtgatttctcacttgataaccttatttttgacgaggggaaattaataggttgtattgatgttggacgagtcggaatcgcagaccgataccaggatcttgccatcctatggaactgcctcggtgagttttctccttcattacagaaacggctttttcaaaaatatggtattgataatcctgatatgaataaattgcagtttcatttgatgctcgatgagtttttctaagcggcgcgccatcgaatggcgcaaaacctttcgcggtatggcatgatagcgcccggaagagagtcaattcagggtggtgaatatgaaaccagtaacgttatacgatgtcgcagagtatgccggtgtctcttatcagaccgtttcccgcgtggtgaaccaggccagccacgtttctgcgaaaacgcgggaaaaagtggaagcggcgatggcggagctgaattacattcccaaccgcgtggcacaacaactggcgggcaaacagtcgttgctgattggcgttgccacctccagtctggccctgcacgcgccgtcgcaaattgtcgcggcgattaaatctcgcgccgatcaactgggtgccagcgtggtggtgtcgatggtagaacgaagcggcgtcgaagcctgtaaagcggcggtgcacaatcttctcgcgcaacgcgtcagtgggctgatcattaactatccgctggatgaccaggatgccattgctgtggaagctgcctgcactaatgttccggcgttatttcttgatgtctctgaccagacacccatcaacagtattattttctcccatgaggacggtacgcgactgggcgtggagcatctggtcgcattgggtcaccagcaaatcgcgctgttagcgggcccattaagttctgtctcggcgcgtctgcgtctggctggctggcataaatatctcactcgcaatcaaattcagccgatagcggaacgggaaggcgactggagtgccatgtccggttttcaacaaaccatgcaa | GSGSASKPDIKVGDYVKMGVYNNASILWRCVSIDNNGPLMLADKIVDTLAYDAKTNDNSNSKSHSRSYKRDDYGSNYWKDSNMRSWLNSTAAEGKVDWLCGNPPKDGYVSGVGAYNEKAGFLNAFSKSEIAAMKTVTQRSLVSHPEYNKGIVDGDANSDLLYYTDISEAVANYDSSYFETTTEKVFLLDVKQANAVWKNLKGYYVAYNNDGMAWPYWLRTPVTDCNHDMRYISSSGQVGRYAPWYSDLGVRPAFYLDSEYFVTTSGSGSQSSPYIGSAPNKQEDDYTISEPAEDANPDWNVSTEQSIQLTLGPWYSNDGKYSNPTIPVYTIQKTRSDTENMVVVVCGEGYTKSQQGKFINDVKRLWQDAMKYEPYRSYADRFNVYALCTASESTFDNGGSTFFDVIVDKYNSPVISNNLHGSQWKNHIFERCIGPEFIEKIHDAHIKKKCDPNTIPSGSEYEPYYYVHDYIAQFAMVVNTKSDFGGAYNNREYGFHYFISPSDSYRASKTFAHEFGHGLLGLGDEYSNGYLLDDKELKSLNLSSVEDPEKIKWRQLLGFRNTYTCRNAYGSKMLVSSYECIMRDTNYQFCEVCRLQGFKRMSQLVKDVDLYVATPEVKEYTGAYSKPSDFTDLETSSYYNYTYNRNDRLLSGNSKSRFNTNMNGKKIELRTVIQNISDKNARQLKFKMWIKHSDGSVATDSSGNPLQTVQTFDIPVWNDKANFWPLGALDHIKSDFNSGLKSCSLIYQIPSDAQLKSGDTVAFQVLDENGNVLADDNTETQRYTTVSIQYKFEDGSEIPNTAGGTFTVPYGTKLDLTPAKTLYDYEFIKVDGLNKPIVSDGTVVTYYYKNKNEEHTHNLTLVAAKAATCTTAGNSAYYTCDGCDKWFADATGSVEITDKTSVKIPAPGHTAGTEWKSDDTNHWHECTVAGCGVIIESTKSAHTAGEWIVDTPATATTAGTKHKECTVCHRVLETQPIPSTGTELKIIAGDNQIYNKASGSDVTITCNGDFAKFTGIKVDGSVVDSSNYTAVSGSTVLTLKASYLGTLTDGSHTITFVYTDGEANANLTVRTAGSGHIHDYGTEWKSNADNHWHECNCGDKKDEAAHSFKWVVDKEATATKKGSKHEECKICGYKRSAVEIPATGTSTAPTDTTKPNDTTKPGNTNGSEKGSHHHHHH |

**Supplementary Discussion – Validating the Gel-Based Immunoglobulin A Protease Kinetic Assay**

*Developing a Versatile Gel-Based Kinetic Assay for Immunoglobulin A Proteases*

The substrate (intact IgA1) and products (IgA1 Fc and Fab domains) of the IgAP-catalyzed reaction are easily visualized and separated on an SDS-PAGE gel (*Supp. Fig. 8a*). Monomeric IgA1 has a quaternary structure consisting of two heavy chains (HC) and two light chains (LC) (*Supp. Fig. 8b*).^1^ Under the reducing and denaturing conditions of the SDS-PAGE gel, the covalent and non-covalent bonds that hold the quaternary structure break apart and lead to two discrete bands on the gel. When monomeric IgA1 is treated with IgAP, the HC is cleaved into two products associated with the Fab or Fc.^2^ The HC region associated with the Fc is seen at ~37.5 kDa (HC fragment #1; H1) on an SDS-PAGE gel and the HC region associated with the Fab is seen at ~30 kDa (HC fragment #2; H2) (*Supp. Fig. 8b*). Although H1 and H2 have protein components that are very similar in size, they migrate differently on the gel due to the glycosylations on each fragment.^3^ H1 thus typically runs higher than H2 due to the N-linked glycosylations on the Fc.^4^

The gel-based IgAP assay capitalizes on the distinct banding patterns of the substrate and product to determine the initial rates of IgA1 cleavage at various substrate concentrations. This was done by quantifying the intensity of these bands using densitometry and tracking how this intensity changed over time to extract initial rates. It was important to validate the assay prior to making kinetic measurements of *T. ramosa* IgAP constructs to ensure that the experimental conditions followed Michaelis-Menten assumptions.

The first step of validation was to determine the functional range of the assay by fitting measurements within that range to a standard curve. Various amounts of monomeric IgA1 ranging from 10 to 100 pmol were visualized on a gel and the intensities of the HC and LC were digitally integrated for quantification (*Supp. Fig. 9a*). It was apparent that the intensity measurements at higher amounts of IgA1 deviated from the lower end of the curve (*Supp. Fig. 9b*). However, the integrated signal was linear (R^2^ > 0.99) up to and including 60 pmol IgA1 for the HC band and 40 pmol IgA1 for the LC band (*Supp. Fig. 9cd*). 60 pmol was thus set as the maximum amount of IgA1 that can be loaded onto a single gel lane while maintaining linearity with respect to the amount of IgA1 loaded.

The functional range must also account for the fact that the assay would need to be able to reliably distinguish between intensities in a reaction that progresses only 5-10%, as this is typically assumed to fulfill initial-rate conditions. Under the restrictions imposed above of a maximum loading capacity of 60 pmol IgA1 per lane, the assay would need to distinguish between intensities that fall within 3-6 pmol of cleaved IgA1. A subsequent standard curve was done where fully cleaved IgA1 was titrated from 1.25 to 60 pmol per lane (*Supp. Fig. 10a*). As was seen in *Supp. Fig. 9* for the HC, the intensity of the H1 band was linear for the entire titration range up to and including 60 pmol (*Supp. Fig. 10b*). This held true when the data were replotted with only the lower end of the standard curve (*Supp. Fig. 10c*), which was more representative of the intensity measurements that would be seen in real data under initial-rate conditions. These two standard curves suggest that the assay can detect bands reliably within the range of 1.25 to 60 pmol of IgA loaded and that these intensity measures are strongly correlated with the corresponding amount of IgA loaded per lane. Based on these results, all SDS-PAGE samples were prepared such that they contained 40 pmol of IgA regardless of the concentration of IgA in the reaction mixture to ensure that all band intensities would fall within the functional range of the assay. This meant that 20 μL reactions with substrate concentrations greater than 2 μM were diluted down to 2 μM in the SDS-PAGE sample and substrate concentrations equal to or less than 2 μM were sampled undiluted.

The second step of validation was to determine the best SDS-PAGE band to use for densitometric analysis. This is because the disappearance of the HC, appearance of the H1, and appearance of the H2 bands can all be theoretically used to measure the progress of the reaction. IgA1 was incubated with the *T. ramosa* IgAP MD and sampled over time to test the suitability of all three bands (*Supp. Fig. 11a*). Although all bands behaved as expected over the course of the reaction, the H1 band (*Supp. Fig. 11c*) had the strongest linear correlation of its band intensity with respect to time and thus was the best band to use for densitometric analysis.

It became apparent during the course of validation that an external calibration curve (*Supp. Fig. 10*) was not able to reproducibly determine the amount of product present during a reaction due to variations in band intensities between SDS-PAGE gels for the same sample. To address this, an internal standard was used to normalize the H1 band to a constant band in the same gel so that results could be compared between gels.

The light chain (LC) of IgA1 was the first internal standard tested as it is theoretically proportional to the amount of IgA1 loaded into each lane and hence would account for differences in sample preparation and gel loading. It would also be subject to the same gel workup as the H1 band and thus account for differences in gel staining, destaining, and imaging. Unfortunately, the LC was an unreliable internal control due to the diffuse nature of the LC band, which was prone to error in processing because of inaccurate background subtraction. This was made worse by the fact that the H2 band overlapped with the LC in samples with a larger proportion of IgA1 cleaved.

Most of the variation in the measured band intensities between gels seemed to be from the gel workup rather than sample preparation and loading. This was supported by the fact that bands within a gel seemed to be internally consistent without normalization (*Supp. Fig. 10* and *Supp. Fig. 11c*). This is in contrast to band intensities across gels, which were susceptible to the batch-to-batch variation of SDS-PAGE stain and destain used as well as the exposure time of the image. The SDS-PAGE staining step was particularly variable due to the differing amounts of Coomassie R-250 dye that were lost to the filter paper when the stain was made.

As an alternative approach to the LC, a batch of IgA1 was fully cleaved with the *T. ramosa* IgAP, diluted to 40 pmol per sample, and frozen in 25 μL aliquots with SDS-PAGE loading dye to serve as internal standards for each gel. The intensity of the H1 band for the internal standard would be used as the reference to determine the percentage of IgA1 cleaved at a given time point. This would obviate the need to use an external standard curve to convert changes in intensity to rate to avoid errors associated the gel workup. *Supp. Fig. 12* is a representative example of data with this new internal standard that was obtained under initial-rate conditions. The following steps were used to convert densitometry measurements into the initial rate. The H1 band intensities (*Supp. Fig. 12a*) were first normalized to the intensity of the internal standard to convert them into the percentage of IgA1 cleaved at each time point. This was done with the equation:

$$\% IgA1 Cleaved= \frac{Integrated intensity of H1 band in sample}{Integrated intensity of H1 band in internal standard}*100\%$$

The percentage of IgA1 cleaved was plotted with respect to time and used to calculate the slope from a simple linear regression (*Supp. Fig. 11b*). The unit of this slope was percentage of IgA1 cleaved per minute. Although samples from 20 μL reactions with substrate concentrations greater than 2 μM were diluted to 2 μM to fall within the functional range of the assay, the value of the slope is independent of the substrate concentration as it represents the proportion of IgA1 cleaved relative to the total amount of substrate. The initial rate is therefore simply calculated with the following equation and takes on the units of molarity per minute:

$$Initial rate \left( v \right)=Slope of linear regression*[S]$$

It was important to verify that the measurements used to calculate the linear regression were in the initial-rate region of the progress curve. This was done by assessing the correlation coefficient between the time points to ensure strong linearity (R^2^ > 0.98) and by making sure that the y-intercept of the linear regression was close to zero. Non-zero y-intercepts resulted from fitting the regression to the non-linear portion of the progress curve and run contrary to the assumptions of underlying initial rates as they imply that there is some amount of product that exists at time zero.

All *T. ramosa* IgAP enzyme constructs were surprisingly robust in fulfilling these initial-rate conditions as they were not significantly affected by product inhibition (*Supp. Fig. 13*). This made these enzymes relatively easy to characterize as experimental conditions did not need to be fully optimized for each enzyme and substrate concentration to ensure linearity. Initial-rate conditions were still valid for a reaction that progressed to nearly 50% completion (*Supp. Fig. 13ab*), which is much more than what is generally accepted for initial-rate kinetics.^5^ Product inhibition and substrate depletion only began to have an effect when the reaction proceeded to near completion (*Supp. Fig. 13cd*).

In summary, this section has presented experimental conditions where the gel-based IgAP assay does not violate the assumptions inherent to the Michaelis-Menten equation. The assay is therefore suitable for understanding the potential functions of the *T. ramosa* IgAP non-proteolytic domains in enzymatic activity. The above experiments provide data to support the validity of the assay with respect to the Michaelis-Menten assumptions as follows:

(1) The enzyme concentration (0.15-1.5 nM) was over 650 times less than the lowest concentration of substrate (1 μM) used in the reaction.
(2) The assay can detect small amounts of product from reactions that have progressed as little as ~3% (1.25 pmol in a 40 pmol sample). This falls in the rule-of-thumb range where issues of product inhibition and substrate depletion are assumed to be negligible.
(3) The integrated intensity of the H1 band has a strong linear correlation (R^2^ > 0.99) with the amount of IgA1 product loaded over a large functional range (1.25 to 60 pmol) that covers the initial-rate region as described in (2).
(4) The assay can distinguish between reactions that are in the linear and non-linear regions of the progress curve by assessing the correlation coefficient and y-intercept of the linear regression.

*Limitations to the Gel-Based Immunoglobulin A Protease Kinetic Assay*

There are five aspects of the gel-based IgAP assay that require further discussion:

Firstly, there is no practical way to quantify experimental error in the measurements or the derived kinetic parameters. Due to the way the experiment is set up and how the reaction progress is monitored, the assay only yields one experimental measurement for each time point and substrate concentration. True replication would involve setting up multiple identical assays on different days to obtain replicates of the full Michaelis-Menten curve. Although this can be done in theory, it is not practical given the cost and amount of substrate required as well as the amount of time it takes to carry out the assay. One of the reasons why the reaction was monitored over time was to provide some confidence in the derived initial rates, as linear regression can detect the level of variation, despite the lack of a real measure of error.

Secondly, if this assay is to be used for other IgAPs or Ig proteases, the functional range of the assay may not be sufficient for proteases that are more sensitive to product inhibition. Qualitative results for the *Streptococcus pneumoniae* and *Haemophilus influenzae* IgA1Ps suggest that they bind much tighter to IgA1 as the inactive mutants of the protein can form complexes with IgA1 that can be isolated and characterized separately (data not shown). This is the basis for how the structure of the *S. pneumoniae* IgA1P-IgA1 complex was able to be determined by cryo-electron microscopy.^6^ It is thus reasonable to assume that the enzymes also have some affinity for the IgA1 cleavage products based on their structural similarities to the intact IgA1. If this is the case, the assay may need to reliably detect product from reactions that progress less than 5%, which begins to fall outside of the tested range of the assay.

Preliminary semi-quantitative testing with the gel-based assay suggests that the *S. pneumoniae* and *H. influenzae* IgA1Ps are also significantly faster than the *T. ramosa* IgAP (data not shown). If their k_cat_/K_M_ values are much larger than the *T. ramosa* IgAP, the assay may need to use substrate concentrations much lower than 1 μM to fully define the curve, which may fall below the lower limit of detection for a Coomassie R-250-stained SDS-PAGE gel. Initial testing using silver stain instead of Coomassie R-250 suggested that silver staining was not suitable for kinetic analysis (data not shown). This was because the gels were easily overdeveloped in a matter of seconds and thus it was not a reproducible way to ensure that the gel bands remained in the functional range of the assay.

In addition, the *H. influenzae* IgA1P cleaves IgA1 in the middle of the hinge (*Fig. 1*) and splits the HC into two nearly-equivalent fragments. This means that H1 and H2 are no longer separable on an SDS-PAGE gel and cannot be quantified individually. This can be alleviated by pre-treating IgA1 with neuraminidase to remove negatively charged sialic acids from the IgA1 glycosylations, which changes the migration behavior of H1 and H2 such that they can be separated on a gel.^3^ The effect of removing sialic acid from the substrate may impact its ability to be cleaved and hence be semi-artifactual, however, initial studies on the *Neisseria gonorrhoeae* and *N. meningitidis* IgA1Ps suggest that desialyation does not impact activity.^3^ Regardless, this may limit its use towards characterizing certain IgAPs and is an additional point of consideration for the generalizability of the assay.

Finally, this gel-based kinetic assay assumes that the rate of cleavage and the enzyme’s specificity are identical for a fully intact IgA1 molecule (two Fab domains) and a half-cleaved IgA1 molecule (only one Fab). This is important to note as there is only one active site per enzyme and, as such, the complete cleavage of IgA into one Fc and two Fab domains must occur in two sequential reactions. Due to the reducing and denaturing conditions of the SDS-PAGE gel, it is impossible to distinguish whether the product bands (H1 and H2) come from a half-cleaved or fully cleaved IgA1 molecule. However, the fact that the rate remains linear up to ~50% of the reaction (*Suppl. Fig. 13ab*) suggests that there may not be significant kinetic differences between the two substrates as the rate does not appreciably change under conditions where a non-negligible amount of half-cleaved IgA1 is present in solution.

*Kinetic Comparisons of Thomasclavelia ramosa Immunoglobulin A Protease Constructs*

The Michaelis-Menten curves and parameters for various *T. ramosa* IgAP domain constructs are shown in *Fig. 4* and *Supp. Table 2*, respectively.

k_cat_/K_M_ represents the effective rate of substrate capture into a catalytically competent state.^7^ It is mainly impacted by the rate-determining step, which varies from enzyme to enzyme and can be the step of substrate binding, chemical catalysis, or product release.^8^ It is hypothesized that the rate-determining step for the *T. ramosa* IgAP is substrate binding. This is because the conformationally heterogeneous active site seen in the crystal structure suggests that conformational changes in and distal to the active site are necessary for catalysis, similar to the known mechanisms of selectivity for the S6 and M26 IgAPs.^6,9,10^ These conformational changes require large rearrangements of the substrate and enzyme which may lead to a greater entropic penalty that increases the energetic barrier of the substrate-binding transition states.

Assuming the rate-determining step does not change when the enzyme is minimized, which may hold true due to the modularity of the enzyme, the fact that k_cat_/K_M_ increases as the enzyme is truncated suggests that the non-proteolytic domains do not contribute to substrate binding. If the NTD and/or C-terminal tail significantly contributed to substrate binding, k_cat_/K_M_ would be expected to decrease as the enzyme was truncated. The changes in catalytic efficiency are therefore postulated to be due to steric effects from the non-proteolytic domains, which lowers the frequency of forming a catalytically competent complex with the protease. If there were any energetic benefits involved in substrate binding from the non-proteolytic domains, the experimental results indicate that these may be outweighed by this steric factor. The differences in k_cat_/K_M_ between MD and MD+CTD1 also support this steric hypothesis, assuming that they are statistically significant when true experimental error is measured.

k_cat_ represents the rate of regenerating free enzyme after it is captured into the enzyme-substrate complex.^7^ Because k_cat_ is calculated from a fit of the Michaelis-Menten equation at infinite substrate concentration, under these conditions, the time it takes to form the enzyme-substrate complex is negligible and thus the substrate-binding step cannot be measured using k_cat_.^7^ The main bottlenecks that contribute to k_cat_ are thus chemical catalysis and product release.^7^ There is no experimental evidence to suggest that one step contributes more than the other, however as the same trend is seen in k_cat_ as k_cat_/K_M_, one can postulate that the major contributor to k_cat_ is product release for the same reasons of steric interference preventing the product from leaving the enzyme-product complex. Because a large increase in k_cat_ was seen when the NTD was removed, the NTD may inhibit product release as it may have some affinity to the cleaved IgA products.

**References**

1.	Woof, J. M. & Kerr, M. A. The function of immunoglobulin A in immunity. *J Pathol* **208**, 270–282 (2006).

2.	Mistry, D. & Stockley, R. A. IgA1 protease. *Int J Biochem Cell Biol* **38**, 1244–1248 (2006).

3.	Gilbert, J. V, Plaut, A. G., Longmaid, B. & Lamm, M. E. Inhibition of Microbial IgA Proteases by Human Secretory IgA and Serum. *Mol Immunol* **20**, 1039–1049 (1983).

4.	Yoo, E. M. & Morrison, S. L. IgA: An immune glycoprotein. *Clinical Immunology* **116**, 3–10 (2005).

5.	Srinivasan, B. A guide to the Michaelis–Menten equation: steady state and beyond. *FEBS Journal* **289**, 6086–6098 (2022).

6.	Wang, Z. *et al.* Mechanism and inhibition of Streptococcus pneumoniae IgA1 protease. *Nat Commun* **11**, (2020).

7.	Northrop, D. B. On the Meaning of Km and V/K in Enzyme Kinetics. *J Chem Educ* **75**, 1153 (1998).

8.	Park, C. Visual Interpretation of the Meaning of kcat/Km in Enzyme Kinetics. *J Chem Educ* **99**, 2556–2562 (2022).

9.	Redzic, J. S. *et al.* A substrate-induced gating mechanism is conserved among Gram-positive IgA1 metalloproteases. *Commun Biol* **5**, (2022).

10.	Johnson, T. A., Qiu, J., Plaut, A. G. & Holyoak, T. Active site gating regulates substrate selectivity in a chymotrypsin-like serine protease. The structure of Haemophilus influenzae IgA1 protease. *J Mol Biol* **389**, 559–574 (2009).
